# Amygdalostriatal transition zone neurons encode sustained cue responses to guide defensive behaviors

**DOI:** 10.1101/2022.10.28.514263

**Authors:** Fergil Mills, Christopher R. Lee, James R. Howe, Hao Li, Maria N. Keisler, Shan Shao, Felix H. Taschbach, Mackenzie E. Lemieux, Faith Aloboudi, Jesse White, May G. Chan, Matilde Borio, Laurel R. Keyes, Hannah S. Chen, Fabiha Bushra, Gates P. Schneider, Dani P. Lemmon, Kyung J. Lee, Alexa L. Gross, Kanha Batra, Reesha R. Patel, Meenakshi M. Asokan, Jeremy Delahanty, Christian Cazares, Christopher R. Heyman, Nicholas B. Poll, Liezl Maree, Romy Wichmann, Talmo D. Pereira, Marcus K. Benna, Cory M. Root, Kay M. Tye

## Abstract

To ensure survival, the brain must rapidly identify threats and maintain defensive behaviors for as long as danger is present. The amygdala has been studied as a key site for fear responses, but responses to threat cues in amygdala neurons are largely transient and shorter in duration than defensive responses observed. Here, we present the amygdalostriatal transition zone (ASt) as a missing piece of the circuits mediating fear responses. Using single-nucleus RNA sequencing (snRNA-seq), we demonstrate that the ASt is genetically distinct from adjacent striatal and amygdalar structures. *In vivo* electrophysiology and calcium imaging reveal that ASt neurons have robust, sustained responses to shock-predicting cues. Further, photostimulation of the ASt is sufficient to drive freezing and avoidance behaviors, and optogenetic inhibition experiments show that *Drd2+* ASt neurons are necessary for cue-conditioned fear responses. Our findings establish the ASt as a previously unappreciated yet critical structure for encoding learned associations and directing defensive behaviors.

**HIGHLIGHTS:**

- ASt neurons have a unique genetic identity
- ASt neurons encode sustained responses to conditioned shock cues
- Activation of ASt neurons is sufficient to drive freezing and avoidance
- *Drd2+* ASt neurons are necessary for expression of cued fear responses

## INTRODUCTION

When danger is present, the brain must recognize threat-predictive cues and orchestrate an evolutionarily conserved “fear response” that includes selecting and maintaining defensive behaviors.^1–9^ These responses are shaped by associative learning,^4,10–16^ which enables the appropriate responses to environmental stimuli based on the rewards or punishments they predict.

For decades, the amygdala has been studied as the central site for fear learning and defensive behaviors.^3–6,11–13,17–22^ In the “canonical” model of fear learning, neurons in the lateral amygdala (LA) and basolateral amygdala (BLA) integrate cue and outcome information,^23–25^ synaptic plasticity strengthens inputs carrying threat-predictive stimuli,^25–30^ and outputs from the central nucleus of the amygdala (CeA) mediate defensive responses such as freezing and flight.^3,5, 6,9,31,32^ However, there is a considerable discrepancy in timescales between activity observed in the amygdala complex and the behaviors elicited by threats. Neurons in the amygdala show increased responses to a conditioned stimulus (CS) following fear learning, but these responses are often only transient, lasting less than a second from the onset of the stimulus.^20,33–36^ This does not match the duration of behaviors that can be elicited by these stimuli, which include defensive responses expressed for the full duration of threat presentations. Mediating these behaviors would ideally require a neural substrate that is connected to circuits for learning, exhibits rapid and sustained responses to threats, and is upstream of circuits for directing behavioral responses.

The amygdalostriatal transition zone (ASt) is uniquely situated to play a role in these processes. Like the LA, the ASt receives converging inputs from thalamic and cortical pathways that are known to relay sensory information critical for learned associations. Notably, the ASt receives robust input from thalamic projections carrying threat information^25,37^ and a direct projection from the LA,^38,39^ both of which preferentially target the ASt compared with the adjacent tail of striatum (TS). ASt neurons also respond to auditory, visual, and somatosensory stimuli relevant to fear learning with faster latencies than amygdala neurons.^40,41^ However, the outputs of the ASt diverge from canonical amygdala pathways and are integrated with striatal circuits known to control action selection.^42–47^ ASt neurons project to the substantia nigra pars lateralis (SNL)^38,48^ and external globus pallidus (GPe), known targets of the striatal “direct pathway” neurons expressing the dopamine D1 receptor (*Drd1a*) and “indirect pathway” neurons expressing the dopamine D2 receptor (*Drd2*).^42–46,49^ Thus, the ASt is at a crucial intersection of systems in the brain, bridging amygdala circuits for fear learning with basal ganglia circuits that direct behavior.

Here, using a systematic comparison spanning systems, circuit, and molecular levels, we show that the ASt is a genetically distinct region that encodes sustained responses to threats across behavioral timescales. Our findings identify the ASt as a previously unappreciated missing piece of the circuitry for associative learning and directing defensive responses to threats.

## RESULTS

### The ASt has a distinct transcriptomic identity from the TS or CeA

We first investigated whether the ASt was genetically distinct from adjacent amygdala and striatal structures. We performed single-nucleus RNA sequencing (snRNA-seq)^50^ on histologically verified tissue samples from the ASt, the CeA, the TS, and the dorsal striatum (DS) (97,434 nuclei; Figures 1A and S1). We visualized clustering of sequenced nuclei by gene expression via uniform manifold approximation projection (UMAP) and identified cluster cell types based on known gene markers.^51,52^ The majority of neurons from all sequenced regions fell into two major clusters of GABAergic medium spiny neurons (MSNs) expressing *Drd1a* or *Drd2* (Figures 1B–1D). Intriguingly, within these clusters, cells from different regions were ordered in high-dimensional expression space according to their anatomical proximity in the brain (DS → TS → ASt → CeA). Most strikingly, ASt nuclei formed visibly distinct subclusters within the *Drd1a+* and *Drd2+* clusters (Figure 1B). We confirmed there were no region-specific differences in expression of markers for nuclear quality^53^ (Figures S1J and S1K) or excessive variation due to batch effects (Figures S1L–S1N), indicating the clustering observed reflected real biological differences between the ASt and other regions.

**Figure 1.**
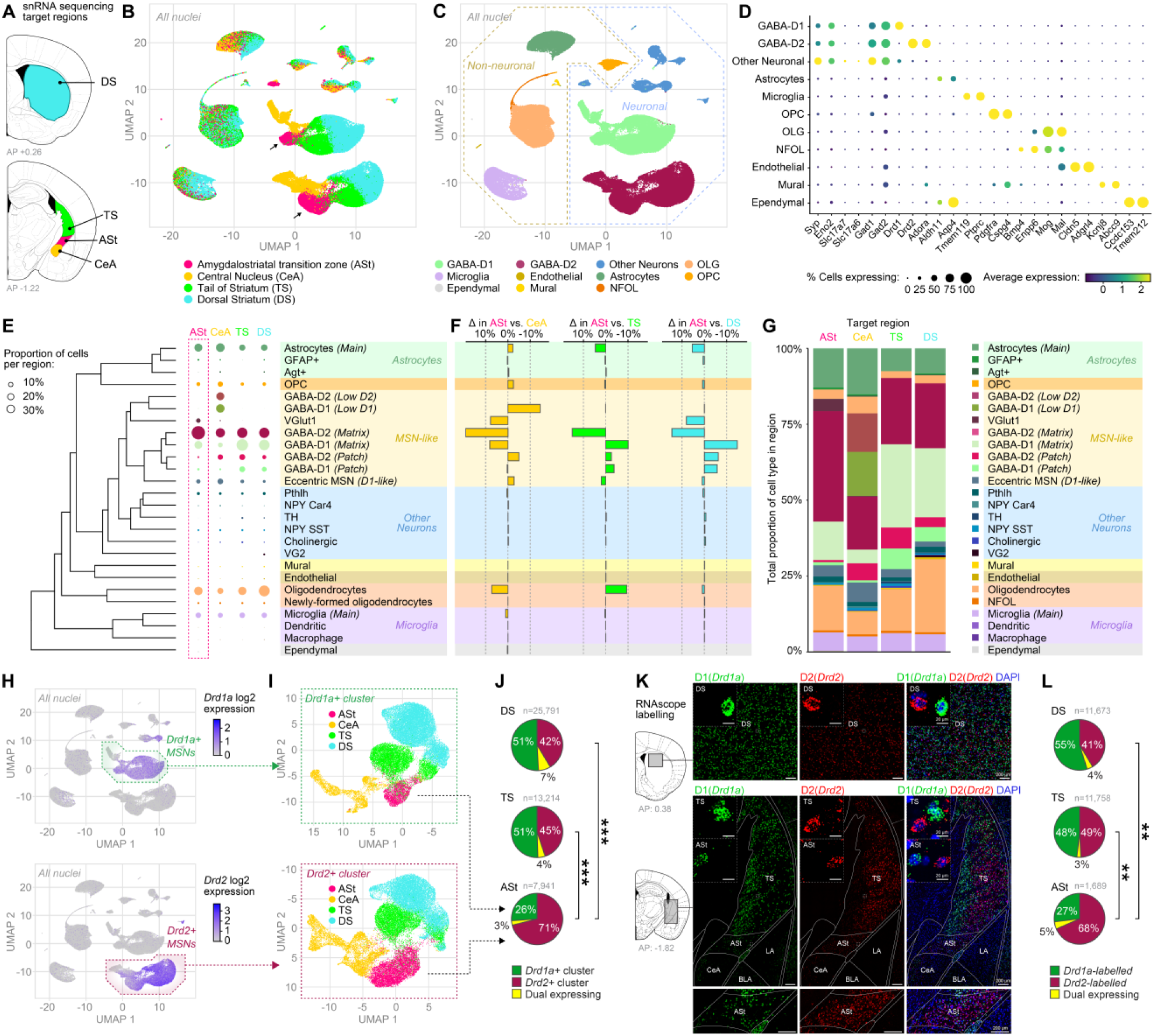
The ASt is genetically distinct from the amygdala and striatum. **(A)** Target regions for single-nucleus RNA sequencing (snRNA-seq). **(B)** Two-dimensional uniform manifold approximation projection (UMAP) of all sequenced nuclei passing quality filters (*n* = 97,434 nuclei, see also Figure S1), colored by region of origin. Arrows indicate distinct ASt subclusters within the GABAergic *Drd1a+* and *Drd2*+ clusters. **(C)** Two-dimensional UMAP, colored by broad cellular identity assigned by graph-based clustering of neuronal and non-neuronal cells (GABA-D1, GABAergic neurons expressing Drd1a cluster; GABA-D2, GABAergic neurons expressing Drd2 cluster; OLGs, oligodendrocytes; NFOL, newly formed oligodendrocytes; OPCs, oligodendrocyte precursor cells). **(D)** Cell-type-specific expression of canonical marker genes indicating broad cellular identity in the brain. Dot size is proportional to the percentage of nuclei expressing the marker, with the color scale representing normalized expression level. **(E)** Dendrogram of cell-type classification and proportion of cells of each class in the ASt and other target regions. **(F)** Difference in overall proportion of each cell type in the ASt compared with the CeA, TS, and DS. **(G)** Total proportion of cells of each identified type in each target region. **(H)** All nuclei, colored by expression levels of *Drd1a* (top) or *Drd2* (bottom), with cells identified as part of the major *Drd1a+* or *Drd2*+ clusters highlighted. **(I)** *Drd1a+* cluster (top) or *Drd2*+ cluster (bottom) neurons, with individual nuclei colored by region of origin. **(J)** Relative proportion of nuclei classified in *Drd1a+* or *Drd2*+ clusters in each striatal target region. Nuclei in the *Drd1a+* cluster that also expressed *Drd2*, or in the *Drd2*+ cluster that expressed *Drd1a*, were classified as “dual expressing.” ASt *Drd1a*+*:Drd2+* proportions were significantly different from those in the TS or DS (χ^2^ tests, Holm-corrected: ASt vs. TS, χ^2^(1) = 1,329, *p* = 1.4 × 10^−290^, V = 0.26; ASt vs. DS, χ^2^(1) = 1,866, *p* = 3.6 × 10^−407^, V = 0.24). Due to the high N, the TS vs. DS also differed significantly (TS vs. DS, χ^2^(1) = 15.9, *p* = 6.7 × 10^−5^, V = 0.02) but had an effect size an order of magnitude smaller than either ASt comparison. ASt: 7,941; TS: 13,214; DS: 25,791 nuclei. \*\*\**p* < 0.001. **(K)** Representative images of *in situ* RNAscope labeling of *Drd1a* RNA (green) and *Drd2* RNA (red) in striatal target regions. Bottom panel shows ASt with increased contrast to visualize labeled neurons. **(L)** Relative proportion of cells in each target region positively labeled for *Drd1a* RNA, *Drd2* RNA, or both (dual expressing). *Drd1a+*:*Drd2*+ proportions compared by paired *t* tests on per-animal log-ratios, Holm-corrected for 3 pairwise comparisons: ASt vs. TS, t(6) = 5.51, *p* = 0.0039; ASt vs. DS, t(7) = 5.17, *p* = 0.0039; TS vs. DS, t(6) = 2.22, *p* = 0.068. DS: *n* = 8 mice, 11,673 cells; TS: *n* = 7 mice, 11,758 cells; ASt: *n* = 8 mice, 1,689 cells. \*\**p* < 0.01.

We then analyzed relative proportions of cell types in each region and found the cell-type composition of the ASt was distinct from the CeA, TS, and DS (Figures 1E–1G). The DS and TS had roughly equal proportions of the two major MSN types, consistent with previous reports,^54,55^ but the ASt was markedly enriched for *Drd2+* MSNs (71% *Drd2+* vs. 26% *Drd1a+*; Figures 1H–1J). To validate these findings with an independent method, we used RNAscope labeling to examine *Drd1a* and *Drd2* expression *in situ*, which confirmed the enrichment of *Drd2+* neurons in the ASt relative to other regions of the striatum (68% *Drd2+*/27% *Drd1a+*; Figures 1K and 1L). Notably, while CeA neurons fell into the *Drd1a+* and *Drd2+* MSN clusters based on overall transcriptomic similarity, many of these neurons did not actually express the *Drd1a* or *Drd2* genes themselves (Figures 1B and 1H). This indicates the existence of “D1-like” and “D2-like” transcriptomic superclusters that include neurons lacking expression of those defining receptor genes. This pattern was consistent with our RNAscope data, which showed few *Drd1a+* or *Drd2+* labeled cells in the CeA (Figure 1K).

ASt neurons within both the *Drd1a+* and *Drd2+* clusters had large numbers of differentially expressed genes (DEGs) compared with neurons from other regions (Figures S2 and S3). Among these DEGs were synaptic adhesion molecules and axon guidance factors, which are likely candidates responsible for establishing the unique connectivity of the ASt. We examined two candidates, *Cdh6* and *Slit2*, and used RNAscope labeling to confirm they were enriched in the ASt (Figure S4). For both genes, expression was highest in the ASt and decreasing in a dorsoventral gradient through the TS, reaching its lowest levels in the DS. There was also a sharp decrease in expression from the ASt to the CeA, adding another molecular feature that distinguishes these structures. Intriguingly, *Cdh6* and *Slit2* expression was as high in the LA as in the ASt. Both genes play roles in axon-target matching and synapse formation,^56,57^ and their shared enrichment in the ASt and LA closely mirrors the pattern of thalamocortical inputs to these structures, which collateralize to both the LA and the ASt.^25,37^

Taken together, these results demonstrate major diversity of gene expression within *Drd1a+* and *Drd2+* neurons from different striatal subregions and provide the first characterization of a unique genetic identity of the ASt relative to the CeA, TS, and DS.

### ASt neurons encode sustained conditioned responses to threat cues

We then examined whether ASt neurons encoded conditioned responses to threat and reward-predicting cues. We recorded the activity of ASt neurons in freely moving mice using chronically implanted probes for *in vivo* electrophysiology (Figures 2A–2C and S5A–S5C) during a two-tone discrimination task, in which distinct conditioned stimuli (20 s auditory cues, 3.5 or 20 kHz, counterbalanced) predicted either foot shock (“CS-shock”) or the delivery of reward (“CS-reward”) (Figure 2C). Mice were trained on this task until they showed robust discriminative behavioral responses to each CS type (Figures S5D–S5G), allowing us to examine their neural activity for encoding of the learned associations and conditioned behaviors. Our probes targeting the ASt also recorded neuronal responses from TS and CeA (Figure 2B), allowing direct comparison of conditioned responses across structures.

**Figure 2.**
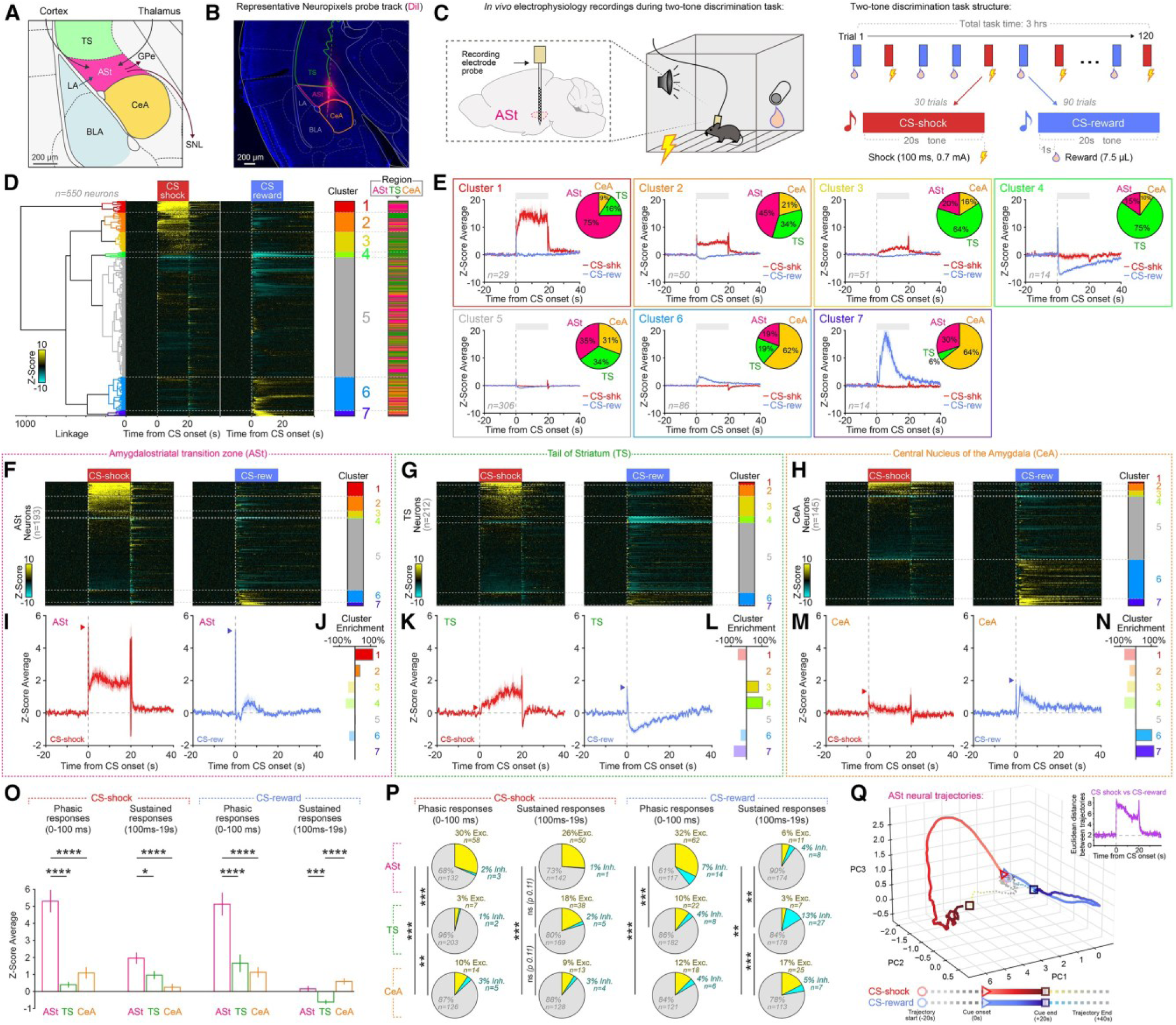
ASt neurons exhibit sustained responses to shock-predictive cues. **(A)** The amygdalostriatal transition zone (ASt), surrounding regions, and major ASt inputs and outputs. **(B)** Representative Neuropixels probe tracks targeting the ASt, tail of striatum (TS), and central nucleus of the amygdala (CeA). **(C)** Two-tone discrimination task design for *in vivo* electrophysiology recordings. **(D)** Hierarchical clustering of all recorded neurons based on responses to CS-shock and CS-reward. Dendrogram (left) shows clustering of neurons based on similarity of cue responses. Heatmap rows (middle) show trial-averaged *Z* score responses for individual neurons. Colored bars (right) indicate cluster identity and region of origin for each neuron. **(E)** Mean *Z* score responses to CS-shock and CS-reward for all neurons in each functional cluster. Pie charts show the normalized proportion of neurons from ASt, TS, and CeA in each cluster. ASt neurons were the largest proportion of clusters 1 and 2, which exhibited sustained responses to CS-shock. **(F–H)** Heatmaps of *Z* score responses to CS-shock and CS-reward in ASt (F), TS (G), and CeA (H). Colored bars indicate cluster identity. **(I, K, and M)** Mean *Z* scored responses to CS-shock and CS-reward in the ASt (I), TS (K), and CeA (M). Arrowheads indicate the mean peak *Z* score during the phasic window (0–100 ms). **(J, L, and N)** Relative enrichment of functional clusters in ASt (J), TS (L), and CeA (N). **(O)** ASt neurons show greater phasic and sustained responses to CS-shock than neurons in the TS or CeA (one-way ANOVA with region as factor, CS-shock phasic, *F*(2, 547) = 38.46, *p* < 0.0001; CS-shock sustained, *F*(2, 547) = 9.41, *p* < 0.0001; CS-reward phasic, *F*(2, 547) = 15.57, *p* < 0.0001; CS-reward sustained, *F*(2, 547) = 16.38, *p* < 0.0001, followed by Tukey’s multiple comparison test; \**p* < 0.05, \*\**p* < 0.01, \*\*\**p* < 0.001, \*\*\*\**p* < 0.0001). **(P)** Proportions of neurons in the ASt, TS, and CeA that were significantly excited or inhibited by CS-shock or CS-reward in the phasic and sustained time periods (\*\**p* < 0.01, \*\*\**p* < 0.001, chi-squared test). **(Q)** ASt neuron population trajectories in principal-component space show diverging responses to CS-shock and CS-reward. Inset shows the Euclidean distance between trajectories of CS-shock and CS-reward responses in ASt. ASt: *n* = 15 mice, 193 neurons; TS: *n* = 7 mice, 212 neurons; CeA: *n* = 8 mice, 145 neurons. Data shown as mean ± SEM.

We began by using hierarchical clustering to group all recorded neurons based on the similarity of their responses to CS-shock and CS-reward (Figure 2D). Then, we examined the proportion of neurons in each cluster that were from the ASt, TS, or CeA (Figure 2E, pie charts). There was considerable heterogeneity of responses within regions, and neurons from the ASt, TS, and CeA were found in every one of the activity-defined clusters observed. Remarkably, however, ASt neurons were the dominant representative in two clusters (clusters 1 and 2) that showed robust and sustained responses to CS-shock lasting the full 20 s of the cue. By contrast, TS neurons were overrepresented in a cluster showing gradually increasing CS-shock responses (cluster 3) and a cluster inhibited during CS-reward (cluster 4). CeA neurons dominated clusters showing excitatory responses to CS-reward (clusters 6 and 7).

We then examined each region’s neural responses to the two CS types (Figures 2F–2N). ASt neurons showed conditioned responses to both CS-shock and CS-reward at the population level (Figures 2F–2J), a previously unknown role for this structure, and were enriched for the sustained response clusters. High-temporal-resolution visualization of these responses (Figure S6) showed a rapid phasic component within the first 100 ms followed by a sustained plateau, so we quantified responses to each CS in these “phasic” (0–100 ms) and “sustained” (100 ms–19 s) windows (Figures 2O and 2P). The ASt had robust phasic responses to both CS types and sustained responses to CS-shock of significantly greater magnitude than the TS or CeA (Figures 2O and S6). A greater proportion of ASt neurons also showed phasic responses to each CS and sustained responses to CS-shock (Figure 2P). TS neurons showed distinct responses from the ASt, with almost no rapid phasic responses to CS-shock and lower-magnitude responses that appeared gradually during the sustained window. The CeA had phasic responses to both CS types but less sustained responses. Overall, each region examined had a unique pattern of responses to each CS, and the magnitude of phasic and sustained responses to CS-shock was a key feature differentiating the ASt from both the TS and CeA. We also confirmed the ASt responses observed were not simply sensory responses, as they were not seen in control mice where the same CS cues were explicitly unpaired from unconditioned stimulus (US) outcomes (Figure S7).

We next compared neural trajectories^58,59^ of ASt responses to CS-shock and CS-reward and found that they diverged in orthogonal directions from cue onset both at the population level (Figure 2Q) and in individual trial pairs (Figures 2 Q, S8, and S9). Robust sustained responses were only seen in response to CS-shock and were not present during CS-reward cues prior to US-reward collection (Figure S9). This indicated that though subsets of ASt neurons responded to both CS-shock and CS-reward, the ASt encoded these two stimulus types distinctly, rather than a general salience signal.

We then examined in more detail the relationship between ASt neuron activity and behavioral responses to the CS-shock. We quantified total defensive behaviors (freezing and darting)^2,7^ to CS-shock for individual mice and used hierarchical clustering to separate them into “high” and “low” defensive performance groups (Figures S10A–S10D). In the high defensive group, ASt neuron phasic and sustained responses were over 3-fold greater than in the low defensive group (Figures S10E–S10H). We then analyzed differences in ASt neuron activity between different quartiles of trials with high or low defensive behaviors (Figures S10I–S10K). Trials with the most defensive behavior (top quartile) showed 26% greater phasic and 60% greater sustained ASt responses than trials with the least defensive behavior (Figures S10L and S10M). We also performed similar analyses of reward responses but found there was no relationship between ASt neuron activity and reward behavior (Figures S10N–S10U). These results revealed that ASt neuron activity in response to a shock-predicting CS was correlated with the amount of defensive behaviors elicited by that cue.

### ASt neuron activity is sufficient to drive freezing and avoidance behavior but does not strongly encode motor actions

We next examined whether activation of ASt neurons was sufficient to drive behavioral responses. We targeted channelrhodopsin-2 (ChR2) to GABAergic ASt neurons using a Cre-conditional viral vector in *VGAT*-Cre mice (Figures 3A and S11), preventing any confounding ChR2 expression in excitatory neurons in the adjacent LA.^60,61^ Photostimulation of ASt neurons (473 nm, 10 mW, 20 Hz) produced a striking increase in freezing during “laser ON” epochs of an open field test (Figure 3B). Additionally, in a real-time place preference task, ChR2 group mice showed avoidance of the arena side paired with ASt stimulation (Figure 3C). These data indicated that activation of ASt neurons was innately aversive and sufficient to drive freezing and avoidance behaviors, consistent with a role for the ASt in encoding aversive stimuli to direct behavioral responses.

**Figure 3.**
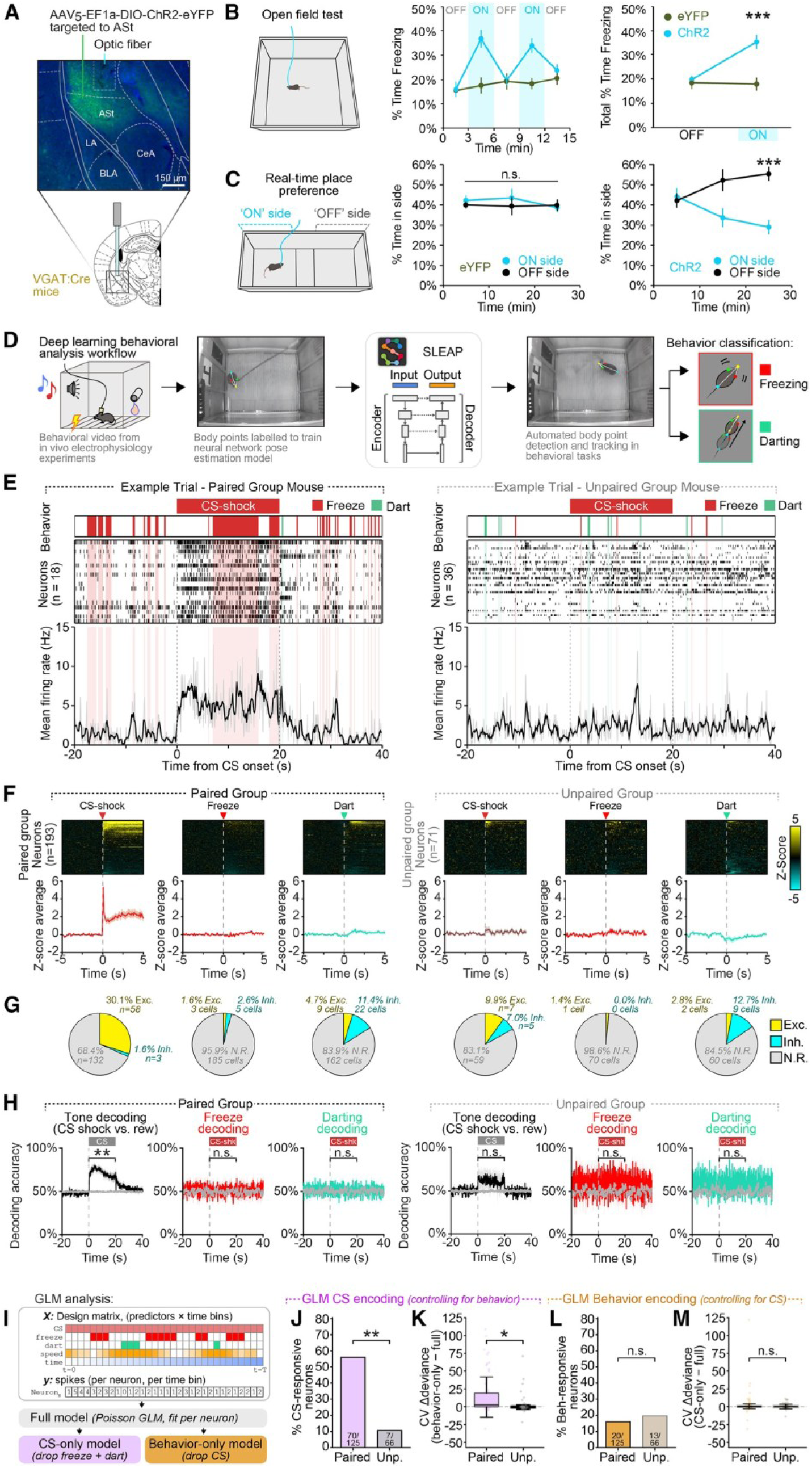
ASt neuron activity drives freezing and avoidance but encodes conditioned cues rather than subsecond motor output. **(A)** Targeting strategy and representative image of ChR2 expression in ASt neurons. **(B)** Optogenetic activation of ASt neurons during light *ON* epochs of an open field task increased freezing in ChR2 group mice (two-way ANOVA, group × laser interaction: *F(1,32)* = 10.48, *p* = 0.0028; Bonferroni post hoc test: \*\*\**p* < 0.001). **(C)** Activation of ASt neurons was sufficient to drive real-time place avoidance of the stimulus-paired “on side” in ChR2 group mice (two-way RM ANOVA, ChR2: effect of side F(1,14) = 8.187, *p* = 0.0126; eYFP: effect of side F(1,18) = 0.2911, *p* = 0.5961; Bonferroni post hoc analysis \*\*\**p* < 0.001). ChR2: *n* = 8 mice, eYFP: *n* = 10 mice. **(D)** Workflow for classification of defensive behaviors using SLEAP. **(E)** Representative CS-shock trial from individual mice in the paired and unpaired groups during the discrimination task, overlaid with defensive behavior. In each example trial, neurons shown are all ASt neurons recorded from that mouse during that trial. **(F)** Average *Z* score responses of each ASt neuron aligned to all onsets of CS-shock, freezing, and darting during the discrimination task in paired and unpaired group mice. **(G)** Proportion of ASt neurons that were significantly excited, inhibited, or non-responsive to the onset of CS-shock, freezing, and darting in paired and unpaired group mice. **(H)** Logistic regression decoder accuracy over time relative to chance level (gray line) for decoding tone identity, freezing, or darting in paired and unpaired group mice. Inset statistics indicate a Mann-Whitney U test of decoding accuracy during the CS window vs. chance (\*\**p* < 0.01). Line plots and trace data are shown as mean ± SEM. **(I)** Schematic of generalized linear model (GLM) analysis. Per-neuron Poisson GLMs were fit with CS, freezing, darting, locomotion speed, and within-trial time as predictors. **(J)** Fraction of CS-responsive neurons after controlling for freezing, darting, and locomotion speed. Using GEE Poisson models with AR(1) working correlation and trial-level permutation tests (1,000 iterations, false discovery rate [FDR]-corrected), a significantly larger proportion of neurons in paired mice showed CS encoding (70/125; 56.0%) compared with unpaired controls (7/66; 10.6%; Fisher’s exact test, \*\*\**p* < 0.001). **(K)** Cross-validated prediction improvement from CS identity. Positive CV-ΔDeviance values indicate that adding CS type to the model improved out-of-sample spike count prediction beyond behavioral covariates alone. This improvement was significantly greater in paired than in unpaired mice (animal-level Mann-Whitney U, *p* = 0.029). **(L)** Fraction of behavior-responsive neurons after controlling for CS identity. The proportion of neurons whose firing was significantly predicted by freezing, darting, and locomotion speed was low and did not differ between groups (paired 20/125; unpaired 13/66; Fisher’s exact test, *p* = 0.549). **(M)** Cross-validated prediction improvement from behavioral covariates. Adding freezing and darting to the model did not meaningfully improve out-of-sample prediction in either group, and the effect did not differ between paired and unpaired mice (animal-level Mann-Whitney U, *p* = 0.953).

We then sought to examine any possible contribution of motor output to ASt activity observed and determine whether ASt activity during CS-shock was encoding the conditioned cue or specific defensive behaviors during the cue. To distinguish between these possibilities, we obtained pose estimation data using Social LEAP Estimates Animal Poses (SLEAP)^62^ and classified individual bouts of freezing and darting behavior during the discrimination task (Figure 3D). Examination of individual trials (Figures 3E and S11D) showed that ASt neuron activity closely tracked the CS-shock in paired group mice. Bouts of defensive behaviors increased during the CS-shock but were not closely aligned with subsecond changes in ASt neuron firing. No net change in average ASt activity was observed at motor bout onsets, and few individual ASt neurons showed excitatory motor-aligned responses (Figures 3F and 3G). Logistic regression^59^ showed that CS identity could be decoded from ASt activity above chance in paired but not unpaired mice, while freezing and darting could not be decoded in either group (Figure 3H).

We then applied generalized linear model (GLM) analysis to our electrophysiology data, fitting nested models with CS type, behavioral state (freezing, darting), locomotion speed, and within-trial time as predictors for each neuron (Figure 3 I; see also Methods). In paired group mice, 56% of ASt neurons showed significant CS encoding after controlling for freezing, darting, and locomotion speed, compared with only 10% in unpaired controls (Figure 3J). Cross-validated model comparison further showed that including CS identity improved out-of-sample prediction of spike counts beyond behavioral covariates alone, and this improvement was significantly greater in paired than in unpaired mice (Figure 3K). By contrast, fewer than 20% of neurons in either group showed significant encoding of freezing or darting after controlling for CS identity (Figure 3L), and adding behavioral covariates did not improve prediction in either group (Figure 3M). Together, these data indicated that ASt activity represented conditioned cues and not co-occurring motor activity.

In the DS, *Drd1a+* and *Drd2+* MSNs define the “direct” and “indirect” pathways, which play complementary roles in action selection and motor output.^43–46^ Given the established importance of these two cell types and the enrichment of *Drd2+* MSNs in the ASt, we asked whether these populations had dissociable roles in mediating behavioral responses in the ASt. We first examined the behavioral effects of activating these two populations separately, targeting a viral vector conditionally expressing ChR2 to the ASt of *Drd1a-Cre* or *Drd2-Cre* mice (Figures 4, S12A, and S12B). Photostimulation of *Drd2+* ASt neurons drove robust freezing (Figure 4E) and real-time avoidance of a side paired with stimulation (Figures 4G, S12D, and S12F), while activation of *Drd1a+* ASt neurons had no effect on either behavior (Figures 4D, 4F, S12C, and S12E). To test whether the freezing behavior observed could be due to “motor arrest,” or disruption of the animal’s ability to make controlled movements, we also tested mice on an accelerating rotarod but found that activation of *Drd1a+* or *Drd2+* neurons did not affect performance on this task (Figures 4H and 4I). Activation of *Drd1a+* and *Drd2+* ASt neurons did not have a significant effect on time spent in open arms of an elevated plus maze (Figures S12G and S12H). Together, these results showed that activation of the *Drd2+* population of ASt neurons was sufficient to drive freezing and avoidance behavior that was not attributable to motor disruption.

**Figure 4.**
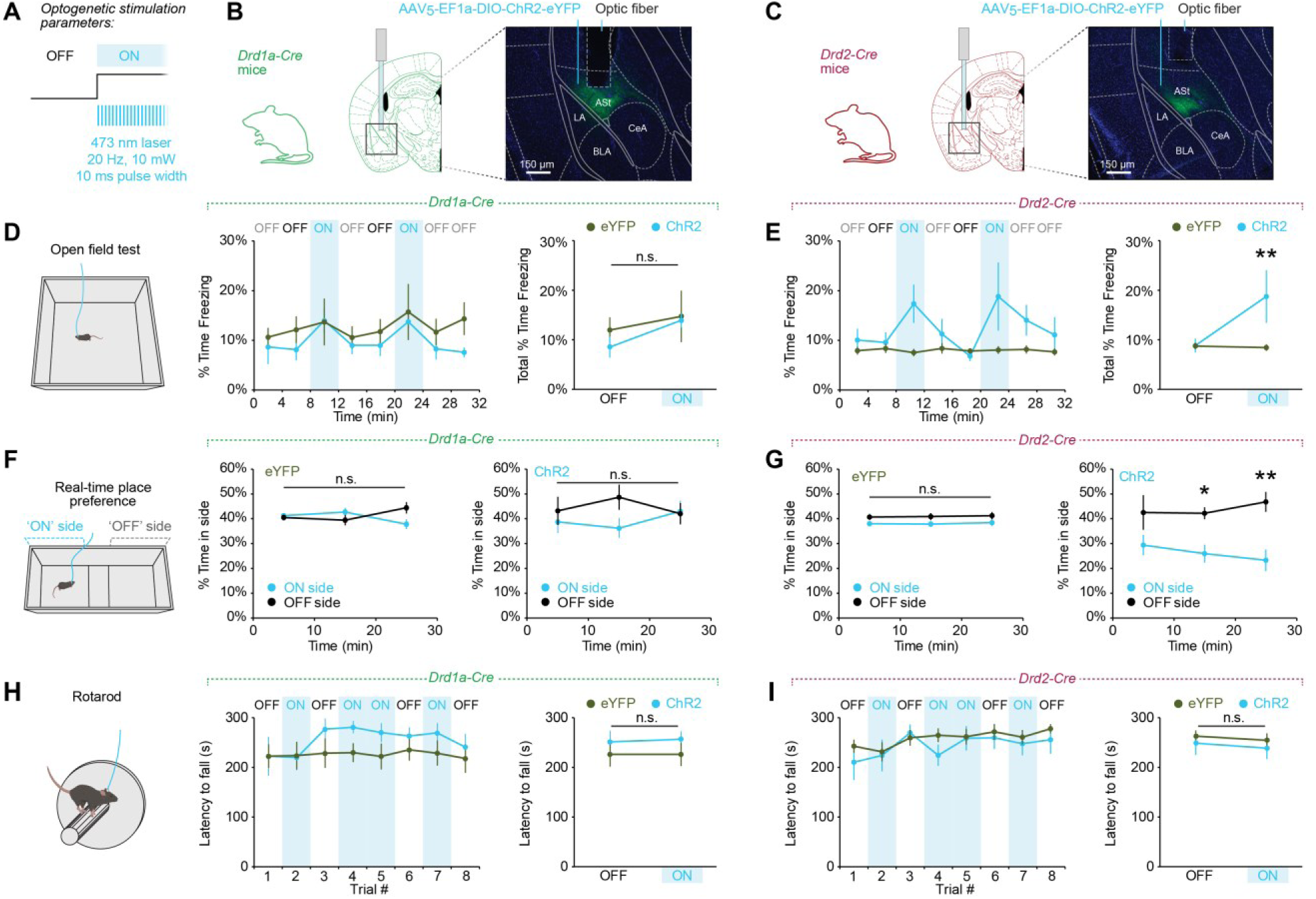
Activation of *Drd2*+ ASt neurons drives freezing and avoidance behaviors but does not disrupt gross motor coordination. **(A)** Schematic indicating optogenetic stimulation paratmeters. **(B-C)** Targeting strategy and representative image of ChR2 expression in *Drd1a+* ASt neurons (B) and *Drd2+* ASt neurons (C). **(D)** Optogenetic stimulation of *Drd1a+* ASt neurons had no effect on overall freezing during an open field task (two-way ANOVA, group × laser interaction F(1,22) = 0.1124, *p* = 0.7406; *Drd1a*-Cre ChR2: *n* = 5 mice, eYFP: *n* = 8 mice). **(E)** Optogenetic stimulation of *Drd2*+ ASt neurons resulted in a significant increase in freezing during light *ON* epochs compared with the *OFF* epochs preceding them (two-way ANOVA, group × laser interaction F(1,40) = 8.152, *p* = 0.0068; Bonferroni post hoc analysis \*\**p* < 0.01; *Drd2*-Cre ChR2: *n* = 6 mice, eYFP: *n* = 16 mice). *OFF* epochs labeled with black font indicate those used for analysis; *OFF* epochs with gray labels were excluded. **(F)** Optogenetic stimulation of *Drd1a+* ASt neurons in a real-time place preference (RTPP) task did not result in any change in preference for the side paired with stimulation (*ON* side) (two-way RM ANOVA, ChR2: effect of side F(1,8) = 0.6705, *p* = 0.4366; eYFP: effect of side F(1,12) = 0.2469, *p* = 0.6282; *Drd1a*-Cre ChR2: *n* = 5 mice, eYFP: *n* = 7 mice). **(G)** Stimulation of *Drd2*+ ASt neurons resulted in decreased time spent in the “on side,” indicating stimulation was aversive (two-way RM ANOVA, ChR2: effect of side F(1,10) = 9.117, *p* = 0.0129; eYFP: effect of side F(1,30) = 3.442, *p* = 0.0734; Bonferroni post hoc analysis \**p* < 0.05, \*\**p* < 0.01; *Drd2*-Cre ChR2: *n* = 6 mice, eYFP: *n* = 16 mice). **(H)** Optogenetic stimulation of *Drd1a+* ASt neurons had no effect on latency to fall in an accelerating rotarod task (two-way ANOVA, group × laser interaction F(1,18) = 0.0160, *p* = 0.9007; *Drd1a*-Cre ChR2: *n* = 5 mice, eYFP: *n* = 6 mice). **(I)** Optogenetic stimulation of *Drd2*+ ASt neurons had no effect on latency to fall in an accelerating rotarod task (two-way ANOVA, group × laser interaction F(1,38) = 0.00295, *p* = 0.9570; *Drd2*-Cre ChR2: *n* = 6 mice, eYFP: *n* = 15 mice). Data presented as mean ± SEM.

### *Drd2+* ASt neurons show robust responses to conditioned stimuli predicting shocks

We next used cell-type-specific calcium imaging to determine how *Drd1a+* and *Drd2+* ASt neurons encode different conditioned stimuli. We expressed GCaMP7f in these populations and imaged calcium transients via a head-mounted miniature microscope^63^ during a two-tone discrimination task, modified with shorter cues (10 s) to minimize imaging time and photobleaching on each trial (Figures 5A–5K and S13). At the population level, *Drd2+* ASt neurons showed robust responses to CS-shock over both short (0–1 s) and long (1–10 s) timescales that were significantly greater than unpaired controls (Figure 5J). *Drd1a+* ASt neuron population average responses to CS-shock were not significantly different from unpaired controls (Figure 5H). At the individual neuron level, considerable heterogeneity of responses could be seen in both populations. To better characterize the diversity of responses seen, we performed hierarchical clustering of neurons based on their responses to each CS (Figure 5L) and sorted these clusters based on the proportion of *Drd2+* to *Drd1a+* neurons in each (Figure 5M). We found that the clusters with more *Drd2+* neurons had greater responses to CS-shock and showed rapid or sustained responses to CS-shock (clusters 3, 2, 1, and 4) or were inhibited by CS-reward (cluster 7). By contrast, the clusters with more *Drd1a+* neurons had greater responses for CS-reward; they were either inhibited to CS-shock (clusters 9 and 6) or excited by CS-reward (cluster 5). *Drd1a+* neurons were the majority of the “low responding cluster” (cluster 8), though this group did show a small transient response preference for CS-shock. All clusters contained cells from both cell types, indicating heterogeneous responses to the two cues within each population. However, it was clear that Drd2+ neurons were the largest contributor to conditioned responses to CS-shock in the ASt. The rapid onsets and sustained responses seen in *Drd2+* neurons were also consistent with the requirements for directing defensive responses across behavioral timescales.

**Figure 5.**
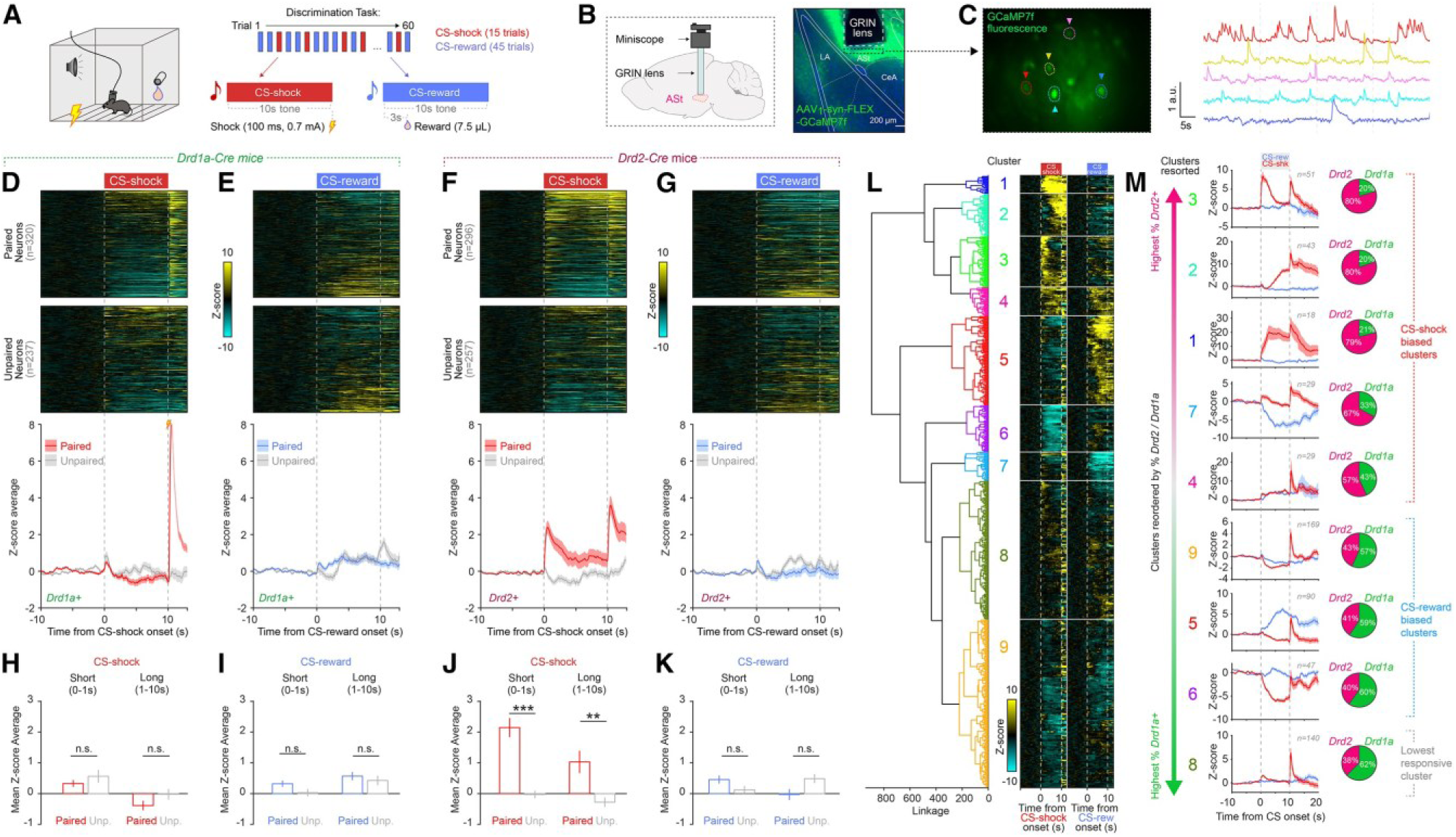
*Drd2*+ ASt neurons exhibit robust conditioned responses to CS-shock. **(A)** Two-tone discrimination task design for *in vivo* calcium imaging of ASt neurons. **(B)** GCaMP7f expression and GRIN lens targeting to ASt. **(C)** Calcium imaging field of view and representative traces of fluorescence changes in individual neurons. **(D–G)** Heatmaps (top) and group average traces (bottom) of ASt neuron *Z* score responses to conditioned stimuli in paired and unpaired group mice. (D) *Drd1a+* ASt neuron responses to CS-shock. (E) *Drd1a+* ASt neuron responses to CS-reward. (F) *Drd2*+ ASt neuron responses to CS-shock. (G) *Drd2*+ ASt neuron responses to CS-reward. **(H and I)** Average *Drd1a+* ASt neuron paired group responses to CS-shock (H) and CS-reward (I) were not significantly different from unpaired controls (CS-shock: short: *t555* = −1.07, *p* = 0.286, long: *t555* = −1.58, *p* = 0.116, CS-reward: short: *t555* = 1.86, *p* = 0.064, long: *t555* = 0.70, *p* = 0.482; *Drd1a+*: *n* = 10 mice, 320 neurons paired group, *n* = 8 mice, 237 neurons unpaired group). **(J)** *Drd2*+ ASt neurons paired group average responses to CS-shock were significantly greater than unpaired controls (short: *t551* = 6.21, *p* < 0.0001; long: *t551* = 3.16, *p* = 0.00163). **(K)** *Drd2*+ ASt neuron responses to CS-reward were not significantly different from unpaired controls (short: *t551* = 1.85, *p* = 0.065, long: *t551* = −1.68, *p* = 0.094; *Drd2+*: *n* = 9 mice, 296 neurons paired group; *n* = 6 mice, 257 neurons unpaired group). **(L)** Agglomerative hierarchical clustering of calcium imaging responses of neurons from *Drd1a-Cre* and *Drd2-Cre* paired group mice. **(M)** Clusters resorted by normalized % composition of *Drd2*+ /*Drd1a+* neurons. Mean *Z* score traces of responses to CS-shock and CS-reward (middle) and proportion of *Drd1a-Cre* and *Drd2-Cre* neurons in each cluster (right). Clusters with the highest *Drd2*+ proportion preferentially responded to CS-shock (clusters 3, 2, 1, and 4) or were inhibited by CS-reward (cluster 7). The clusters with a high *Drd1a+* proportion preferentially responded to CS-reward (cluster 5) or were inhibited by CS-shock (clusters 9 and 6). A large proportion of *Drd1a+* neurons was also seen in cluster 8, which had the lowest-magnitude responses to stimuli but did show a bias toward CS-shock.

In addition to conditioned cue responses, we also analyzed ASt *Drd1a+* and *Drd2+* neuron responses to unpredicted presentations of the shock and reward US (Figure S14A). Subsets of both populations responded to each US type, and both *Drd1a+* and *Drd2+* neurons showed scaling of responses to US-shock intensity, though not US-reward (Figures S14B–S14I). We examined co-responsiveness of neurons to different CS and US pairings and saw considerable heterogeneity in response types (Figures S14J–S14Q). Notably, a large proportion of *Drd2+* ASt neurons excited by CS-shock were also excited by the US-shock (Figure S14L), while few ASt neurons showed “salience” responses in the same direction to rewarding or aversive stimuli. In the case of CS responses in *Drd2+* ASt neurons (Figure S14Q), only 2.4% of neurons exhibited responses consistent with salience (excited or inhibited to both CS-shock and CS-reward), while 38% of neurons significantly responded to only one CS type or responded in opposite directions to CS-shock and CS-reward, consistent with cue discrimination or valence encoding.

We also examined whether ASt neurons encoded conditioned responses to contexts (Figure S15). However, following contextual fear conditioning, neither *Drd1a+* nor *Drd2+* ASt neurons showed any changes in neural activity in the conditioned context compared with an unconditioned one (Figures S15F–S15K). Consistent with our electrophysiology recordings, we found ASt neuron activity was also not correlated with bouts of freezing behavior during the task (Figures S15L–S15Q). These results indicated that ASt neuron encoding of conditioned stimuli was specific to sensory cues and not contexts.

### *Drd2+* ASt neurons are necessary for the expression of conditioned fear behaviors

Finally, we examined whether ASt activity was in fact necessary to direct the expression of conditioned defensive responses to cues. We bilaterally targeted a construct conditionally expressing the inhibitory opsin halorhodopsin (NpHR) to the ASt in *Drd2-Cre* mice (Figures 6A, 6B, and S16B). Mice were first trained to distinguish CS-shock and CS-reward cues and then examined in a version of the discrimination task where 593 nm light was delivered via optic fibers to the ASt on an interleaved subset of trials (laser *ON* trials), allowing us to assess changes in conditioned behaviors caused by silencing *Drd2+* ASt neurons, compared with trials without inhibition (laser *OFF* trials). We found that optogenetic inhibition of *Drd2+* ASt neurons caused a striking reduction in conditioned defensive responses to CS-shock during laser *ON* trials compared with laser *OFF* trials in NpHR group mice (Figures 6C and 6E). Remarkably, we also observed a significant reduction in both freezing and darting when analyzed separately (Figures S16G–S16J), consistent with our electrophysiology data finding that the ASt encodes the CS-shock cue rather than motor output (Figures 3E– 3H). Inhibition of *Drd2+* ASt neurons had no effect on responses to CS-reward (Figures 6D and 6F).

**Figure 6.**
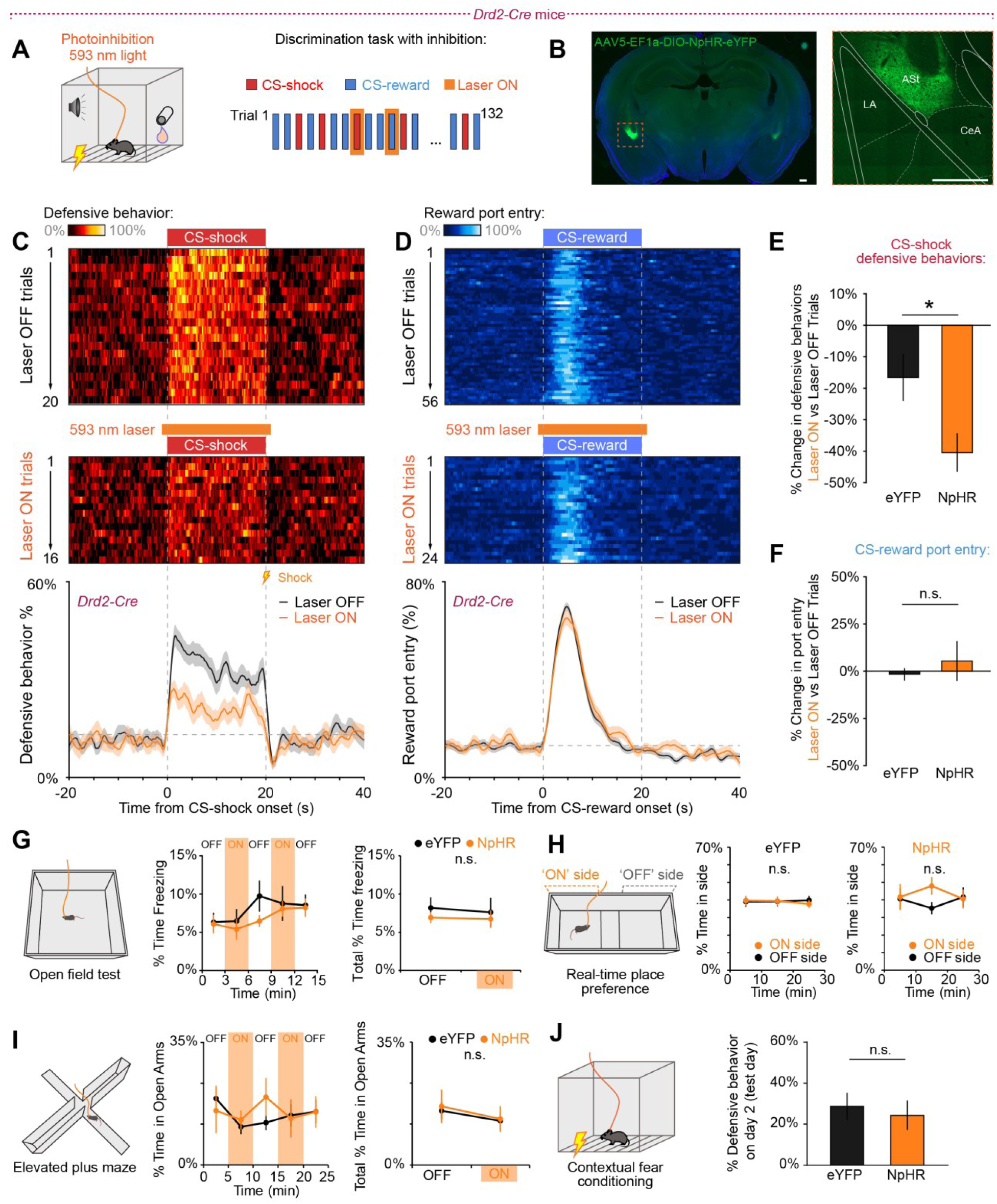
*Drd2*+ ASt neurons are necessary for expression of defensive responses to threat cues. **(A)** Discrimination task with optogenetic inhibition of *Drd2*+ ASt neurons on an interleaved subset of trials. **(B)** Representative images of AAV-DIO-NpHR-eYFP expression in *Drd2*+ ASt neurons (scale bars, 250 μm). **(C)** Defensive behavior heatmaps for CS-shock laser *OFF* (top) and laser *ON* trials (middle), and average defensive behavior during CS-shock trials with and without photoinhibition (bottom). **(D)** Port entry behavior heatmaps (top, middle) and group averages (bottom) during CS-reward trials with and without photoinhibition. **(E)** Inhibition of *Drd2*+ ASt neurons significantly reduced conditioned defensive behaviors to CS-shock compared with eYFP controls (t19 = 2.235, \**p* = 0.038, Student’s *t* test; NpHR: *n* = 8 mice, eYFP: *n* = 13 mice). **(F)** Inhibition of *Drd2*+ ASt neurons had no effect on port entry during CS-reward (*t19* = 0.809, *p* = 0.428, Student’s *t* test; NpHR: *n* = 8 mice, eYFP: *n* = 13 mice). **(G)** Optogenetic inhibition of *Drd2*+ ASt neurons had no effect on overall locomotion in an open field task (two-way ANOVA, group × laser interaction, *F(1,32)* = 0.011, *p* = 0.9167; NpHR: *n* = 5 mice, eYFP: *n* = 13 mice). **(H)** Inhibition of Drd2+ ASt neurons did not cause preference or aversion for an area paired with inhibition in an RTPP task (two-way RM ANOVA, NpHR: effect of side F(1,8) = 0.6896, *p* = 0.4304; eYFP: effect of side F(1,24) = 0.022, *p* = 0.8833). NpHR: *n* = 5 mice, eYFP: *n* = 13 mice. **(I)** Inhibition of *Drd2*+ ASt neurons caused no overall changes in exploration of open arms in an elevated plus maze (two-way ANOVA, group × laser interaction *F(1,32)* = 0.011, *p* = 0.9177; NpHR: *n* = 5 mice, eYFP: *n* = 13 mice). **(J)** Inhibition of *Drd2*+ ASt neurons had no effect on defensive behaviors on day 2 of a contextual fear conditioning task (Student’s *t* test *t16* = 0.395, *p* = 0.6984; NpHR: *n* = 5 mice, eYFP: *n* = 13 mice).

Inhibition of *Drd2+* ASt neurons had no effect on overall locomotion or unconditioned freezing behavior in an open field task (Figure 6G), indicating that the changes in conditioned behaviors observed in NpHR group mice during the discrimination task were not simply due to gross changes in motor output. Inhibition of *Drd2+* ASt neurons also had no effect on preference for the laser *ON* side in a real-time place preference task (Figure 6H) and did not cause any overall change in preference for open arms in an elevated plus maze task (Figure 6I). Inhibition of *Drd1a+* ASt neurons had no effect on responses to either CS type or other behaviors examined (Figures S16 and S17).

Interestingly, inhibition of *Drd2+* ASt neurons also had no effect on freezing to a conditioned context compared with eYFP controls (Figure 6J). This further showed that inhibition of *Drd2+* ASt neurons was not simply driving a motor effect that prevented mice from freezing altogether and that the *Drd2+* ASt neuron population was critical for defensive behavioral responses to conditioned cues but not contexts.

## DISCUSSION

### The ASt encodes sustained responses to conditioned threat cues

Here, we demonstrate the ASt is a genetically distinct brain region that plays a critical role in encoding conditioned stimuli to direct behavioral responses, a previously uncharacterized role for this structure.

A striking feature of ASt neurons was the sustained activity seen throughout the full duration of shock-predicting conditioned stimuli, which was greater in the ASt than in the adjacent TS or CeA. In the brain, sustained responses can be seen at both “lower” and “higher” level representations of stimuli. Sensory regions such as the thalamus have been shown to represent stimuli for their full duration,^64,65^ but sustained responses are also seen in the prefrontal cortex (PFC), where, in both primates and humans, they serve as high-level neural correlates for the representation of task features.^66^ Rodents trained in similar behavioral paradigms to this study have also shown sustained responses to conditioned stimuli in PFC neurons but comparatively transient responses in the amygdala.^67,68^ The ASt appears unique in that it is integrated with amygdalar circuits^20,33,36^ but shows both rapid and sustained activity tracking the full duration of threat cues. While transient responses to stimuli may be suitable to trigger aspects of emotional responses such as endocrine and stress hormone responses,^69^ we propose that a neural correlate tracking the full duration of aversive stimuli is ideally suited to orchestrate rapid and continuous behavioral responses necessary for survival.

Consistent with this proposed role, we found that activity of ASt neurons does indeed mediate defensive behaviors. ASt activity was correlated with total defensive responses to CS-shock; stimulation of ASt neurons was sufficient to drive freezing; and, most critically, inhibition of *Drd2+* ASt neurons caused a striking reduction (nearly 50%) in defensive behaviors in response to a shock-predicting cue. Our findings also consistently supported ASt activity encoding threat cues, rather than simply motor output. Analysis of subsecond behavior during electrophysiology recordings revealed that ASt neuron firing was not aligned with individual bouts of either freezing or darting, and logistic regression decoding and GLM analyses further showed that ASt activity predominantly represented the conditioned cue rather than specific behaviors. Inhibition of *Drd2+* ASt neurons also reduced *both* freezing and darting to the cue, rather than driving a unidirectional change in motor output. Together, these findings suggest the ASt does not encode individual defensive movements directly but rather represents the threat cue itself to direct behavioral responses.

### The ASt in models of amygdala and striatum function

Importantly, our findings complement rather than contradict the large body of existing work examining amygdala circuits mediating learned associations. Many studies have shown outputs from the CeA to the periaqueductal gray (PAG) are essential for expression of conditioned fear,^3,5,6^ and ASt neurons likely act in concert with these canonical circuits to drive defensive behavior. However, in some cases the ASt and amygdala may have distinct roles. The BLA and CeA are essential for contextual fear conditioning,^70,71^ but our calcium imaging and inhibition experiments both showed that ASt neurons did not appear to play a role in conditioned responses to contexts. Interestingly, the hippocampus, a key structure for spatial and contextual learning,^72,73^ projects to the BLA but not the ASt.^74^ The ASt may therefore be specialized for responses to acute sensory cues, while amygdala circuits play a central role in responses to context.

How does the ASt fit in the hierarchy of established circuit models for directing behavior? With respect to the amygdala, two major models are possible. First, the ASt may be “downstream” of the amygdala complex. The ASt receives robust input from the LA,^38,39^ and so it is possible that learned associations encoded in the LA^13,20^ are routed to the ASt and are necessary for ASt neurons’ conditioned responses to stimuli. In this model, the ASt could be considered an “output nucleus” of the amygdala complex like the CeA, with a specialized role for defensive responses to threat cues. Alternatively, the ASt may act as a “parallel pathway” to the amygdala, capable of encoding conditioned stimuli and directing behavioral responses independently. The ASt receives the same thalamocortical inputs as the LA^25,37^ and a dopaminergic signal that encodes novelty and external threat,^75^ which could facilitate synaptic plasticity in the ASt to encode learned associations without amygdala input. It is also possible that amygdala input could play a role in acquisition, rather than expression, of conditioned responses in the ASt. Determining the role of the LA→ASt projection and examining plasticity in the ASt during learning will be essential next steps to distinguish between these models. Additionally, the ASt lies directly in the dorsal path of manipulations targeting the amygdala, but off-target effects on ASt function have not been considered as a confound in past studies; careful histological controls will be necessary to disentangle the roles of these two structures.

Within the striatum, the unique features of the ASt support a specialized functional role. The ASt receives robust input from the amygdala,^38,39^ cortex,^76^ and thalamus,^25,37^ positioning it to connect circuits for threat detection and evaluation to downstream basal ganglia pathways mediating action selection. The ASt is also markedly enriched for *Drd2+* neurons, which are broadly implicated in responses to aversive stimuli,^43,44,77^ consistent with the predominant role of the ASt in responses to threat cues that we observed. Different striatal subregions receive distinct inputs,^78,79^ but their projections converge on shared downstream target structures of the direct and indirect pathways.^80^ *Drd2+* neurons are part of the striatal indirect pathway, which is posited to suppress competing motor outputs.^45,81^ The sustained ASt activity encoding threat cues observed in this study could therefore be integrated with signals from other striatal regions ^16,43,44,82^ in downstream target structures, suppressing some actions to bias selection toward defensive behaviors.

In recent years, a growing body of evidence has implicated the TS in responses to threats. The TS receives a dopaminergic projection from the SNL that encodes novelty and external threat^75,83,84^ rather than the classic prediction error signals typically reported in medial dopamine neurons,^75^ and this projection from the SNL can also be seen to collateralize to the ASt.^75^ Further, the SNL→TS dopaminergic projection has been shown to reduce fear learning and expression.^15,16,85^ Inhibition of TS neurons themselves has also been shown to affect expression of fear responses.^16,86^ In response to a moving threat object, *Drd1a+* TS neurons have been shown to play a critical role in mediating avoidance behavior.^82^ In the ventral TS, *Drd1a+* neurons were also reported to play a role in learning and expression of cue-fear responses and motor-aligned activity.^86^ While our findings indicated *Drd2+* neurons had a central role in the ASt, these studies suggest that responses to different threats are mediated by variable involvement of *Drd1a+* and *Drd2+* neurons across different striatal subregions and behavioral paradigms.

### ASt neurons have distinct genetic identities

Examination of the cell-type composition and transcriptomic profile of the ASt revealed many notable features. The ASt had a greater proportion of *Drd2+* MSNs compared with other regions of the striatum, and ASt neurons were identifiable as genetically distinct subclusters within each MSN type relative to the TS, DS, and CeA. These differences in clustering were due to hundreds of DEGs in *Drd1a+* and *Drd2+* neurons from the ASt. Intriguingly, these DEGs included many synaptic adhesion molecules and axon guidance factors, which likely contribute to establishing the unique connectivity of the ASt.^25,38,39^ A major anatomical feature distinguishing the ASt is the density of input it receives from several key sources: the LA,^38,39^ calretinin-expressing thalamic neurons representing CS/US information,^25^ and parvocellular subparafascicular nucleus (SPFp) neurons carrying threat information of different modalities.^37^ All of these projections can be seen to richly collateralize to the ASt and LA but then fade in density in a dorsal gradient through the TS.^25,37–39^ Interestingly, this pattern of collateralization closely matched the two ASt-enriched genes we examined, *Cdh6* and *Slit2*, which showed high expression in the ASt and LA that decreased in a dorsoventral gradient throughout the TS. Across the striatum, many inputs and outputs vary topographically, including the cortical regions that project to each subregion of the striatum,^78,87,88^ and the changes in expression of genes in MSNs that we observed throughout different subregions could be an underlying molecular basis for establishing these broad patterns of connectivity.

### A role for the ASt in valence encoding and disorders

The ASt appears to serve a specialized role in mediating defensive responses to threat cues. We found that the ASt was enriched for *Drd2+* neurons that showed greater responses to CS-shock, and inhibition of these neurons only affected cue-conditioned defensive behaviors to that CS and not reward responses. Can these ASt responses be considered to encode “negative valence”? There are several points to consider with this interpretation. First, as discussed above, ASt neurons did not appear to reflect fear-conditioned contexts, which might be expected for “universal” valence-encoding cells. Our study also used only a single CS and US of each type, and while our results are compatible with valence encoding, they do not rule out ASt activity encoding specific stimulus identities, value, or modalities. The definition of valence and its standards for experimental demonstration remain an ongoing area of discussion,^89–93^ and we believe that concurrent examination of different modalities and magnitudes of stimuli will facilitate analysis of valence encoding in future studies.

Together, our findings establish the ASt as a missing piece of the circuits mediating learned associations and motivated behavior. Consequently, this study also identifies the ASt as a novel site of interest in neuropsychiatric disorders where responses to stimuli are disrupted, such as anxiety and PTSD.^94,95^ The amygdala has been extensively studied in these disorders,^8^ but the contribution of the ASt is unknown, and the distinct genetic identity and connectivity of the ASt may represent vital new targets for investigation of these disorders.

## RESOURCE AVAILABILITY

### Lead contact

Further information and requests for resources and reagents should be directed to and will be fulfilled by the lead contact, Kay M. Tye.

### Materials availability

This study did not generate any new, unique reagents.

### Data and code availability

snRNA-seq data have been deposited at GEO: GSE211437. Raw and processed data are made freely available at https://dandiarchive.org/dandiset/001203. Code freely available from our laboratory Github at https://github.com/Tyelab/Mills-et-al-2026.

## ACKNOWLEDGMENTS

We thank Chris Leppla and Praneeth Namburi for their contributions to the project. We also thank Alcino Silva, Peyman Golshani, Daniel Aharoni, Tristan Shuman, and Denise Cai for their assistance and contributions to UCLA Miniscope resources used in the project. K.M.T. is an HHMI Investigator and the Wylie Vale chair at the Salk Institute for Biological Studies.

This article is subject to HHMI’s Open Access to Publications policy. HHMI lab heads have previously granted a nonexclusive CC BY 4.0 license to the public and a sublicensable license to HHMI in their research articles. Pursuant to those licenses, the author-accepted manuscript of this article can be made freely available under a CC BY 4.0 license immediately upon publication.

This work was supported by the Waitt Advanced Biophotonics Core Facility of the Salk Institute (RRID: SCR_014838) with funding from NIH-NCI CCSG P30 CA014195, NIH-NIA San Diego Nathan Shock Center P30 AG068635, the Shared Instrumentation grant S10-MH124757 (Olympus FV3000), the Henry L. Guenther Foundation, and the Waitt Foundation.

Funding was provided by the JPB Foundation; the New York Stem Cell Foundation; the Klingenstein Foundation; the McKnight Foundation; the Salk Institute; the Howard Hughes Medical Institute; the Clayton Foundation; the Kavli Foundation; the Dolby Family Fund; R01-MH115920 and R37-MH102441 (NIMH); Pioneer Award DP1-AT009925 (NCCIH); the CIHR Postdoctoral Fellowship (F.M.); the K99/R00 NIH Pathways to Independence Award (K99 MH121563 to F.M., K99 DA055111-01 to H.L., and K99 AA029180 to R.R.P.); the Margolis Scholar Award (F.M.); China Scholarship Council grant 201806010370 (S.S.); and the NIH/NIMH Blueprint D-SPAN Award (K00 MH132569) (C.C.).

## AUTHOR CONTRIBUTIONS

Conceptualization, F.M. and K.M.T.; methodology, K.M.T., F.M., J.R.H., and F.H.T.; software, L.M. and T.D.P.; formal analysis, F.M., C.R.L., J.R.H., H.L., M.N.K., S.S., F.H.T., M.E.L., F.A., J.W., M.G.C., M.B., L.R.K., F.B., D.P.L., K.J.L., A.L.G., J.D., C.C., C.R.H., L.M., T.D.P., and K.B.; investigation, F.M., C.R.L., J.R.H., H.L., M.N.K., S.S., M.E.L., F.A., J.W., M.G.C., M.B., H.S.C., G.P.S., A.L.G., R.R.P., M.M.A., R.W., and N.B.P.; data curation, F.M., J.R.H., L.R.K., R.W., and K.M.T.; writing, F.M., J.R.H., M.N.K., and K.M.T.; supervision, F.M., R.W., T.D.P., M.K.B., C.M.R., and K.M.T.; funding acquisition, F.M., T.D.P., M.K.B., C.M.R., and K.M.T.

## SUPPLEMENTAL INFORMATION

Supplemental information can be found online at https://doi.org/10.1016/j.neuron.2026.08.012.

Document S1. Figures S1–S17

Data S1. Histology

Document S2. Article plus supplemental information

## METHODS

### Experimental model and study participant details

Adult male wild-type C57BL/6J mice (RRID:IMSR_JAX:000664), *Drd1*a-Cre mice (RRID:MMRRC_030989-UCD), *Drd2*-Cre mice (RRID:MMRRC_032108-UCD), and *VGAT*-Cre mice

(RRID:IMSR_JAX:028862) were used in all experiments as stated in the text. Mice were housed in the Salk Institute for Biological Studies on a 12-h reverse light/dark cycle with ad libitum access to food and water except when otherwise stated in experimental procedures. All experiments were conducted during the dark cycle phase. All experimental procedures were carried out in accordance with NIH guidelines and approval of the Salk Institutional Animal Care and Use Committee.

### Method details

#### Stereotactic surgery procedures

All surgeries were conducted under aseptic conditions. Mice were anesthetized with an isoflurane/oxygen mixture (4-5% for induction, 1-2% for maintenance) and placed in a stereotaxic head frame (David Kopf Instruments, Tujunga, CA, USA). A heating pad was placed beneath the mice to maintain body temperature, and Sterile Lubricant Eye Ointment (Stye, INSIGHT Pharmaceuticals Corp., Langhorne, PA) was applied to the eyes to prevent drying. An incision was made along the midline to expose the skull and a dental drill was used to perform a craniotomy. During all surgeries, animals were injected subcutaneously with 1 mL of Ringer’s solution, Buprenorphine-ER (1 mg/kg), and Meloxicam (5 mg/kg). For recovery animals were placed in a clean cage on a heating pad. Animals were given >7 days of recovery period before being subjected to behavioral paradigms.

Stereotaxic coordinates were measured relative to bregma. Coordinates for injections targeting amygdalostriatal transition zone (ASt) were +/- 3.17 mm ML (mediolateral), -1.52 mm AP (anteroposterior), and -4.28 mm DV (dorsoventral) relative to bregma. All injections of viral vectors were performed using glass pipettes (Drummond Scientific) pulled to a 130-140 μm tip diameter with a pipette puller (Narishige PC-10, Amityville, NY, USA). Pipettes were either attached to 10 μL microsyringes (Hamilton Microlitre 701, Hamilton Co., Reno, NV, USA) with a microsyringe pump (UNP3; WPI, Worcester, MA, USA) and digital controller (Micro4; WPI, Worcester, MA, USA), or to the Nanoject III Programmable Nanoliter Injector (Drummond Scientific, Broomall, PA, USA) with digital controller (Drummond Scientific, Broomall, PA, USA). For each injection, micropipettes were slowly lowered to the target site and viral vectors were delivered at a rate of 0.1-5.0 nL per second. The pipette was then slowly raised 0.02 mm and left in place for 15-20 min to allow diffusion of the virus, then slowly withdrawn.

#### Surgery for *in vivo* electrophysiology

For single-wire *in vivo* electrophysiology recordings an electrode consisting of a bundle of 16 single nichrome wires of 9 μm diameter (Stablohm 675, CFW Material #: 100-188, California Fine Wire Company, Grover Beach, CA, USA) was implanted unilaterally in the ASt. Electrodes were lowered into the brain at a rate of 0.01 mm/s. Electrodes were secured with C&B-Metabond Quick adhesive Luting Cement (Parkell, Long Island, NY, USA) and dental cement (Ortho-Jet powder, Lang Dental, Wheeling, IL, USA). Two to three skull screws (00-96 x 1/16 (stainless steel) 1.6mm cut length, Plastics One) were secured in the skulls around the implanted electrode, with a layer of C&B-Metabond Quick adhesive Luting Cement and dental cement for stabilization. For additional *in vivo* electrophysiology experiments, animals were implanted with Neuropixels probes. Neuropixels probes (Version 1.0) were assembled into a custom-made 3D printed probe holder prior to implantation. Before surgery, the probe shank was soaked in 70% ethanol for 20 minutes. Petroleum jelly was gently applied around the base of the probe shank to form a hermetic seal. The remaining length of the probe shank was dipped into a fluorescent DiI (Vybrant DiI Cell-Labeling Solution, Waltham, MA, USA). For implantation, the probe holder with the probe was attached to a stereotaxic arm and centered at ML ± 3.17 mm, AP -1.52 mm relative to bregma. The probe was then lowered to DV -5.00 mm relative to bregma at a rate of 0.01 mm/s.

### Surgery for optogenetic manipulations

For optogenetic stimulation experiments mice received bilateral injections of 50 nL AAV_5_-DIO-ChR2-eYFP in the ASt. For optogenetic inhibition experiments, mice received bilateral injections of 50 nL AAV_5_-DIO-NpHR-eYFP in the ASt. For both experiments a 300-μm diameter optic fiber (0.37 numerical aperture) encased in a 1.25 mm ferrule was then implanted with the fiber tip 0.30 mm above the injection site. Identical parameters were used for the non-opsin controls except the viral vector injected was AAV_5_-DIO-eYFP.

#### Surgery for *in vivo* calcium imaging

For calcium imaging experiments, 50 nL of AAV_1_-Syn-FLEX-GCaMP7f was injected in the ASt at a rate of 1 nL per second. Following injection, a 0.5 mm x 6.1 mm 0.5 pitch gradient refractive index (GRIN) lens (Inscopix Inc, Mountain View, CA, USA) was centered over the injection site and lowered to a depth 0.05 mm above the injection site at a rate of 0.01 mm/s. On the contralateral side, a screw (00-96 x 1/16 (stainless steel) 1.6mm cut length, Plastics One) was secured into the skull. Both the lens and the skull screw were secured with super glue (Krazy Glue All Purpose, The Original Super Glue Corporation, Ontario, CA, USA) and dental cement (Ortho-Jet powder, Lang Dental). A small platform flush with the GRIN lens was created with dental cement, and a plastic cover fixed over the GRIN lens with dental cement to prevent damage to the lens prior to securing the baseplate. For v 3.2 Miniscope recordings, approximately 4 weeks following implantation, a 1.8 mm objective 0.25 pitch GRIN lens (Edmund Optics, Barrington, NJ) was placed over the GRIN lens with dental cement at a height resulting in the optimal focal plane. A small baseplate (miniscopeparts.com) was then secured to the previously formed dental cement.

### Electrode construction

For recording single-unit activity, multi-electrode arrays were constructed. Arrays consisted of one multi-channel single wire probe (Omnetics Connector Corp., Minneapolis, MN) as the structural component. This array was connected directly to the plug using a plastic spacer. The array was comprised of 16 single wires (Stablohm 675, CFW Material #: 100-188, California Fine Wire Company, Grover Beach, CA). Wires were aligned using the length of a syringe needle (2-3 mm). The wire insulation was then stripped using forceps and wires were connected to the Omnetics connector using a conductive adhesive (Super Shield Silver Conductive Paint; MG Chemicals, Burlington, ON). Super glue and epoxy were used to ensure connections were secure. Serrated tungsten scissors (Fine Science Tools, Foster City, CA) were used to cut the arrays to length. A 50/50 gold plating solution consisting of gold solution (Neuralynx, Bozeman, MT) and 1 μM polyethylene glycol was then applied to decrease the impedance of the wires to 150-250MΩ. Finally, a low impedance bare silver wire (California Fine Wire, Grover Beach, CA) was soldered to the last pin on the connector, and then the connection was covered with dental cement.

### Behavioral tasks

All behavioral tasks involving optogenetic manipulation or calcium imaging were performed 6-8 weeks after injection of viral vectors to allow sufficient expression of opsins or genetically-encoded calcium indicators. For at least four days prior to behavioral experiments mice were habituated to experimenter handling and attachment of optic fiber patch cables to surgically implanted optic fibers or tethers for calcium imaging and electrophysiology recordings. All behavioral experiments except the discrimination task were analyzed using Ethovision XT software (Noldus, Wageningen, Netherlands) and recorded with a digital video camera above the test arena. For optogenetic loss-of-function experiments, mice received bilateral optogenetic stimulation to silence targeted ASt neurons in both hemispheres, and behavioral data was excluded from analysis if viral targeting or leakage was inadequate in either hemisphere, particularly if any expression was detected in the CeA. For optogenetic gain of function experiments, mice received unilateral stimulation in each hemisphere in two repetitions of each task, and data from stimulation of either hemisphere was excluded if histology revealed inadequate accuracy of targeting of the virus, with expression in the CeA again resulting in exclusion of that data, as possible off-target effects would present a confound to interpretation of results.

### Open field test

Mice were placed in a 53 x 50 cm arena with four transparent plexiglass walls, and allowed to move freely throughout the arena for 15 min. Light stimulation was delivered during the 3-6 min and 9-12 min ‘ON’ epochs (ChR2 mice and eYFP controls: 10 mW 473 nm light at 20 Hz, 5 ms pulse width. NpHR mice and eYFP controls: 3 mW 593 nm light, constant delivery). In the *Drd1a-Cre* and *Drd2-Cre* cohorts, a total task time of 32 min was used, with ‘ON’ epochs during 8-12 min and 20-24 min time windows, and a 10 ms pulse width. Overall locomotion and time spent in edges (50% of arena area closest to walls) and center region of arena during each epoch was analyzed for each mouse.

### Elevated plus maze

Mice were placed in the center of the elevated plus maze (EPM) apparatus, consisting of two open arms (30 x 5 cm) and two enclosed arms (30 x 5 x 30 cm) extending from the center platform (5 x 5 cm). The entire apparatus was elevated 75 cm from the floor. Mice were given 1-5 min to recover from handling before the testing phase began. Light stimulation was delivered during the 3-6 min and 9-12 min epochs (ChR2 mice and eYFP controls: 10 mW 473 nm light at 20 Hz, 5 ms pulse width. NpHR mice and eYFP controls: 3 mW 593 nm light, constant delivery). In the *Drd1a-Cre* and *Drd2-Cre* cohorts, a total task time of 32 min was used, with ‘ON’ epochs during 4-8 min and 20-24 min time windows, and a 10 ms pulse width. Total time spent in open and closed arms of the apparatus throughout each epoch was quantified.

### Real-time place preference

Mice were placed in the center of a transparent Plexiglass chamber (57.15 x 22.5 x 30.5 cm). The chamber was divided into ‘ON side’ and ‘OFF side’ compartments (counterbalanced between and within groups). The chamber was then illuminated with 30 lux ambient light. Mice were allowed to freely explore the chambers for 30 min during which entry into the ‘ON side’ triggered photostimulation (ChR2 mice and eYFP controls: 10 mW 473 nm light at 20 Hz, 5 ms pulse width, 10 ms pulse width for *Drd1*-Cre and *Drd2*-Cre cohorts. NpHR mice and eYFP controls: 3 mW 593 nm light, constant delivery) until the side was exited, and total time spent in each side throughout the task was quantified.

### Rotarod task

A rotarod (3 cm diameter rotating cylinder, San Diego Instruments) was used to assess motor coordination and balance in mice. Mice were trained on the rotarod for 3 to 6 consecutive days. During each trial, the speed of the rotating rod increased linearly to 16 RPM over the course of 120 seconds. Training continued until the group reached an average latency to fall of over 120 seconds at 16 RPM. Mice that successfully maintained balance for at least 120 seconds on the rotating rod advanced to the test day. On the test day, a ramp protocol was employed, during which the rod’s rotation speed gradually increased to 16 RPM over the course of 300 seconds. Each mouse was subjected to 8 test trials, with interleaved stimulation and no-stimulation conditions. During the stimulation trials, mice received photostimulation (ChR2 and eYFP mice: 10 mW, 473 nm light at 20 Hz, 10 ms pulse width). The control trials (no-stimulation trials) were conducted under identical conditions but without any stimulation. Each trial lasted for a maximum of 300 seconds or until the mouse fell from the rod, whichever occurred first. Latency to fall was detected by a beam-break sensor below the rod, and recorded for each mouse across all trials, and differences between stimulated and non-stimulated conditions were analyzed.

### Two-tone discrimination tasks

Animals were trained and tested in standard operant chambers (23 x 30 x 40 cm; Med Associates Inc.) within a sound attenuating cubicle. Each chamber was equipped with speakers for tone delivery, a syringe pump for reward delivery, a grid floor for shock delivery, infrared house lights and a camera for behavioral recording. A reward port with an infrared beam was used to detect port entries, and implanted in a custom-made 3D printed port which minimized the possibility of beam breaks not being associated with licking. In our experimental apparatus, it was not possible to use ‘lickometers’ as they rely on active electric current which can interfere with *in vivo* electrophysiology recordings. We visually monitored mice during tasks to ensure that beam breaks were accurately reflecting robust reward consumption, though it is possible that some instances of beam breaks may have been due to nose or limb pokes rather than licking the reward spout.

Prior to two-tone discrimination experiments, mice were sequentially trained to recognize reward-predicting and shock-predicting cues. In the first phase of training (Reward conditioning – 4 days) animals were trained to associate a conditioned stimulus (‘CS-reward’) with delivery of a high-calorie reward. Animals were presented with CS-reward trials in which a pure auditory tone cue (20 kHz or 3.5 kHz, counterbalanced) predicted the delivery of a reward (7.5 μL per trial, chocolate Ensure, Abbott Laboratories) to the reward port. Collection of the reward prior to the end of the tone resulted in termination of the tone. The inter-trial interval was variable with an average of 90 sec (+/- 20 sec). Unpaired mice were presented with the same number of CS-rewards and Ensure deliveries, explicitly unpaired from one another and presented in a randomized order. Animals underwent four days of reward training (1 session per day). On all tasks with rewards, mice were food-restricted (3.0 g food per mouse per day) prior to testing.

In the second phase of training animals were trained to associate a conditioned stimulus (CS-shock) with a shock delivery. During fear conditioning animals were presented with trials in which a tone cue (3.5 kHz or 20 kHz, counterbalanced, and reversed from the tone pairing used for CS-reward for each mouse) predicted the delivery of a 0.1 sec shock (0.7 mA). The delivery of the shock co-terminated with the predicting tone. ITI was variable with an average of 90 sec (+/- 20 sec). Unpaired mice were presented with the same number of CS-shocks and shock deliveries, explicitly unpaired from one another and presented in a randomized order. Animals underwent two sessions of fear training (‘training’ and ‘test’ sessions, 1 session per day). In the second session (‘test’), animals were presented with tones only and no shocks.

Following these training phases to recognize each CS, mice were then trained in a two-tone discrimination task. During the discrimination task, animals were presented with both CS-reward trials (3.5 kHz or 20 kHz, counterbalanced) and CS-shock trials (20 kHz or 3.5 kHz, counterbalanced). Trials were presented in a pseudorandom manner, and the ITI was variable with an average of 90 sec (+/- 20 sec). Unpaired mice received the same number of CS and US deliveries, explicitly unpaired from one another and presented in a randomized order. Animals underwent discrimination testing for 4-6 sessions (1 session per day) until they achieved 85% reward collection.

For electrophysiology experiments the ‘CS-reward’ and ‘CS-shock’ conditioned tones associated with chocolate Ensure and shock delivery, respectively, were 20 seconds in length. 90 trials of CS-reward and 30 trials of CS-shock were presented in a single discrimination task session. Recordings from electrodes were collected continuously throughout the session.

For calcium imaging experiments, task details were similar, but the overall length of trials and number of trials was reduced to prevent photobleaching of GCaMP7f. Conditioned tones associated with the ‘CS-reward’ and ‘CS-shock’ were 10 seconds in length, and 45 trials of CS-reward and 15 trials of CS-shock were presented. Additionally, delivery of the reward was delayed to 3s following the beginning of CS-reward in order to allow a window of analysis of CS response free from US response. For the discrimination task, recordings from miniscopes were collected on select trials (every shock trial, every other reward trial), using a 5V TTL signal to trigger miniscope recordings. Blue LED houselights were also included in the operant chamber to reduce changes in illumination caused by activation of the blue excitation LED from the miniscope. In our initial cohorts, we recorded calcium imaging data from a 10s baseline period, the 10s period, and 3s post-CS. However, in subsequent cohorts, we extended the post-CS imaging window to 10s to allow visual confirmation that ASt neuron activity did indeed return to baseline after termination of the CS and shock. As a result, it can be seen that some cells’ responses terminate at 3s post-CS in the heatmaps and group averages of the calcium imaging discrimination task data. However, the data collected during the CS window was identical, and pooled for subsequent analysis of CS response magnitudes. In the group average traces, cells that were only recorded for 3s post-CS were excluded from the average trace in the 3-10s post-CS time period, not treated as ‘zero’.

For optogenetic inhibition experiments, the ‘CS-reward’ and ‘CS-shock’ conditioned tones associated with chocolate Ensure and shock delivery, respectively, were 20 seconds in length. On a subset of interleaved CS-shock and CS-reward trials both NpHR and eYFP control mice received laser delivery (593 nm light, 3 mW, constant delivery) beginning 1s prior and ending 1s after the tone on period. For CS-reward, 84 trials took place, and of these 56 were ‘Laser OFF’ trials with no inhibition, and 28 were ‘Laser ON’ trials where ASt neurons were inhibited. For CS-shock, 36 trials took place, and of these 20 were ‘Laser OFF’ trials with no inhibition, and 16 were ‘Laser ON’ trials where ASt neurons were inhibited. Again, a pseudorandom trial structure was used so that inhibited and non-inhibited trials of each CS type were presented in an interleaved and unpredictable manner throughout the entire discrimination task session.

### US experiment

For calcium imaging cohort mice, animals underwent an additional session of US testing to determine responses to both shock and reward when unpredicted by the CS. In this recording session, animals were presented with CS-shock and CS-reward cues as in the discrimination task, and in addition unpredicted foot shocks of different intensities (0.1 mA, 0.7 mA or 1.4 mA shock intensity, 250 ms duration) or unpredicted reward deliveries of different magnitudes (1.875, 3.75 or 7.5 μL chocolate Ensure), presented in a pseudo-random order.

### Contextual fear conditioning

For NpHR cohort mice, animals underwent two sessions of contextual fear conditioning over two days. On the first day (‘training’) mice were placed in a novel environment, allowed to habituate for five minutes, and ten foot shocks (1s, 0.7 mA) were delivered in a randomized order over ten minutes. On the second day (‘test day’) mice were returned to the environment, and both NpHR and eYFP control mice received laser delivery (593 nm light, 3 mW, constant delivery) during a 10 minute test session, and the behavioral responses to conditioned context were recorded.

For calcium imaging cohorts, a different task design was used to allow comparison of ASt neuron activity in conditioned and unconditioned contexts. Mice were first habituated to two contexts (Context A and Context B - two operant chambers which were in different rooms, and had distinct visual cues on the walls) for 30 mins per day each, for two days. Then, on day 3, mice underwent fear conditioning in Context A; after 5 mins of exposure, they received 10 foot shocks (0.5s, 0.7 mA) over 10 minutes with pseudorandom spacing. Then, on day 4, mice were returned to Context A (conditioned), and also context B (unconditioned), and neural activity was recorded in *Drd1a+* or *Drd2+* ASt neurons using 1-photon calcium imaging. Mice who failed to successfully acquire the conditioned context associations were removed from the analysis. The order of presentation of the two contexts was randomized across mice, and counterbalanced to prevent any order effects. Defensive behaviors in the conditioned context were quantified using SLEAP pose estimation, as in the discrimination tasks.

## Electrophysiology recordings

### *In vivo* electrophysiology data acquisition

Electrophysiology data from single-wire probes was recorded using an Open Ephys acquisition board^96^ in conjunction with 16 channel Intan headstages (Intan Technologies, Los Angeles, CA). Data was collected at a sampling rate of 30 kHz with a band pass filter to only collect signals between 1 and 7000 Hz.

### Neuropixels data acquisition

During behavior, Neuropixels data was acquired using SpikeGLX software (http://billkarsh.github.io/SpikeGLX/) via a National Instruments DAQ (PXIe-1071, PCIe-8381, and PXI-6133) at 30 kHz. Event timestamps were acquired in TTL signals to the National Instruments DAQ at 10 kHz.

## Calcium imaging

### Calcium imaging data acquisition

UCLA miniscopes v3.2 and v4 (http://miniscope.org) were used to collect calcium imaging data.^63^ During behavior, a 5V TTL signal was used to trigger the miniscope recording on select trials. When the scope was triggered, imaging data from the CMOS imaging sensor (Aptina, MT9V032) was transferred to data acquisition (DAQ) electronics and USB Host Controller (Cypress, CYUSB3013) via a coaxial cable. Images were acquired at 15 frames per second at a resolution of 752 x 480 and saved as uncompressed.avi files.

### Histology

Following experiments, mice were deeply anesthetized with sodium pentobarbital (200 mg/kg intraperitoneal injection). Animals were transcardially perfused with 10 mL of Ringer’s solution followed by 10 mL of cold 4% PFA in 1X PBS. Following decapitation and extraction, brains were fixed in 4% PFA in 1X PBS for 24 hrs at 4°C. Brains were then transferred to 30% sucrose in 1X PBS at 4°C until they sank to the bottom of the sucrose solution. Brains were sectioned coronally at 50 μm using a microtome (Thermo Scientific) and were stored at 4°C in 1X PBS until staining. Sections were mounted directly onto glass microscope slides and coverslipped with EMS-Shield Mounting Medium w/ DAPI. Slides were then imaged at 10x using the Keyence BZ-X710 Fluorescence microscope.

Histology from all mice was examined to determine accurate placement of virus and implants. For Neuropixels implants, the trajectory of each Neuropixels probe was reconstructed from brain slices based on The Allen Mouse Brain Common Coordinate Framework (https://www.sciencedirect.com/science/article/pii/S0092867420304025) using Allen CCF tools (https://github.com/cortex-lab/allenCCF).

## RNAscope *in situ* hybridization

### Preparation of fresh frozen sections for fluorescence *in situ* hybridization

8-10 week old C57BL/6J mice from Jackson Laboratories were anesthetized with 5% isoflurane, and brains were immediately extracted and covered with powdered dry ice for 2 minutes, and then stored at -80°C. Brains were then sectioned at 20 μm thick using a cryostat (CM3050 S; Leica, Buffalo Grove, IL) at -16°C. Sections were placed on a glass slide, gently heated from the bottom with the tip of a finger to encourage adhesion of the slice to the slide, and stored at -80°C until further processing.

### Fluorescence *in situ* hybridization using RNAscope

Fluorescence *in situ* hybridization was performed using the Advanced Cell Diagnostics bio V2 RNAscope kit and protocol (Advanced Cell Diagnostics, Newark, CA) using the RNAscope Multiplex Fluorescent Reagent Kit V2 (Catalog #323100), Fluorescent Multiplex Detection Reagents (#323110), probes for Drd1a (#406491-C1 and #406491-C3), Drd2 (#406501-C3 and #406501-C1), Cdh6 (#519441-C2), Slit2 (#449691-C1), and VGlut1 (#501101-C1), and the Perkin Elmer TSA Plus Fluorescence Palette Kit (NEL760001KT). Fresh frozen slices were fixed in 4% paraformaldehyde for 1 hour at 4°C. Slices were dehydrated in ethanol, and incubated in hydrogen peroxide for 8 minutes, with protease steps omitted to prevent tissue degradation. Slides were incubated in desired probe combinations for 2 hours at 40°C. For each channel, slides were incubated in channel-specific HRP for 10 minutes, followed by incubation in TSA fluorophore for 20 minutes, and then incubation in HRP-blocker for 10 minutes. Between all steps, slides were washed 2 times in 1x RNAscope wash buffer for 1-2 minutes. Slides were then incubated in ACDbio DAPI for 10 minutes, washed, dried for 20 minutes, coverslipped with PVA-DABCO (Millipore-Sigma), and left to dry overnight before imaging.

### Confocal microscopy

For the Drd1a/Drd2 labelling experiment, confocal fluorescence images were acquired on an Olympus FV1000 confocal laser scanning microscope using a 40x/1.30NA oil immersion objective, and serial Z-stack images were acquired using the FluoView software (Olympus, Center Valley, PA) at a thickness of 1.5 μm per Z-slice, with only the best-targeted Z-slice used for each target area. For the *Cdh6*/*Slit2* labelling experiment, settings were the same, except Z-slice thickness was 0.88 μm. For the VGlut1 labelling experiment, images were taken using a 20x objective at a Z-slice thickness of 3.7 μm (7-11 slices per section) with a max projection of all slices analyzed. All images in each experiment were acquired with identical settings for laser power, detector gain, and amplifier offset.

### Image processing

For subsequent image analysis, ROI masks of each region were drawn using *Paxinos and Franklin’s the Mouse Brain in Stereotaxic Coordinates* as reference^97^. For the *Drd1a*/*Drd2* labelling experiment, background was subtracted from all channels in all images using the subtract background feature in ImageJ. Each channel was then further thresholded using the brightness and contrast feature in ImageJ. Thresholds were determined by manually thresholding 10 images to a level with no background, and then computing the average minimum threshold necessary to remove background. The average threshold across 10 images was then used as the threshold for all images. Images were then exported to CellProfiler for further cell and puncta identification.

## Single-nucleus RNA sequencing

### Tissue extraction and cryopreservation

All procedures were performed in accordance with Institutional Animal Care and Use Committee protocol S16054 at the University of California, San Diego. All mice in the study were wild-type C57BL/6J background and received directly from Jackson Laboratories at 6 weeks of age and acclimated to the colony prior to experiments. Animals were single-housed and maintained free from noise or disturbance for 24 hours prior to sacrifice to reduce artifactual immediate early gene expression. Sacrifice was performed at P60 ± 3 days (*n* = 10-15 mice per pool for ASt, 10 for CeA, 9-10 for tail of striatum, 5 for body of striatum). Sample size was determined based on amount of expected nuclei per region per mouse: estimates of expected nuclei were determined empirically, though nuclear recovery was approximately 20% of total based on cellular density estimates from the Blue Brain Cell Atlas. 12,000 nuclei were targeted per combination of assay, condition, and region, which was determined using Single-Cell One-sided Probability Interactive Tool (SCOPIT) v1.1.4, allowing potential detection of at least 5 nuclei from 10 rare subpopulations at 0.1% frequency with 95% probability.^98^ All sacrifices were performed during the dark period of the light cycle. Animals were anesthetized via combined intraperitoneal injection of 150 mg/kg ketamine and 15 mg/kg xylazine. Once unconscious, mice were transcardially perfused with ice-cold, carbogen-bubbled (95% O_2_, 5% CO_2_), nuclease-free, 0.22 μm sterile-filtered artificial cerebrospinal fluid (ACSF) with a composition of 93 mM *N*-methyl-D-glucamine, 2.5 mM KCl, 1.2 mM NaH_2_PO_4_, 30 mM NaHCO_3_, 20 mM HEPES, 25 mM glucose, 5 mM sodium ascorbate, 2 mM thiourea, 3 mM sodium pyruvate, 13.2 mM trehalose, 12 mM *N*-acetyl-cysteine, 0.5 mM CaCl_2_, 10 mM MgSO_4_, and 93 mM HCl, at pH 7.3-7.4.^99,100^ Following transcardial perfusion, brains were immediately extracted and submerged into ice-cold carbogen-bubbled ACSF, with less than 5 minutes between the beginning of perfusion and final submersion after extraction. Brains were serially sectioned in ice-cold, carbogen-bubbled ACSF on a VT1000S vibratome (Leica) with polytetrafluoroethylene-coated razor blades (Ted Pella) at 0.15 mm/sec and 100 Hz, dividing the whole cerebrum into 400 μm coronal slices.

Due to the small size of the ASt and its immediate proximity to both striatal and amygdalar structures, accurate regional sampling was critical to ensure that transcriptomic comparisons reflected true biological differences rather than contamination from adjacent regions. To address this, we employed a stringent microdissection and histological verification pipeline. Target regions were microdissected from these slices under a stereomicroscope using a sterile blunt-end needle (22 gauge for CeA, ASt, and tail of striatum, 16 gauge for dorsal striatum). All regions were targeted using *Paxinos and Franklin’s the Mouse Brain in Stereotaxic Coordinates* as reference^97^. Extracted tissue samples were recovered in ice-cold, nuclease-free, 0.22 μm sterile-filtered cryoprotective nuclear storage buffer, composed of 0.32 M sucrose, 5 mM CaCl_2_, 3 mM magnesium acetate, 10 mM Trizma hydrochloride buffer (pH 8.0), 1 mM dithiothreitol, 0.02 U/μl SUPERase⋅ In RNAse Inhibitor (Invitrogen), and 1X cOmplete Protease Inhibitor Cocktail with EDTA (Roche). Tissue was then snap frozen using a metal CoolRack M90 (Biocision) pre-chilled to -80°C and stored at -80°C until nuclear isolation. Following extraction of tissue regions of interest, remaining portions of sections were fixed in 4% paraformaldehyde and 4′,6-diamidino-2-phenylindole (DAPI) was applied to sections at 1 μg/ml. After fixation and staining, sections were mounted and imaged on a VS120 slide scanner (Olympus). From these images, dissection accuracy was assessed for each region, and individual samples were only selected for downstream nuclear isolation if the extracted tissue fell entirely within the defined target regions.

### Nuclear isolation and sorting

Nuclear isolation procedures were adapted from multiple methods described previously.^101,102^ All procedures were performed on ice, and all solutions were ice-cold, nuclease-free, and 0.22 μm sterile-filtered. Cryopreserved tissue pieces were slow-thawed by incubation at 4°C for 1 hour prior to isolation. Tissue pieces were then pooled and resuspended in nuclear isolation medium composed of 0.25 M sucrose, 25 mM KCl, 5 mM MgCl_2_, 10 mM Trizma hydrochloride buffer (pH 7.4), 1 mM dithiothreitol, 0.04 U/μl RNasin Plus RNAse Inhibitor (Promega), 1X cOmplete Protease Inhibitor Cocktail with EDTA (Roche), and 0.1% Triton-X. The pooled tissue pieces in nuclear isolation medium were transferred to a 2 mL Dounce tissue grinder. Tissue was homogenized by 5 strokes from the loose pestle followed by 15 from the tight pestle, and the resulting homogenate was filtered through a 40 μm Flowmi cell strainer (Bel-Art) into a 1.5 ml Lo-Bind tube (Eppendorf). The homogenate was then centrifuged with a swinging bucket rotor at 4°C and 1000 x g for 8 minutes. Nuclei were then washed with nuclear flow buffer composed of DPBS with 1% bovine serum albumin, 1 mM dithiothreitol, and 0.04 U/μl RNasin Plus RNAse Inhibitor (Promega) and centrifuged at 4°C and 500 x g for 5 minutes, which was subsequently repeated. Nuclei were finally resuspended in nuclear flow buffer containing 3 μM DRAQ7 (Cell Signaling Technology) and again filtered through a 40 μm Flowmi cell strainer into a 5 ml round-bottom polystyrene tube. Each isolation took under 45 minutes to perform, from homogenization to final suspension.

Fluorescence-activated nuclei sorting (FANS) was carried out on a FACSAria II SORP (BD Biosciences) using a 70 μm nozzle at 52 PSI sheath pressure. For FANS, debris was first excluded by gating on forward and side scatter pulse area parameters (FSC-A and SSC-A), followed by exclusion of aggregates (FSC-W and SSC-W), and finally gating for nuclei based on DRAQ7 fluorescence (FSC-A and APC-Cy7-A). Only the set of DRAQ7+ nuclei at the first level of fluorescence were sorted to enrich for singlets, given stoichiometric DNA binding. Nuclei were successively sorted into 1.5 ml LoBind tubes (Eppendorf) under the purity sort mode. The tube contained 10X RT master mix without RT Buffer C. 16,000 total nuclei were targeted for downstream processing, and to account for cytometer errors and subsequent loss of nuclei, up to 21,000 detected nuclei were sorted into the tube to yield 16,000 viable nuclei for downstream sequencing (if fewer than 21,000 nuclei were detected, all were sorted into the tube). Nuclei were then immediately processed for snRNA-seq.

### Library preparation and sequencing

Nuclear suspensions were converted into barcoded snRNA-seq libraries using the Chromium Next GEM Single Cell 3’ v3.1 Reagent Kits v3.1 Single Index (10X Genomics). Library preparation for both assays was performed in accordance with the manufacturer’s instructions. 10,000 nuclei were targeted during each snRNA-seq library preparation run, and each region was performed in duplicate at minimum (n = 2 libraries for ASt, 3 for CeA and tail of striatum, 4 for body of striatum).

10X libraries were first sequenced at low depth on a NextSeq 550 Sequencing System (Illumina) to estimate quality and number of nuclei for each library, followed by deep sequencing on a NovaSeq 6000 Sequencing System. All runs were performed using 2 x 100-bp paired-end reads, outputting data in 28/8/91-bp read format. All sequences were demultiplexed using bcl2fastq.

## Quantification and statistical analysis

### Statistical analysis

Statistical analyses were performed using either GraphPad Prism (GraphPad Software, Inc, La Jolla, CA, USA) or MATLAB (Mathworks, Natick, MA, USA). Group comparisons were made using one-way, two-way, or repeated-measures analysis of variance (ANOVA). One-way ANOVAs and one-way repeated-measures ANOVAs were followed by Tukey’s multiple comparison test or Bonferroni-corrected multiple comparison tests, as indicated in each figure legend. Two-way and two-way repeated-measures ANOVAs were followed by Bonferroni post hoc tests. Non-parametric Wilcoxon signed-rank tests were used to compare in vivo neural firing rates across conditions, using an α = 0.01. Neurons were classified as significantly excited or inhibited when the signed-rank test reached α = 0.01 and the mean z-score exceeded +2 (excited) or fell below −2 (inhibited). Proportions of neurons excited or inhibited were compared across regions or groups using chi-square tests, with each family of comparisons corrected by the Holm-Bonferroni method. The number of animals and the number of neurons recorded is specified in the figures, the figure legends, and the text.

## Statistical analysis of behavior

### Behavioral analysis of open field test, elevated plus maze, and real-time place preference task

Behavioral performance was recorded by digital cameras positioned above the test arena. Ethovision XT software (Noldus, Wageningen, Netherlands) was used to track mouse location. For the open field test, overall locomotion and time in the edge and center regions were quantified per epoch, and for the elevated plus maze, time in the open and closed arms was quantified per epoch; these were analyzed by two-way ANOVA (group × laser epoch). For real-time place preference, the percentage of time spent on the ON and OFF sides was quantified across three time bins and analyzed separately for each genotype by two-way repeated-measures ANOVA (side × time bin, with time bin as the repeated measure). All analyses were followed by Bonferroni post hoc tests.

### Behavioral analysis of two-tone discrimination task

To automatically detect animal defensive behaviors, SLEAP^62^ was used to estimate animal poses in behavioral videos. A training data set of over 15,000 frames was labeled using a 14-point skeleton to represent the mouse (nose, left ear tip, left ear base, right ear base, right ear tip, skull base, shoulders, haunch, tail base, tail segment, left arm, right arm, right leg, and left leg). This data set was used to train a bottom-up model; in this model, first the animal is detected using a centroid model, then the location of each labeled body part is identified using a centered instance model.

For quantification of freezing, our defined criteria was when the total displacement of ALL mouse body points was less than 0.08 cm per frame. This threshold was empirically determined to capture near-total immobility of the mouse and using the displacement of all body points allowed us to distinguish freezing from time periods where the mouse was simply stationary, but grooming, sniffing or exploring. For quantification of darting, our defined criteria was when the center point of the mouse rear body reached a velocity of 45 cm/s. This was close to the maximum speed possible for mice in the limited area of the operant chambers (30 x 40 cm), and markedly distinct from normal locomotion. We visually inspected our behavioral videos overlaid with SLEAP coordinates and freezing/darting labels to confirm their accuracy. Total ‘defensive behaviors’ was calculated as the sum of freezing and darting behavior. To detect reward seeking behavior, port entries and exits were determined from reward port infrared beam breaks (Med-PC IV, Med Associates).

## Statistical analysis of electrophysiology recordings

### Neuropixels recordings

Acquired data were preprocessed using CatGT (https://billkarsh.github.io/SpikeGLX/#catgt). Preprocessed data were then sorted into spike clusters using Kilosort 2.5 (https://github.com/MouseLand/Kilosort), which provides a drift-tracking feature that detects the footprints of the same neuron during a recording session while correcting for drift. To determine the quality metrics of sorted spike clusters, we used modules for processing extracellular electrophysiology data provided by the Allen Institute. All tools described above were combined in Python using custom-written scripts (https://github.com/jenniferColonell/ecephys_spike_sorting). Finally, manual curation was performed to identify individual neurons and to remove multiunit and noise clusters using phy (https://phy.readthedocs.io/en/latest/).

## *In vivo* electrophysiology recordings

Preparation of files for cluster sorting was carried out using a custom MATLAB algorithm. Spikes were thresholded to a 6 sigma criteria and traces were aligned to their depolarization peak. Spikes were exported in a.plx file format to be imported into Offline Sorter (Plexon Inc., Dallas, TX). Once spikes were imported, principal component analysis was used to cluster spikes into individual units.

### Analysis of neural responses to cue delivery

For all electrophysiology experiments, to calculate the neuronal response to each conditioned stimulus, the mean activity for each neuron across all trials of a given trial type was calculated and z-score of changes in firing rate was calculated based on mean and standard deviation for that neuron during a 20 second baseline period immediately preceding the onset of the cue. For hierarchical clustering, each neuron’s z-score response for reward trials and fear trials were horizontally concatenated then clustered using agglomerative hierarchical clustering based on Euclidean distance with a custom MATLAB script. For ‘phasic’ and ‘sustained’ responses, the mean z-score was calculated during a period of 0-100 ms following the cue and 0.1-19s following the cue, respectively.

### Neural trajectory analysis

To visualize the population-level firing rate dynamics, population vectors containing the simultaneous activity of all recorded neurons were constructed for trajectory analysis. For each neuron, activity was averaged across trials of the same condition (CS-shock or CS-reward) to reduce noise, but the individual contribution of each neuron to the population vector was preserved throughout the analysis. A single global Principal Component Analysis (PCA) was applied to a matrix containing these population vectors with standardized neural activity (concatenated activity during shock and reward trials) from all animals (paired and unpaired). This approach enabled the comparison of neural trajectories across experimental groups while maintaining the population-level dynamics. The population vectors for each group of animals and each trial type were then projected into this principal component space and their trajectories within the first three dimensions are shown. The distance between the shock and reward trajectories for each experimental group was calculated as the Euclidean distance within each time-bin using the complete principal component space. For visualization, we smoothed each trajectory by convolving the signal with a one-directional Gaussian.

For pairwise comparison of trials, we computed cosine similarity between population activity vectors from individual trials to assess the robustness of neural representations at the single-trial level. For each trial, we subtracted the mean baseline activity (−20 s to 0 s relative to CS onset) to remove tonic firing rate differences and isolate stimulus-evoked responses. Cosine similarity was computed between baseline-subtracted population vectors at each time bin, where values of 1 indicate identical trajectory directions, 0 indicates orthogonal representations, and −1 indicates opposite directions. Three types of pairwise comparisons were computed: shock vs. shock (two different shock trials), reward vs. reward (two different reward trials), and shock vs. reward (one shock trial vs. one reward trial). To obtain robust estimates, we performed 100 iterations of random trial sampling per animal, yielding time-resolved cosine similarity traces for each comparison type. For statistical testing, cosine similarity values were averaged over the CS presentation period (0–18 s), and within-group comparisons were performed using the Wilcoxon signed-rank test to determine whether shock vs. reward representations were more dissimilar than within-condition comparisons.

### Logistic regression classifier

To test whether ASt neural activity within a trial is correlated with various behavioral variables logistic regression models were used.^59^ For each mouse separately, the neural activity of all trials was divided into five non overlapping folds to use for cross-validation. To classify whether CS-shock or CS-reward was presented during a trial, a logistic regression classifier was trained on the training folds and evaluated time-bin by time-bin on the held-out trials. Similarly, separate classifiers were trained to decode whether the mouse was engaging in freezing or darting behavior. The training data was balanced to ensure there were an equal number of time points for each class.

### Generalized linear model

To determine whether ASt neurons encode conditioned stimulus (CS) identity independently of behavioral state, we fit a series of generalized linear models with incrementally restricted predictor sets for each neuron’s activity during the CS presentation period (0–18 s following CS onset). Animals with fewer than six ASt neurons were excluded from analysis. Neurons for which any model failed to converge were also excluded. Spike times were binned into 100 ms intervals and subsampled to 500 ms effective resolution to reduce computational burden while preserving temporal structure for modeling autocorrelation. Three behavioral covariates were aligned to each time bin. Freezing and darting were scored as binary variables derived from SLEAP-detected bout onset/offset times. Locomotion speed (cm s^−1^) was computed from frame-level centroid displacement × video frame rate, averaged within each bin, and z-scored within animal to facilitate model convergence.

Spike counts were modeled using Generalized Estimating Equations (GEE)^103^ with a Poisson distribution and log link function, and first-order autoregressive (AR(1)) working correlation structure, with each trial as the clustering unit. For each neuron, four models were fit:

Full model: *log(μ) = β_0_ + β_CS_ · CS + β_Freeze_ · Freeze + β_dart_ · Dart + β_speed_ · Speed + β_time_ · Time*

Behavior-only model: *log(μ) = β_0_ + β_Freeze_ · Freeze + β_dart_ · Dart + β_speed_ · Speed + β_time_ · Time*

CS-only model: *log(μ) = β_0_ + β_CS_ · CS + β_speed_ · Speed + β_time_ · Time*

Null model: *log(μ) = β_0_*

CS is a binary indicator of tone identity (0 = reward, 1 = shock), Freeze and Dart are binary behavioral state indicators, Speed is z-scored locomotion speed, and Time is normalized time within the CS window (0– 1). Locomotion speed was included in all non-null models as a nuisance regressor so that model comparisons isolate the predictors of interest. The difference between the full and behavior-only models reflects the unique contribution of CS identity, while the difference between the full and CS-only models reflects the unique contribution of freezing and darting, with speed controlled in both cases. Neurons for which any model failed to converge were excluded from downstream analyses.

Because GEE estimates quasi-likelihood rather than true likelihood, standard likelihood ratio tests are not applicable to GEE models. We therefore assessed predictor significance using trial-level permutation tests. For the CS predictor, CS labels were shuffled across trials (held constant within each trial to preserve within-trial temporal structure, AR(1) correlation, and behavior alignment), the full model was refit, and the permuted CS coefficient was recorded. The two-sided p-value was computed as the proportion of 1,000 permutations for which *|β_CS_^perm^| ≥ |β_CS_^obs^|*. For the behavioral predictors, the triplet of time-varying covariates (freezing, darting, and speed) was jointly permuted across trials, reassigning which trial’s behavioral trajectory paired with which trial’s neural data. Significance was assessed using the joint statistic *β^2^_freeze_ + β^2^_dart_ + β^2^_speed_* (one-sided). This trial-level permutation approach preserves the within-trial autocorrelation structure, which standard coefficient-level tests do not account for.^104^ Permutation p-values were corrected for multiple comparisons across neurons within each group using the Benjamini–Hochberg procedure (*α* = 0.05).

To assess model generalizability, we performed five-fold trial-stratified cross-validation for each neuron. In each fold, the full, behavior-only, and CS-only models were fit on training trials and evaluated on held-out trials using Poisson deviance *D = 2Σ[y log(y/μ◻) − (y − μ◻)]*, where *μ◻ = exp(X_test_ · β_train_)*. The cross-validated improvement attributable to CS was quantified as *ΔDeviance_CS_ = Deviance(behavior − only) − Deviance(full)* on held-out data. Positive values indicate that CS identity improves out-of-sample prediction beyond behavioral covariates alone.

For between-group comparisons, we computed per-animal medians of each metric and compared paired and unpaired groups using one-sided Mann–Whitney U tests, treating animals as the independent unit to avoid pseudoreplication. We additionally report Fisher’s exact test on the neuron-level proportion of CS-responsive neurons, noting that neurons within animals are not independent. All analyses were performed in Python 3.11 using statsmodels (v0.14+) for GEE model fitting.

## Statistical analysis of calcium recordings

Imaging data was then concatenated and temporally downsampled by a factor of two using a custom MATLAB script before motion correction (rigid registration) via the NoRMCorre algorithm.^105^ Neuron detection and signal extraction was done using the CNMF-E algorithm.^106^ Using a MATLAB Neuron Deletion GUI, neurons exhibiting abnormalities in morphology and calcium trace were manually excluded. Neuron curation was performed by experimenters blinded to the experimental condition.

To calculate the neuronal response to each conditioned stimulus, the mean GCaMP7f fluorescence for each neuron across trials for each trial type was calculated, and z-score of fluorescence values calculated based on mean and standard deviation for that neuron during a 10 second baseline period immediately preceding the onset of the cue. Responses to each CS were quantified over a short (0-1 s) and long (1-10 s) window following cue onset, analogous to the phasic and sustained windows used for the electrophysiology data. For clustering, each neuron’s individual z-score response for CS-shock trials and CS-reward trials were horizontally concatenated using a custom MATLAB script and clustered using agglomerative hierarchical clustering based on Euclidean distance, with soft normalization of neurons to a maximum z-score of 10. For the US-only experiment, z-scores were calculated for each neuron based on a 5 second baseline period immediately preceding shock delivery or consumption of reward.

## Statistical analysis of RNAscope in situ hybridization

### Object identification and analysis

Images were analyzed in CellProfiler using a version of the CellProfiler Colocalization pipeline (https://cellprofiler.org/examples/) optimized for each experiment. The pipeline identified the ROIs of DAPI labeled cells and then subsequently identified mRNA puncta associated with those ROIs to allow for quantification of RNA expression per cell. Due to varying image sizes and labeling intensities, different criteria were used for puncta detection in each experiment (*Drd1a/Drd2*, ≥6 puncta, *Cdh6/Slit2* experiment, ≥3 puncta, *VGlut1* experiment: ≥1 puncta). However, within each experiment, analysis settings were kept constant to allow for direct quantitative comparison of RNA labelling across different cells and regions. Total number and density of RNA-labeled cells in each region of interest were calculated from CellProfiler.csv outputs using custom MATLAB code.

### Comparison of Drd1a/Drd2 composition across regions

To compare the *Drd1a/Drd2* balance between striatal regions, the animal, rather than the individual cell, was treated as the unit of analysis. Cells exceeding the puncta threshold (above) for a single probe were classified as *Drd1a*-only or *Drd2*-only, and those exceeding it for both as dual-labeled. For each animal and region, the balance was summarized as the log-ratio log(*Drd1a*-only / *Drd2*-only), which excludes dual-labeled and unlabeled cells. Regions were compared pairwise (ASt vs DS, ASt vs TS, DS vs TS) with two-tailed paired t-tests on these per-animal log-ratios, Holm-Bonferroni corrected across the three comparisons.

## Statistical analysis of single-nucleus RNA sequencing

### Sequence alignment

All samples were processed using CellRanger (v5.0.0).^50^ All processing was done by using CellRanger’s implementation of STAR to align sample sequence reads to their pre-built mm10 vm23/Ens98 reference transcriptome index 2020-A, with predicted and non-validated transcripts removed. All sequencing reads were aligned to both the exons and the introns present in the index. Samples were demultiplexed to produce a pair of FASTQ files for each sample. FASTQ files containing raw read sequence information were aligned to the CellRanger index using the cellranger count command with --chemistry SC3Pv3 and --include-introns flags enabled. CellRanger corrected sequencing errors in cell barcodes to pre-defined sequences in the 10X v3 single-index whitelist within Hamming distance 1. PCR duplicates were removed by selecting unique combinations of corrected cell barcodes, unique molecular identifiers, gene names, and location within the transcript. Raw unfiltered count data was read into R (v4.0.3) using the Seurat package (v4.0).^107–110^ The final result of the pipeline was a barcode x gene expression matrix for further analysis downstream.

### Quality control

We used the raw, unfiltered matrix output from CellRanger as the input to the beginning of the pipeline. However, to apply a more stringent filter, the emptyDrops dirichlet-multinomial model from the DropletUtils package (v1.10.2) was applied to each library individually.^111,112^ Droplets with less than 100 total counts were used to construct the ambient RNA profile and an FDR threshold below 0.001 was used to select putatively occupied droplets. All barcodes with greater than 1000 UMIs were further assumed non-empty. Most quality filtration choices were heavily influenced by the recommendations presented in pipeComp.^113^ All quality control was performed on each library individually prior to merging. Minimal quality filtering for each barcode was performed by setting a floor of 1000 features per barcode for downstream inclusion to ensure the dataset is entirely composed of high-quality nuclei. Next, to remove highly likely multiplet barcodes, barcodes were filtered out if their count depth was more than 5 median absolute deviations above the median count depth. Barcodes were then removed if their proportion of ribosomal or mitochondrial reads was more than 5 interquartile ranges above the 75th percentile. Heterotypic doublets were identified by creating simulated artificial doublets in scDblFinder (v1.4.1), which uses a DoubletFinder-like model to remove barcodes similar to simulated doublets, with an assumed doublet rate of 1% per 1000 nuclei in the library.^114,115^ Scater (v1.18.3) was used to produce initial diagnostic tSNE and UMAP plots for visually checking the influence of each above metric on the structure of the data.^116^ Downstream, one cluster of diffusely-distributed neurons with few marker genes and disproportionately low read depth and disproportionately high mitochondrial and ribosomal gene proportions was removed as low-quality cells.

### Data processing/transformation

All datasets (initially for all nuclei and again for selected subclusters) were formatted into Seurat objects (v4.0.0), merged, and then normalized and transformed individually using the SCTransform (v2) variance stabilizing transform, which performs best according to prior comparisons in pipeComp.^113,117,118^ Following the merge, all genes expressed in 3 or fewer cells were removed from analysis. SCTransform was run returning Pearson residuals regressing out mitochondrial gene expression, first with 5000 highly variable features, and then with 3000 for subsequent iterations on subsets of the data. Dimensionality of the dataset was first reduced using principal component analysis, as implemented in Seurat’s RunPCA function.^119^ The top 50 principal components were retained for downstream analysis. These principal components (top 25 for all nuclei, top 15 for subclusters) were used as input to the non-linear tSNE and UMAP dimensionality reduction methods as implemented by Seurat’s RunTSNE and/or RunUMAP functions (n.epochs = 1000, min.dist = 0.5) at default settings unless otherwise specified.^120–122^

### Differential expression

Marker genes were identified using Wilcoxon rank-sum test as implemented by the FindConservedMarkers function in Seurat, using the region as a grouping variable. Genes were accepted as differentially expressed with a minimum proportion cutoff at 0.1 and minimum fold change at 1.5-fold (log2-fold change of 0.585), with a p-value cutoff of 0.01 after Bonferroni correction. To identify genes differentially expressed by region, single-cell values were converted to pseudo-bulk by batch using the run_de function as implemented in the Libra package (v1.0.0) using default settings, and tested for differential expression using edgeR’s likelihood ratio test.^123,124^ Region-specific genes were identified by comparing batches from one region to all others, while pairwise differentially-expressed genes were identified between batches from just the two regions of interest, and genes were retained with a minimum fold change at 1.5-fold (log2-fold change of 0.585), at a p-value cutoff of 0.01 after Benjamini-Hochberg correction. Neurologically-related genes of interest were determined by filtering region-specific differentially expressed genes through a subset of pre-selected gene families from HGNC with known functions related to neural structure and function.^125^

## SUPPLEMENTARY FIGURES

**Figure S1.**
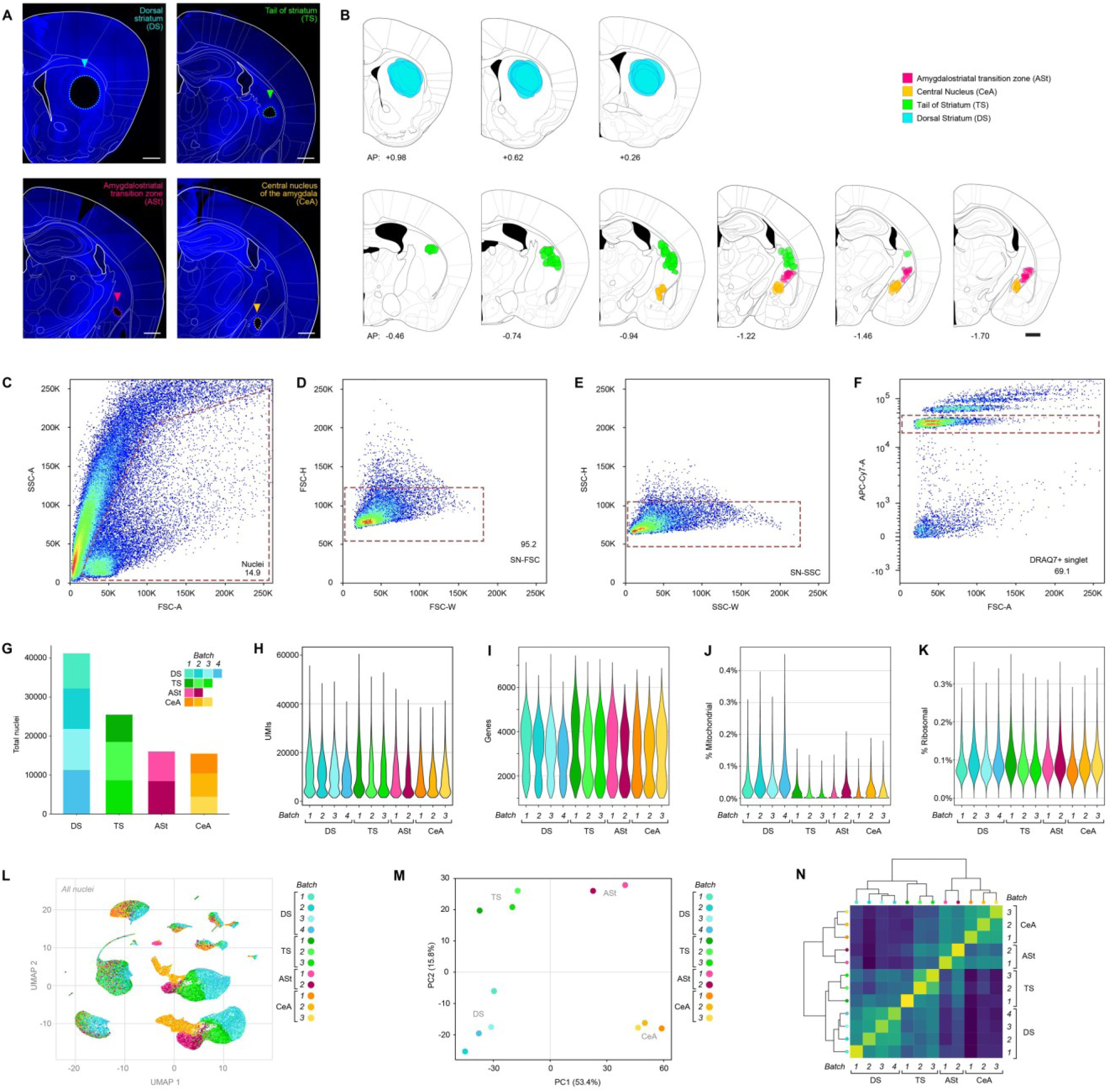
Quality control metrics for single-nucleus RNA sequencing. (**A**) Representative images of tissue microdissection sites from RNA sequencing target regions, the dorsal striatum (DS), tail of striatum (TS), amygdalostriatal transition zone (ASt), and central nucleus of the amygdala (CeA) (Blue = DAPI, scale bars = 500 µm). (**B**) Location of all tissue sample sites used for single-nucleus RNA sequencing, color coded by region. AP = anteroposterior distance from bregma (mm), scale bar = 500 µm. (**C-F**) Density plots outlining the gating strategy for FANS isolation of single nuclei. Nuclei isolated from regions of interest were FANS sorted using 70 μm nozzle at 52 psi. Nuclei were sorted based on size (**C**), duplicates and/or morphology (**D, E**), and by high DRAQ7 signal, which stains DNA in nuclei (**F**). Single DRAQ7⁺ events at the lowest stoichiometric fluorescence multiple were considered nuclei. (**G**) Absolute number and proportion of nuclei passing quality control filters from each batch in each region. (**H**) UMIs detected per nucleus, filtered at the median per library + five times the median absolute deviation. (**I**) Genes detected per nucleus, filtered at minimum 1000 genes. (**J**) Percent mitochondrial reads per nucleus, filtered at median per library + five times the median absolute deviation. (**K**) Percent ribosomal reads per nucleus, no quality filter applied. (**L**) UMAP of all sequenced nuclei colored by both target region and batch identity. (**M**) PCA of pseudobulk samples created from each batch, colored by both target region and batch identity. (**N**) Evaluation of transcriptional homology on a per-batch basis, where the distance matrix is based on Spearman correlation between median expression of genes on a per-region basis, and the dendrogram was created via hierarchical clustering on this correlation matrix.

**Figure S2.**
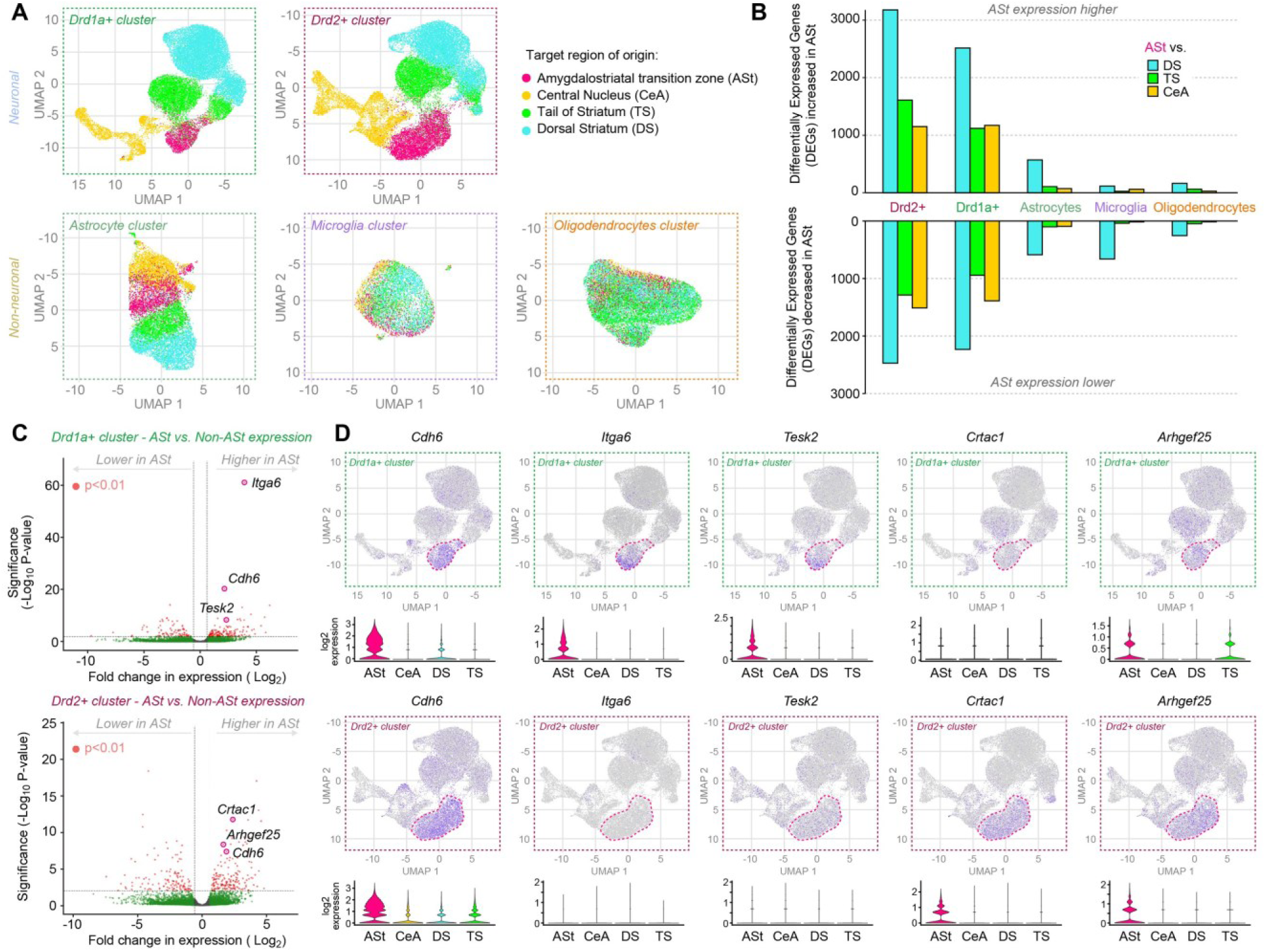
Single-nucleus RNA sequencing identifies unique transcriptomic signatures for ASt cells. (**A**) UMAP projections of nuclei from the largest cell type clusters (*Drd1a+*, *Drd2+*, Astrocytes, Microglia and Oligodendrocytes) colored by tissue region of origin. (**B**) Number of differentially expressed genes (*p* < 0.01, 1.5 log-fold change) with increased expression (top) or decreased expression (bottom) in ASt nuclei relative to nuclei from the central nucleus of the amygdala (CeA), tail of striatum (TS), and dorsal striatum (DS) within each cell type cluster. (**C**) Volcano plot showing differentially expressed genes between ASt and non-ASt nuclei in *Drd1a+* cluster (top) and *Drd2+* cluster (bottom). Dashed lines indicate *p* < 0.01 and a fold-change of 1.5. Dots colored grey fail to meet either standard, dots colored green have sufficient fold change but not significance, while red dots meet both standards. Marker genes of interest for further analysis are circled. (**D**) Expression of specific marker genes of interest in *Drd1a+* cluster (top) and *Drd2+* cluster (bottom) neurons. Expression is visualized via UMAP and violin plot. UMAP expression is colored based on increasing normalized expression intensity. Violin plots show smoothed expression density, colored and split based on regional identity within each cluster.

**Figure S3.**
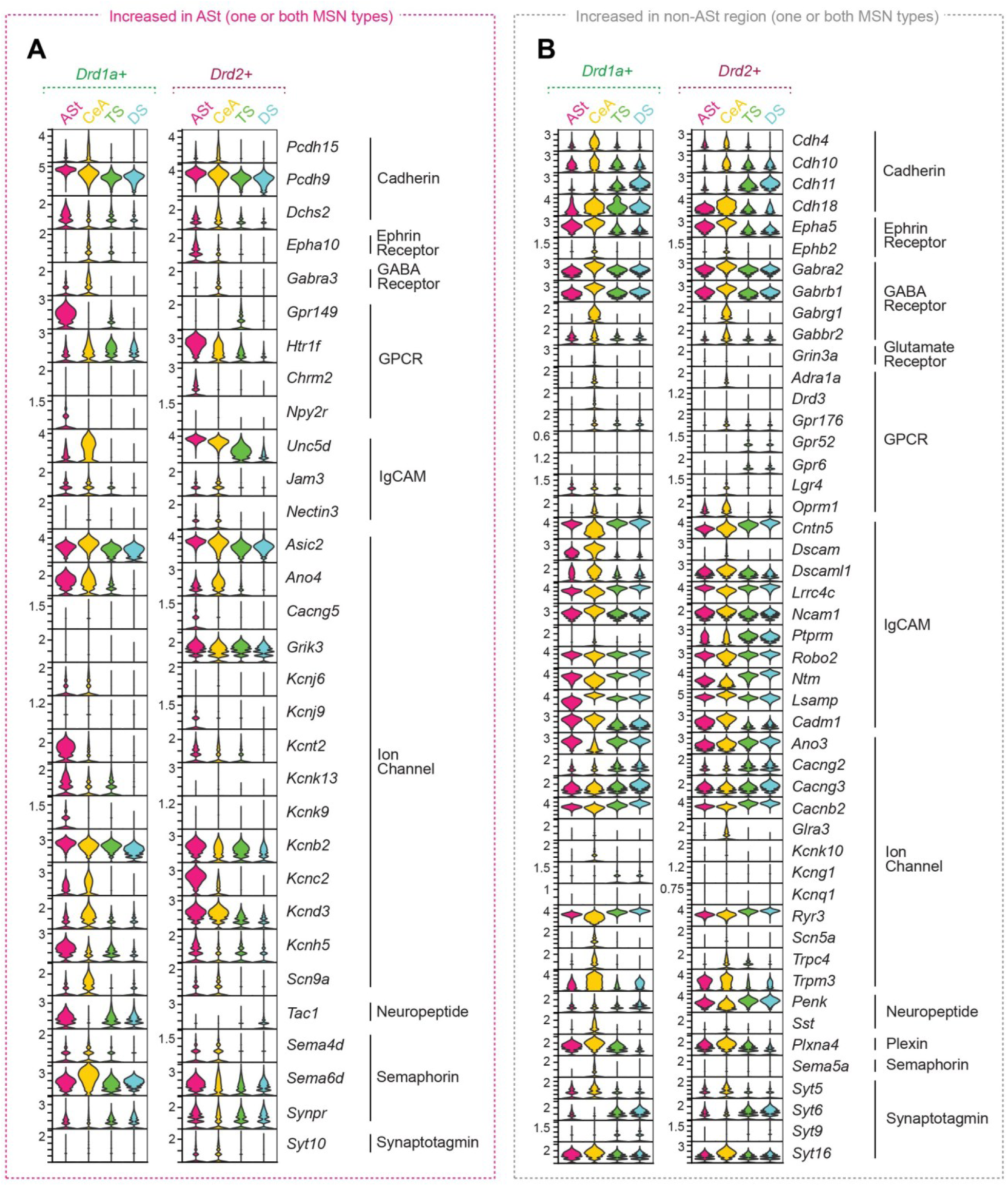
Differentially expressed genes of interest with neurologically-relevant function in striatal subregions of interest. (**A**) Differentially expressed genes with increased expression in ASt falling into predefined HGNC gene families related to neurological function, increased in *Drd1a+* and/or *Drd2+* MSNs. (**B**) Differentially expressed genes with increased expression in batched non-ASt regions (DS, TS, and/or CeA) falling into predefined HGNC gene families related to neurological function, increased in *Drd1a+* and/or *Drd2+* MSNs.

**Figure S4.**
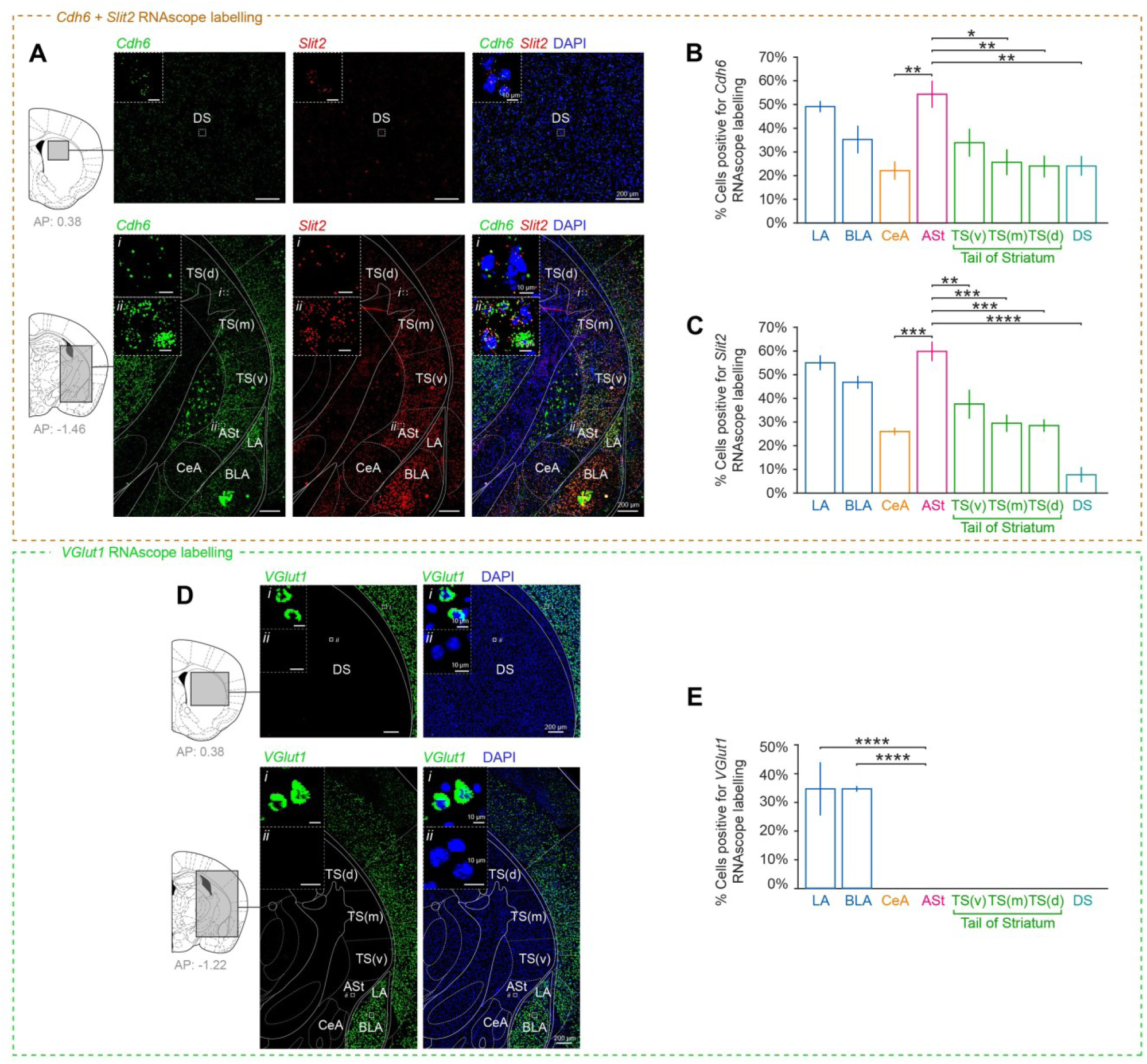
*Cdh6* and *Slit2* are enriched and VGlut1 is absent in the Ast. (**A**) Representative images of *in situ* RNAscope labeling of *Cdh6* RNA (green) and *Slit2* RNA (red) in striatal target regions. For analysis of dorsoventral expression gradients, the tail of striatum was spatially separated into three segments (TS(v), TS(m), TS(d)), each ∼500 µm in length along the DV axis. (**B**) Relative proportion of cells in each target region positively labeled for *Cdh6* RNA. (*n =* 3 mice, One-way ANOVA, F(7, 16) = 6.320, p = 0.0011). (**C**) Relative proportion of cells in each target region positively labeled for *Slit2* RNA (*n =* 3 mice, One-way ANOVA, F(7, 16) = 22.65, p = 3.6 × 10⁻ ⁷). Consistent with our sequencing data, *Cdh6* and *Slit2* showed significantly greater expression in the ASt compared with more dorsal regions of the tail of striatum (TS). *Cdh6* and *Slit2* expression was also greater in the ASt than the CeA, while *Cdh6* and *Slit2* expression in the ASt and LA was similar, consistent with the LA and ASt receiving highly similar axonal inputs ^25,37^. (**D**) Representative images of *in situ* RNAscope labeling of *VGlut1* RNA (green). *VGlut1* expression was not seen in the ASt, indicating the small number of *VGlut1*-expressing nuclei detected in ASt sequencing samples (see **Fig 1, E-G)** were attributable to small inclusions of tissue from adjacent regions with glutamatergic neurons. (**E**) Relative proportion of cells in each target region positively labeled for *VGlut1* RNA. No *VGlut1*-expressing (excitatory) neurons were identified in the ASt, the CeA, or other striatal subregions. (*n =* 3 mice, One-way ANOVA, F(7, 16) = 24.37, p = 2.1 × 10⁻ ⁷). All data presented as mean ±SEM. Post-hoc comparisons between regions, Tukey’s test, *\*p* < 0.05, \**\*p* < 0.01, \*\**\*p* < 0.001, \*\*\**\*p* < 0.0001. For clarity, only significant pairwise comparisons with ASt are shown.

**Figure S5.**
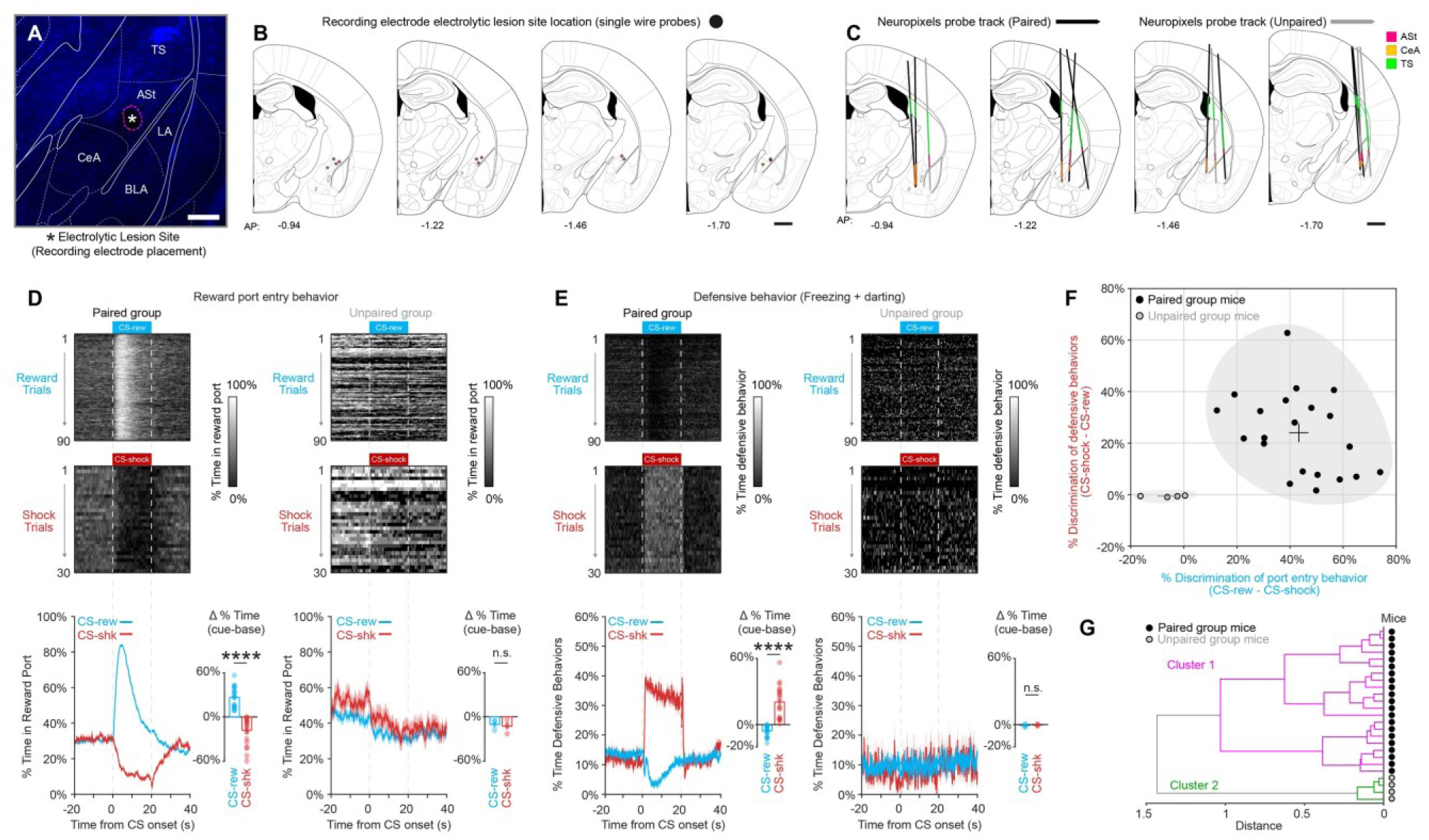
Histological targeting and behavioral validation for *in vivo* electrophysiology recordings. (**A**) Representative image of electrolytic lesion site to confirm recording electrode targeting to the ASt (Blue = DAPI, Scale bar = 100 μm). (**B**) Histologically verified lesion sites of single-wire electrode placements in recording experiments. (**C**) Histologically verified probe tracks of Neuropixels probes in recording experiments. Colors indicate segment of probe that was in a target region analyzed (Pink = ASt, Green = TS, Orange = CeA). (**D-G**) Validation of conditioned behavioral responses during two-tone discrimination task in mice in electrophysiological recording experiments (Paired: n = 21 mice, Unpaired: n = 4 mice). Paired group mice showed distinct responses to the CS-shock and CS-reward tones in reward port entry behavior (Two-tailed paired t-test, t20 = 12.469, ****p < 0.0001) (**D**) and defensive responses (Two-tailed paired t-test, t20 = 6.711, ****p < 0.0001) (**E**), consistent with successful discrimination between the two tones. Neither measure differed between cue types in unpaired mice (port entry, t3 = 0.672, p = 0.550; defensive responses, t3 = 0.658, p = 0.557). (**F**) ‘Discrimination scores’ of reward port entry behavior (CS-rew − CS-shock) plotted against defensive behavior (CS-shock − CS-rew) for each mouse. Crosses indicate group mean. (**G**) Agglomerative hierarchical clustering of electrophysiology mice based solely on their defensive and reward behavior discrimination identified unpaired mice as a distinct cluster relative to all paired group mice.

**Figure S6.**
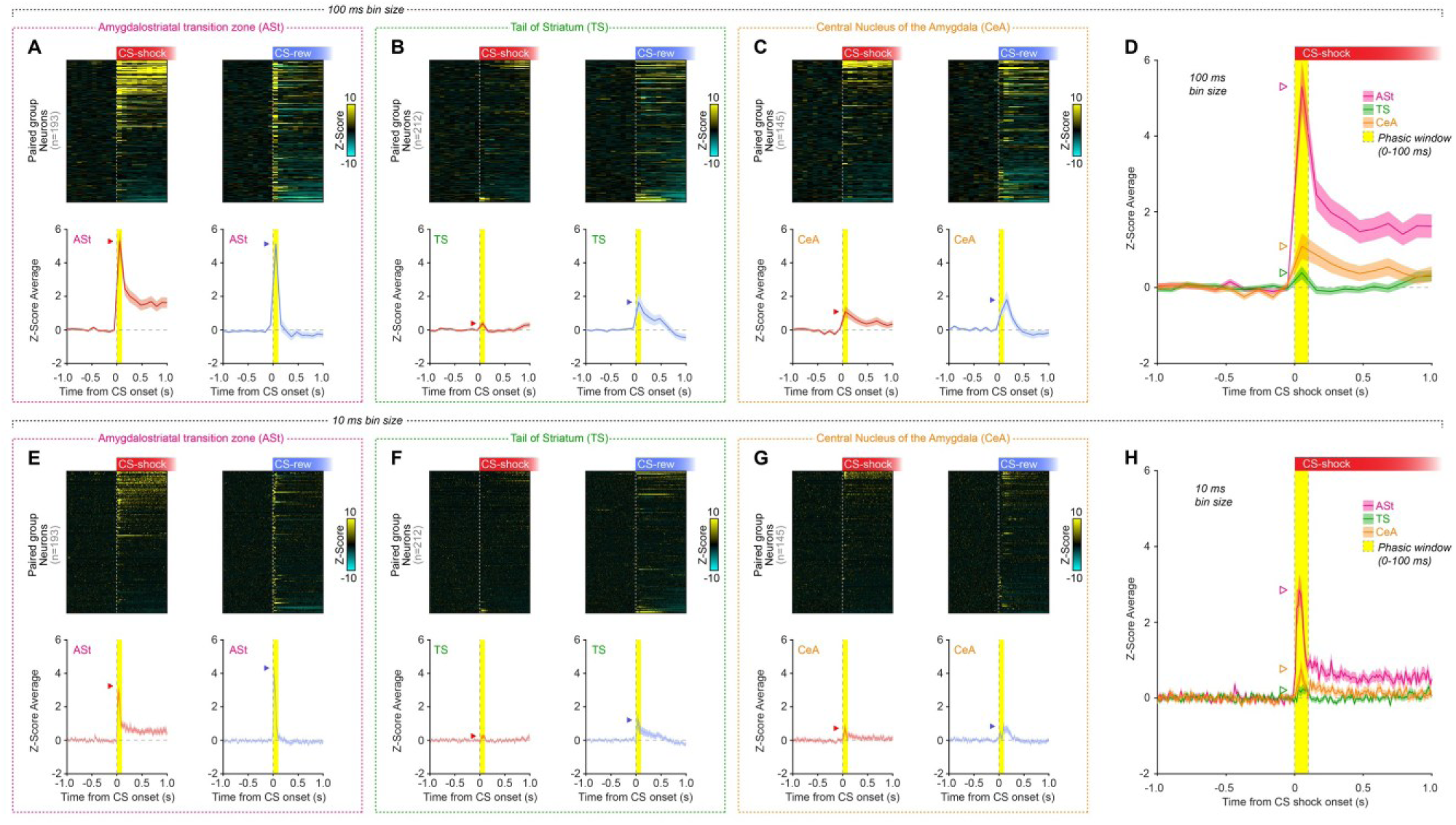
Phasic responses to CS-shock and CS-reward in the ASt, TS, and CeA. (**A-C**) Heat maps and group average of z-score changes in neural firing rate in response to CS-shock and CS-reward in ASt (**A**), TS (**B**), and CeA (**C**). Data shown here are a zoomed-in visualization of data in Figure 2F,G,H. (**D**) Group average z-score changes in neural firing rate to CS-shock in ASt, TS, and CeA, plotted together for easier comparison. Heat maps and group average of z-score changes in neural firing rate, with 10 millisecond bins, in response to CS-shock and CS-reward in ASt (**E**), TS (**F**), and CeA (**G**). Data shown here are a zoomed-in visualization of data in Figure 2F,G,H. (**H**) Group average z-score changes in neural firing rate, with 10 millisecond bins, to CS-shock in ASt, TS, and CeA, plotted together for easier comparison. Heat map rows represent average z-score PSTH for individual neurons. Arrowheads show average peak z-score in the ‘phasic’ window (0-100 ms from CS onset). Data shown as mean ±SEM.

**Figure S7.**
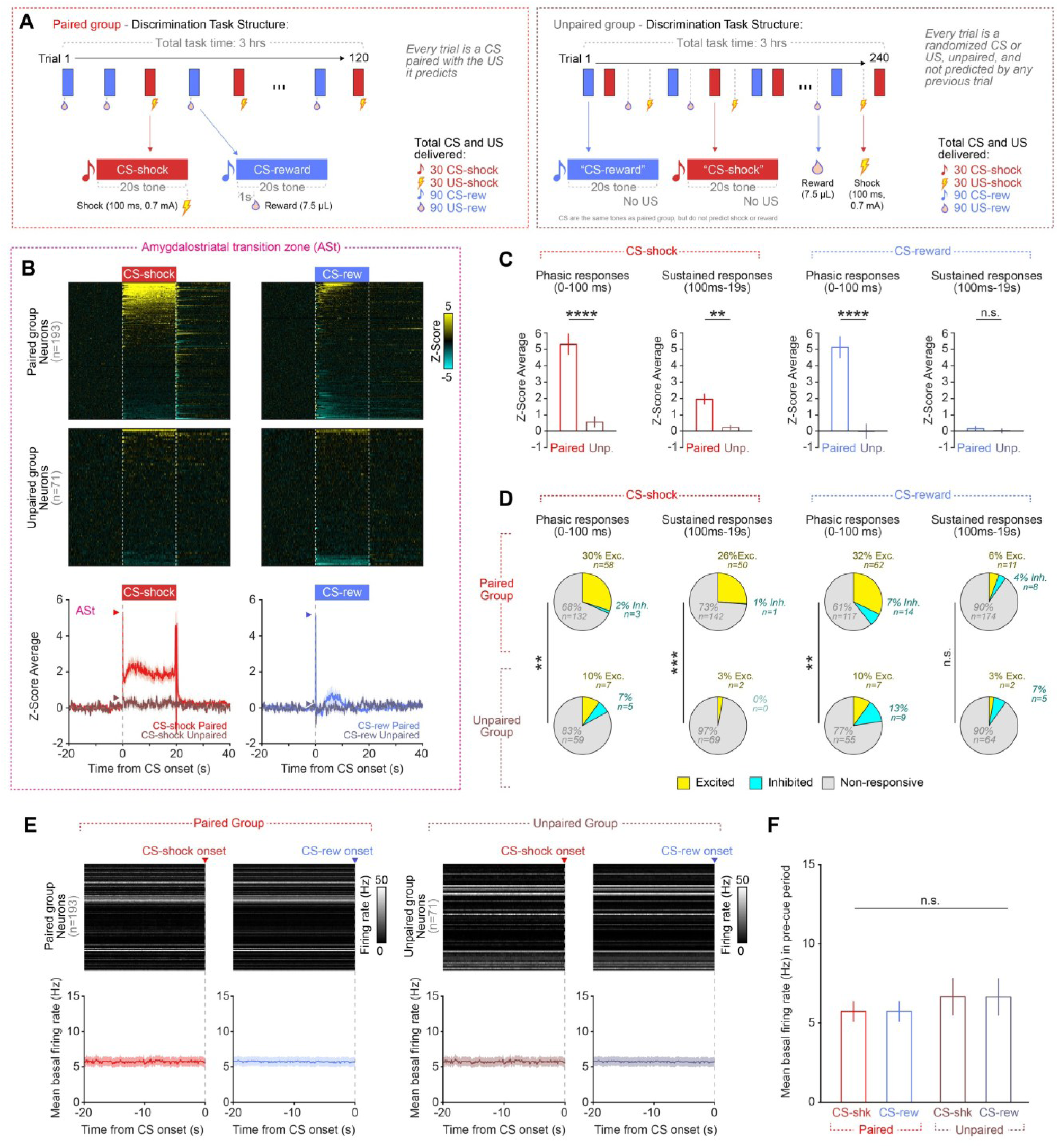
Differences in ASt neuron responses to stimuli in paired and unpaired controls. (**A**) Design of two-tone discrimination task for paired and unpaired groups in *in vivo* electrophysiology recordings. (**B**) Heat maps and group average of z-score changes in neural firing rate in response to CS-shock and CS-reward in ASt paired and unpaired groups. Heatmap rows represent average z-score PSTH for individual neurons. Arrowheads show average peak z-score during phasic window (0-100 ms). (**C**) ASt neurons in paired group mice showed greater phasic responses to CS-shock (two-tailed t-test, t262 = 4.332, ****p < 0.0001), sustained responses to CS-shock (t262 = 3.109, **p = 0.0021), and phasic responses to CS-reward (t262 = 4.511, ****p < 0.0001) than unpaired controls. Sustained responses to CS-reward did not differ between groups (t262 = 0.460, p = 0.646). (**D**) Proportions of ASt neurons in the paired and unpaired groups that were significantly excited or inhibited by CS-shock or CS-reward during the phasic or sustained windows (chi-square test with Holm-Bonferroni correction; CS-shock phasic, χ2(2, N = 264) = 15.31, **p = 0.0014; CS-shock sustained, χ2(2, N = 264) = 18.04, ***p = 0.0005; CS-reward phasic, χ2(2, N = 264) = 13.86, **p = 0.0020; CS-reward sustained, χ2(2, N = 264) = 1.76, p = 0.415). **p < 0.01, ***p < 0.001. (**E**) Heat maps and group average of basal neural firing rates in the 20s window prior to CS onset, for paired and unpaired group ASt neurons. (**F**) No significant differences in basal firing rate were seen between paired and unpaired group ASt neurons, or between CS-shock and CS-reward windows (two-way mixed ANOVA, group [between-neuron] × cue window [within-neuron]: group main effect F(1,262)=0.51, p=0.476; cue main effect F(1,262)=0.01, p=0.926; group × cue interaction F(1,262)=0.12, p=0.728). Paired: n=15 mice, 193 neurons; Unpaired: n=4 mice, 71 neurons.

**Figure S8.**
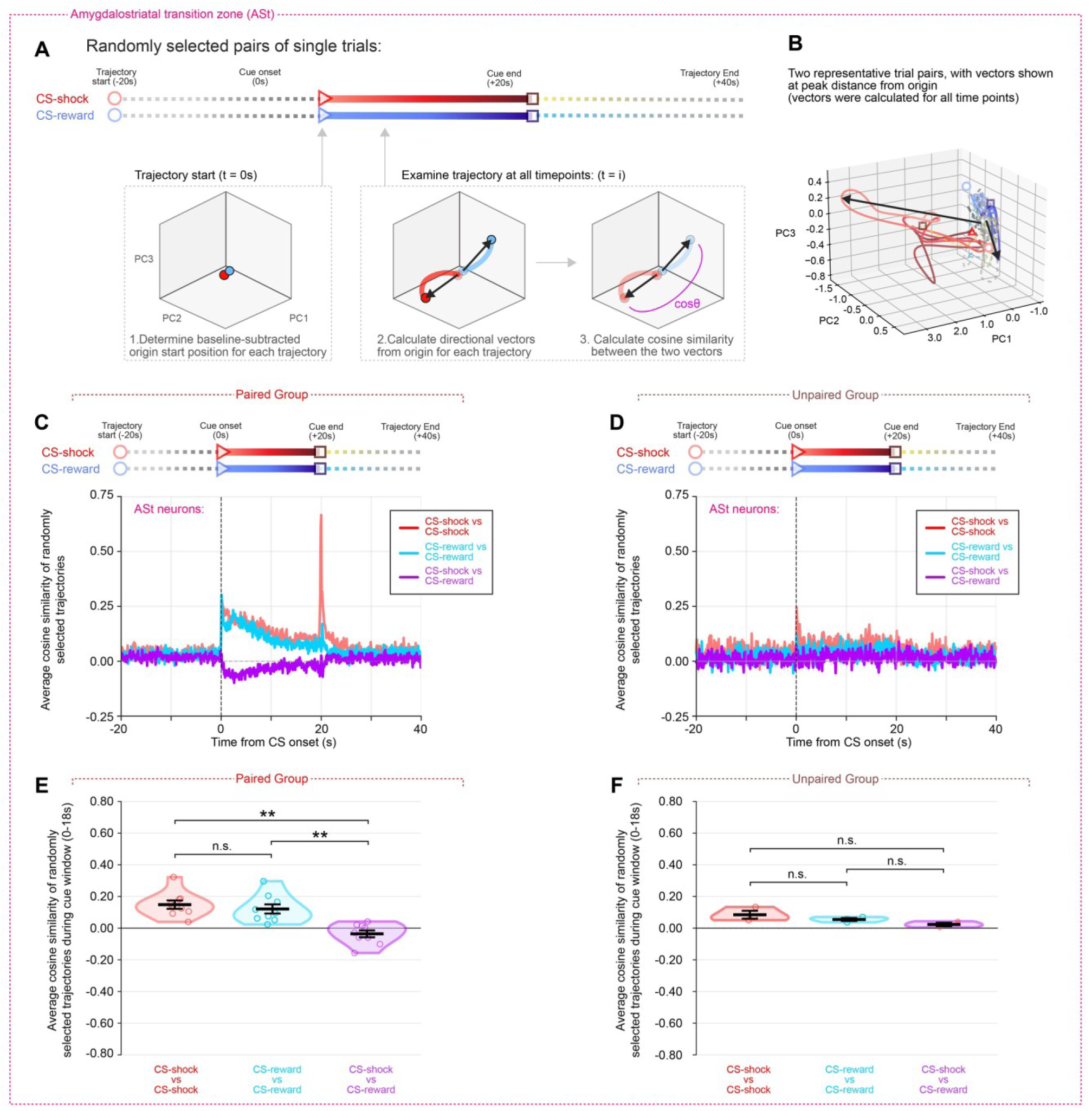
Single-trial population-trajectory analysis using randomly sampled trials demonstrates divergence of ASt population dynamics in CS-shock and CS-reward trials. (**A**) Schematic of the trajectory-similarity method. Trial pairs were randomly selected, and average baseline (-20s to 0s) subtracted to determine normalized trajectory origins. For all subsequent timepoints, the instantaneous trajectory direction was calculated as a vector from the origin, and cosine similarity of the two trajectories calculated. (**B**) Example neural population trajectories for two representative CS-shock (red) and CS-reward (blue) trial pairs. For visualization, arrows indicate the trajectory direction at the time of peak distance from approximate baseline average origin. Similar calculations were performed at all timepoints across the full trajectory. (**C-D)** Average cosine similarity of randomly selected trajectory pairs across time in paired group (**C**) and unpaired group (**D**) mice. Same-CS trajectory pairs (CS-shock vs CS-shock; CS-reward vs CS-reward) show higher similarity following cue presentation, whereas opposite-CS trajectory pairs (CS-shock vs CS-reward) rapidly diverge in paired group, producing markedly lower similarity during and after the cue. In unpaired group mice, the negative divergence of CS-shock and CS-reward trial pairs was not seen. (**E-F**) Summary of mean cosine similarity in the cue window (excluding shock delivery at end of CS) in paired group (**E**) and unpaired group (**F**) mice. Paired animals show significantly reduced similarity between CS-shock and CS-reward trajectories reflecting the emergence of separable, CS-specific neural population dynamics in ASt (** p < 0.01 Wilcoxon signed-rank test). Data shown as mean ±SEM.

**Figure S9.**
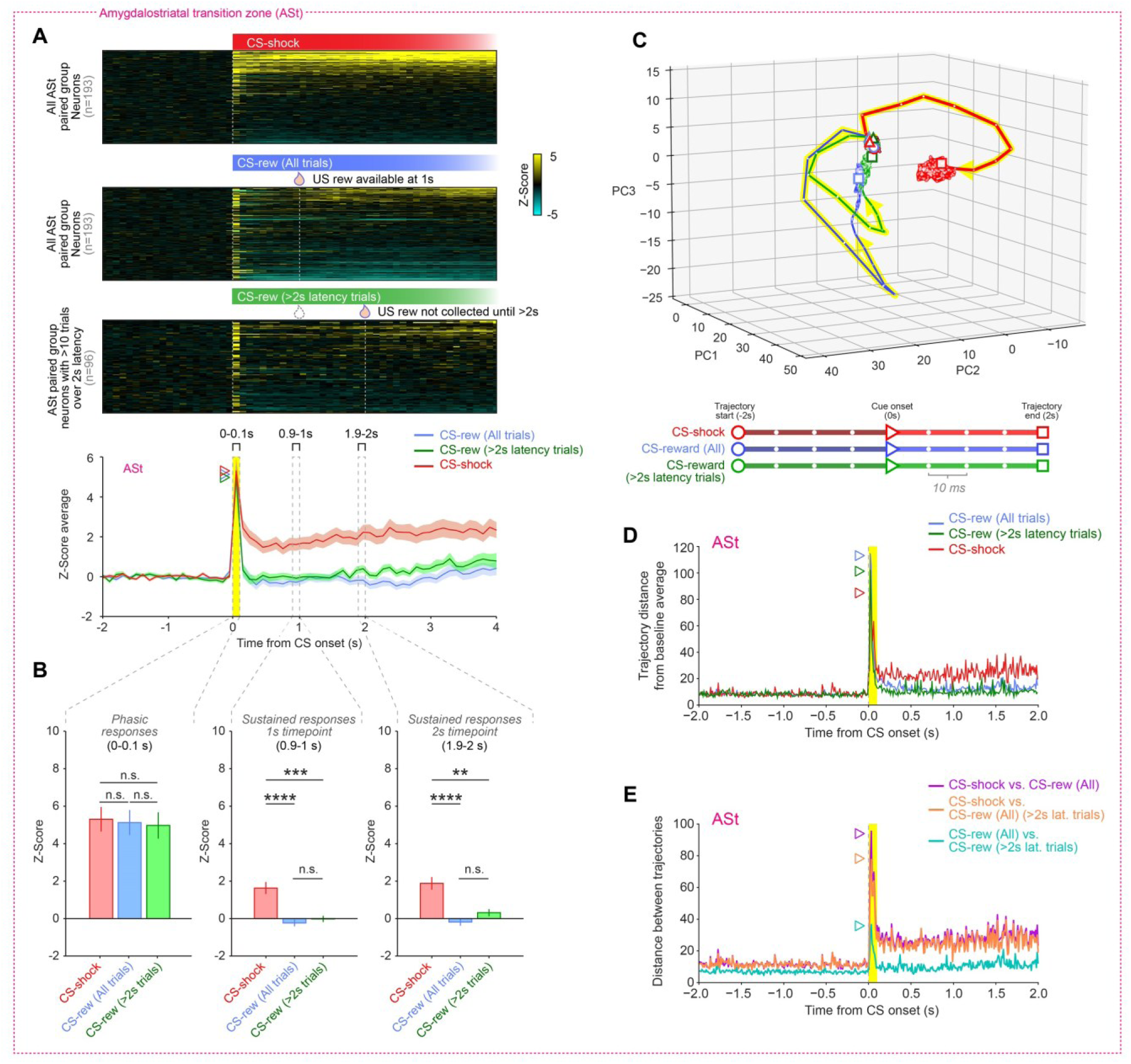
ASt neuron sustained activity is seen in response to CS-shock, but not CS-reward cues. (**A**) Heat maps and group average of z-score changes in ASt neuron firing rate in response to CS-shock, CS-reward in ASt, and the subset of CS-reward trials in which the US reward was not collected until at least 2s (termed “>2s latency trials”). Arrowheads show average peak z-score during phasic window (0-100 ms). (**B**) Mean z-score responses of ASt neurons in response to CS-shock, CS-reward and CS-reward trials with >2s latency to reward collection during the phasic and sustained time windows. No significant difference in phasic responses (0-100 ms) were observed between groups. At the 1s timepoint and the 2s timepoint, sustained responses to CS-shock were present, but these were absent in the CS-reward as well as the CS-reward trials where the latency to US reward collection was >2s. Consequently, the sustained responses were only observed in response to CS-shock, and not CS-reward, including in the portion of CS-reward trials free of any US reward responses (One-way ANOVA: phasic, F(2,479) = 0.05, p = 0.95; sustained 1s timepoint, F(2,479) = 17.38, p = 5.2 × 10⁻ ⁸; sustained 2s timepoint, F(2,479) = 17.40, p = 5.1 × 10⁻ ⁸; followed by Tukey’s multiple comparison test, **p < 0.01, ***p < 0.001, ****p < 0.0001). (**C**) Neural trajectories of neuron firing rates (10 ms bins) in the ASt for the - 2s to 2s window around onset of CS-shock, CS-reward (All), and CS-reward >2s latency trials show that CS-shock trajectories move in a distinct direction from the CS-reward trajectories. Rearward-looking gaussian (σ = 2.2) smoothing with a 50 ms max distance was applied to each trajectory. (**D**) Euclidean distance of CS-shock, CS-reward (All), and CS-reward >2s trajectories from the baseline average position prior to the CS. (**E**) Pairwise Euclidean distance of CS-shock, CS-reward (All), and CS-reward >2s trajectories from each other. The two reward trials are both highly distant from CS-shock, with sharp divergence of the trajectories immediately after CS onset, which is incompatible with a ‘salience’ response. CS-shock and CS-reward: *n =* 15 mice, 193 neurons, CS-reward >2s latency trials: *n =* 9 mice, 96 neurons. Data presented as mean ±SEM

**Figure S10.**
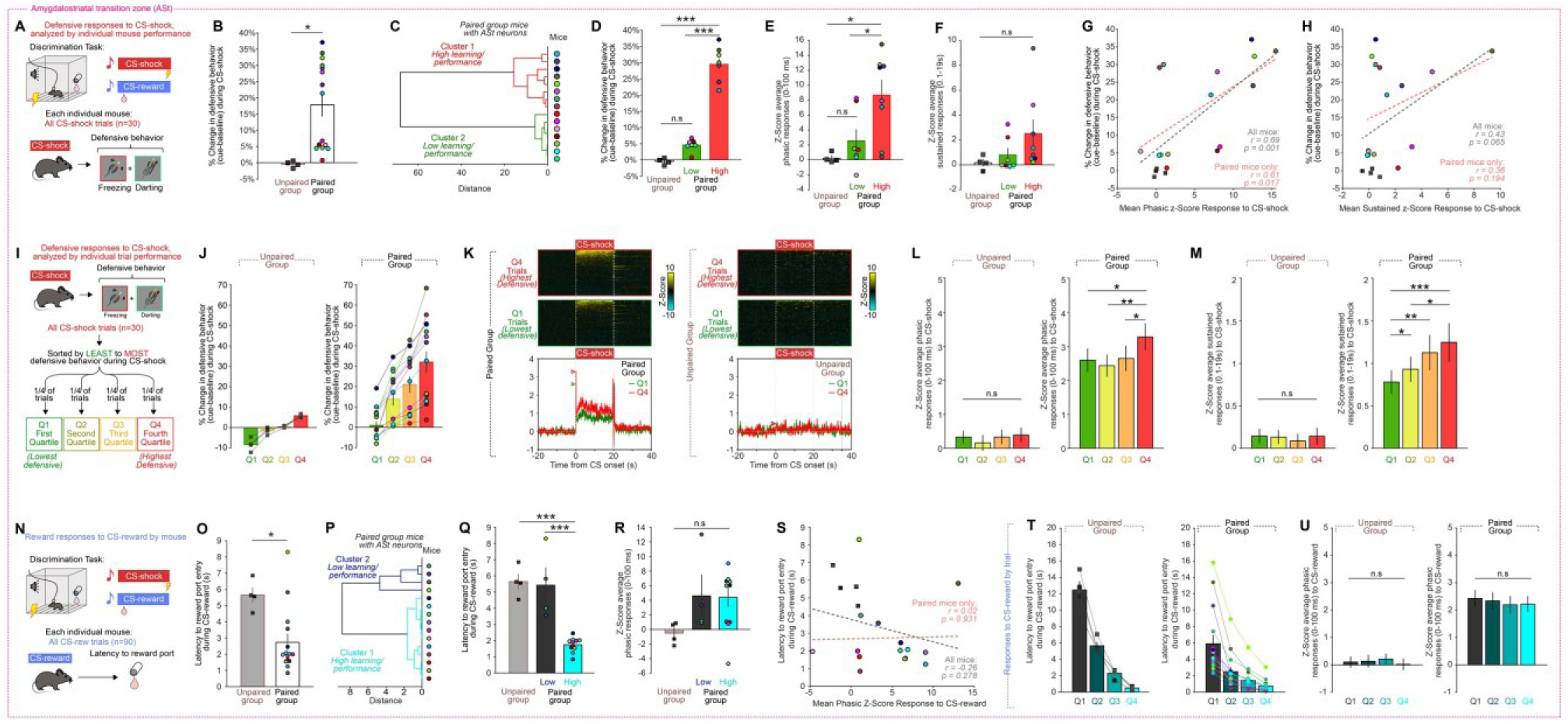
ASt neuron responses to CS-shock are correlated with increased defensive behavior across individual mice and trials. (**A-H**) Analysis of ASt neuron responses to CS-shock and defensive behaviors in individual mice (**A**) Two-tone discrimination task with electrophysiology recordings. Defensive behavior was calculated as the sum of freezing and darting behavior during each CS-shock trial. (**B**) Group average and individual mouse defensive behaviors during CS-shock was higher in paired group mice than unpaired group mice (*t17* = 2.671, *p* = 0.0161, Student’s t-test. Paired group: *n =* 15 mice, Unpaired group: *n =* 4 mice). (**C**) Hierarchical clustering of paired group mice into “low” and “high” learner/performer clusters based on magnitude of defensive behavioral responses. (**D**) Group average defensive behaviors was significantly higher in “high” performance mice than “low” performance and unpaired mice. (One-way ANOVA, F(2,16) = 130.6, *p* < 0.0001. \*\**\*p* < 0.001, Tukey’s multiple comparison test. For panels D-F, Unpaired Group: *n =* 4 mice, “Low” paired group: *n* = 7 mice, “High” paired group: *n =* 8 mice). (**E**) Average ASt neuron phasic responses to CS-shock were significantly greater in the “high” paired group than the “low” group or unpaired controls (One-way ANOVA, F(2,16) = 6.023, *p* = 0.011. *\*p* < 0.05, Tukey’s multiple comparison test). (**F**) Average ASt neuron sustained responses to CS-shock were over 3-fold greater in the “high” paired group than the “low” group, though not significant due to greater variability in sampling of sustained-responding neurons in individual mice (One-way ANOVA, F(2,16) = 1.793, *p* = 0.198). (**G-H**) Average ASt neuron phasic responses (**G**) and sustained responses (**H**) were broadly correlated with increased defensive behavioral responses to CS-shock. (**I-M**) Analysis of ASt neuron activity within mice, across trials with low or high defensive behaviors. (**I**) Separation of CS-shock trials into quartiles from highest to lowest defensive behavior. (**J**) Average defensive behaviors during CS-shock for each quartile in unpaired and paired groups. (**K**) Heat maps (top) and group average traces of ASt neuron z-score changes (bottom) in neural firing rate in response to CS-shock in Q1 and Q4 trials of paired and unpaired group mice. Heatmap rows represent average z-score PSTH for individual neurons. Arrowheads show average peak z-score during phasic window (0-100 ms). (**L**) Average ASt neuron phasic responses were greater in behavioral quartiles with more defensive behavior in paired mice, but not unpaired controls (Repeated measures ANOVA, Unpaired: *F(3, 210)* = 0.521, *p* = 0.668, Paired: *F(3, 570)* = 5.46, *p* = 0.0010. *\*p* < 0.05, \**\*p* < 0.01 Tukey’s multiple comparison test, *n* = 15 mice, 191 neurons Paired group, *n* = 4 mice, 71 neurons Unpaired group). (**M**) Average ASt neuron sustained responses were greater in behavioral quartiles with more defensive behavior in paired mice, but not unpaired controls (Repeated measures ANOVA, Unpaired: *F(3, 210)* = 0.424, *p* = 0.736, Paired: *F(3, 570)* = 10.26, *p* < 0.0001. *\*p* < 0.05, \**\*p* < 0.01, \*\**\*p* < 0.001 Tukey’s multiple comparison test. *n* = 15 mice, 191 neurons Paired group, *n* = 4 mice, 71 neurons Unpaired group). (**N-U**) Analysis of ASt neuron responses to CS-reward across mice and trials grouped by latency of reward responses. (**N**) Two-tone discrimination task with electrophysiology recordings. Reward responses were calculated as the latency to reward port entry after tone onset. (**O**) Group average latency to port entry (reward response) during CS-reward was faster in paired group mice than unpaired group mice (*t17* = 2.803, *p* = 0.0122, Student’s t-test, *n =* 15 mice paired, 4 mice unpaired). **(P)** Hierarchical clustering of paired group mice into “low” and “high” learner/performer clusters based on reward responses. **(Q)** Group average reward responses were significantly faster in “high” performance mice than “low” performance and unpaired mice. (One-way ANOVA, F(2,16) = 28.59, *p* < 0.0001. \*\**\*p* < 0.001, Tukey’s multiple comparison test. *n =* 4 mice Unpaired, *n* = 4 mice “low” paired group, *n =* 11 mice “high” paired group). (**R**) Average ASt neuron phasic responses to CS-reward were not different in the “low” or “high” reward behavior groups (One-way ANOVA, F(2,16) = 2.178, *p* = 0.1457). (**S**) There was little correlation between ASt neuron phasic responses and mean reward latency. (**T**) Average latency to port entry during CS-reward for unpaired and paired groups, separated into quartiles determined based on relative latency of reward responses (Q1 = slowest response trials, Q4 = fastest response trials). (**U**) Average ASt neuron phasic responses were unchanged across different trial quartiles, regardless of changes in reward behavior (Repeated measures ANOVA, Paired: *F(3, 570)* = 0.576, *p* = 0.631). Bar graph and trace data shown as mean ±SEM.

**Figure S11.**
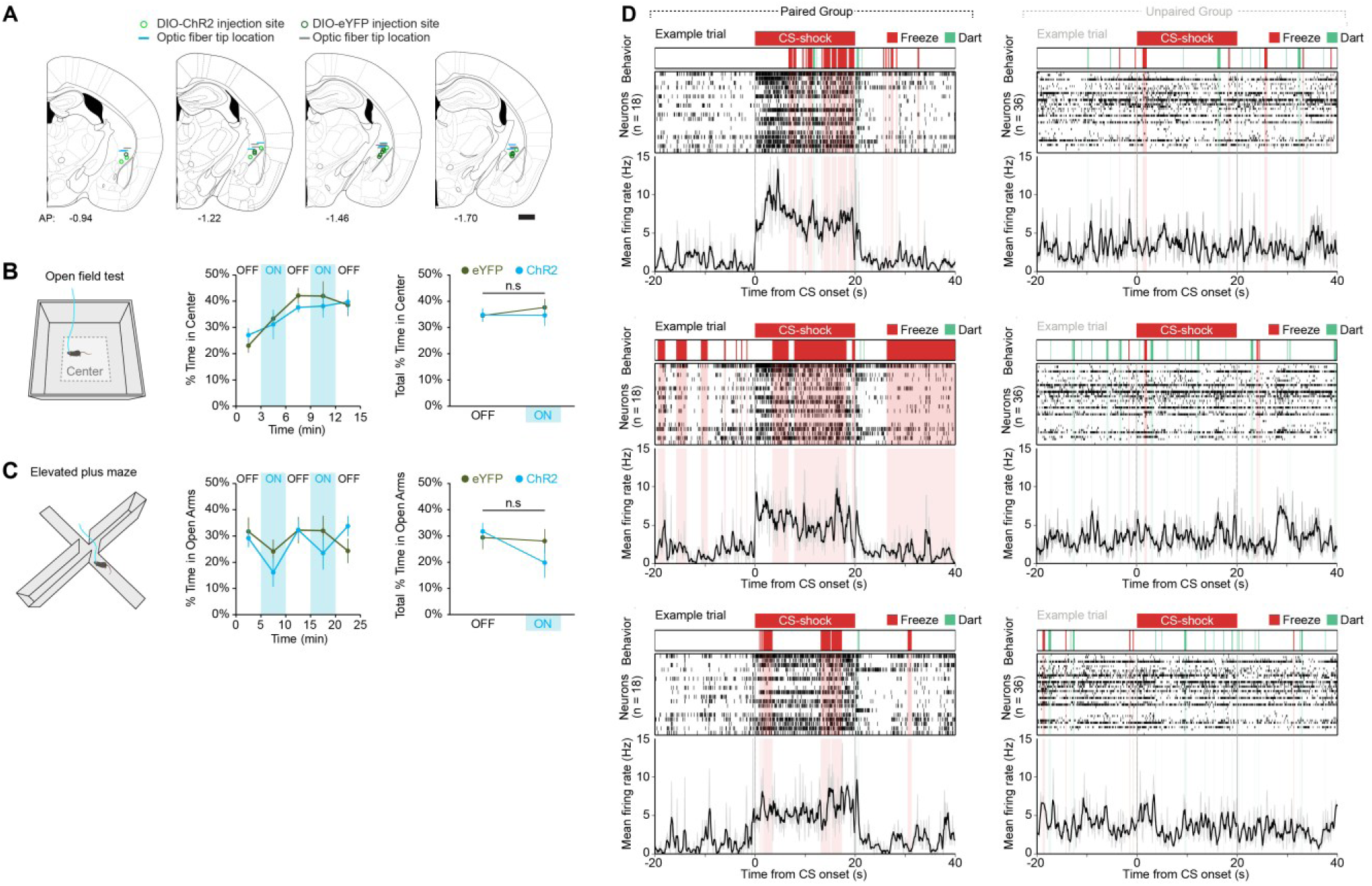
Optogenetic activation of ASt neurons does not affect anxiety-related behaviors. (**A**) Histologically verified optic fiber implant locations and viral injection sites for AAV-DIO-ChR2-eYFP and AAV-DIO-eYFP controls in optogenetic excitation experiments targeting ASt neurons in *VGAT*-Cre mice. Scale bars = 500 µm. (**B**) Optogenetic activation of ASt neurons had no effect on time spent in the center of an open field arena (Two-way ANOVA, group x laser interaction F(1,32) = 0.11, p = 0.74. ChR2: n = 8 mice, eYFP: n = 10 mice). (**C**) Optogenetic activation of ASt neurons had no effect on the proportion of time spent in the open arms of an elevated plus maze (Two-way ANOVA, group x laser interaction F(1,30) = 1.25, p = 0.27. ChR2: n = 7 mice, eYFP: n = 10 mice). (**D**) Additional representative trials from discrimination task experiments showing simultaneous subsecond behavior, neural spikes, and average neuron firing rate during presentation of CS-shock. In each example trial, neurons shown are all ASt neurons recorded from that mouse, during that trial.

**Figure S12.**
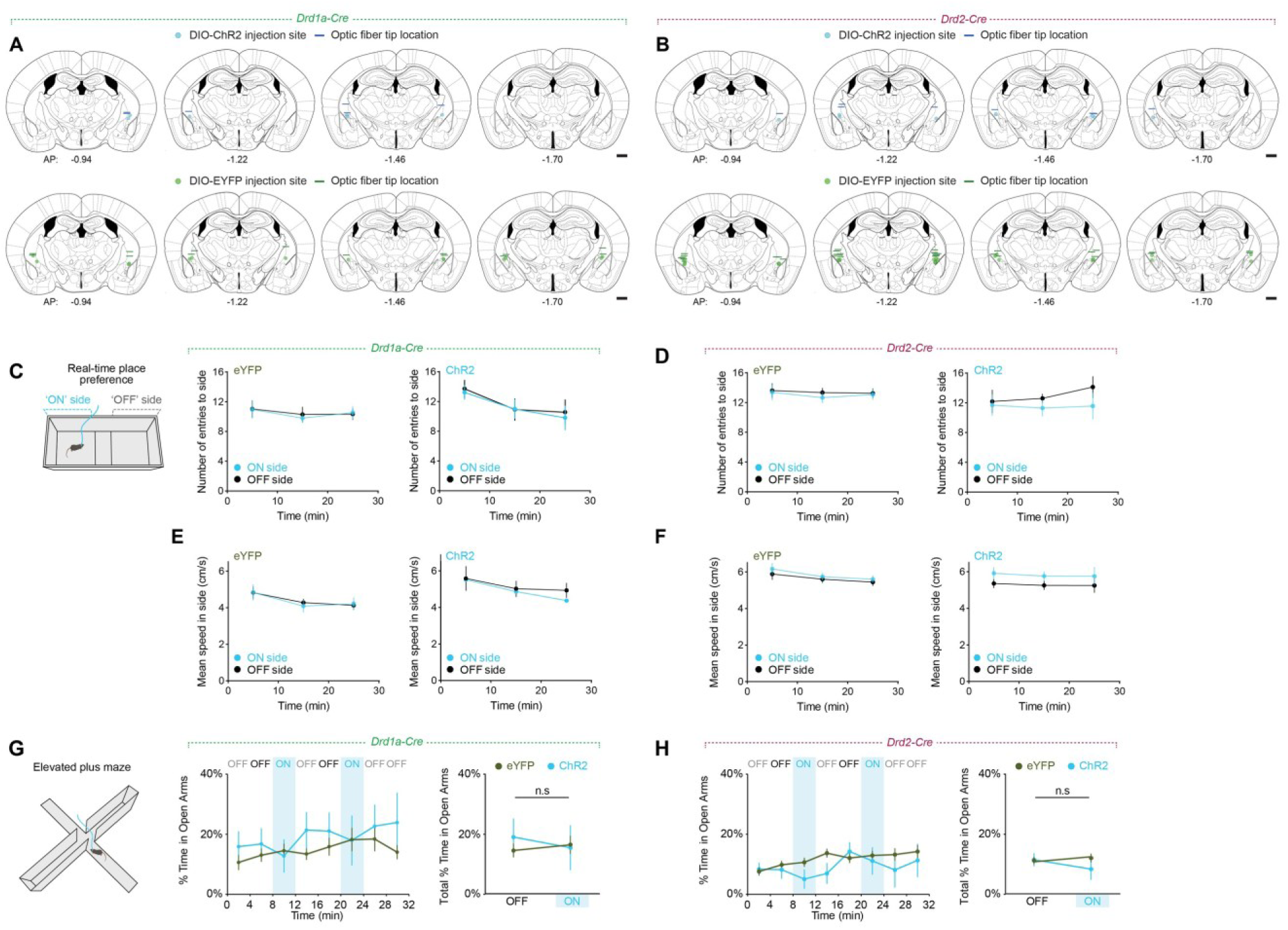
Optogenetic activation of *Drd1a+* or *Drd2+* ASt neurons does not affect side entries or overall locomotion during a real-time place preference task, or elevated plus maze behavior. **(A-B**) Histologically verified optic fiber implant locations and injection sites of AAV-DIO-ChR2-eYFP and AAV-DIO-eYFP in *Drd1a-cre* mice (**A**) and *Drd2-cre* mice (**B**). Scale bars = 500 µm. (**C-F**) In real-time place preference tasks, optogenetic stimulation of *Drd1a+* or *Drd2+* ASt neurons had no significant effect on the number of entries to the ‘ON’ side or ‘OFF’ side (Two-way RM ANOVA, Drd1a-Cre ChR2: effect of side F(1,8) = 0.1086, p = 0.7502; eYFP: effect of side F(1,12) = 0.009449, p = 0.9242. Drd1a-Cre ChR2: n = 5 mice, eYFP: n = 7 mice; Two-way RM ANOVA, Drd2a-Cre ChR2: effect of side F(1,10) = 0.6994, p = 0.4225; eYFP: effect of side F(1,30) = 0.1386, p = 0.7123. Drd2a-Cre ChR2: n = 6 mice, eYFP: n = 16 mice), or the mean speed in the ‘ON’ side or ‘OFF’ side (Two-way RM ANOVA, Drd1a-Cre ChR2: effect of side F(1,8) = 0.2172, p = 0.6536; eYFP: effect of side F(1,12) = 0.0008348, p = 0.9774. Drd1a-Cre ChR2: n = 5 mice, eYFP: n = 7 mice; Two-way RM ANOVA, Drd2a-Cre ChR2: effect of side F(1,10) = 1.951, p = 0.1927; eYFP: effect of side F(1,30) = 0.3750, p = 0.5449. Drd2a-Cre ChR2: n = 6 mice, eYFP: n = 16 mice). (**G-H**) Optogenetic stimulation of *Drd1a+* or *Drd2+* ASt neurons had no effect on time spent in open arms of an elevated plus maze. (Two-way ANOVA, Drd1a-Cre, group x laser interaction F(1,20) = 0.4057, p = 0.5314. *Drd1a*-Cre ChR2: n = 5 mice, eYFP: n = 7 mice; Two-way ANOVA, *Drd2a*-Cre, group x laser interaction F(1,40) = 0.9907, p = 0.3256. Drd2a-Cre ChR2: n = 6 mice, eYFP: n = 16 mice). Data presented as mean ±SEM.

**Figure S13.**
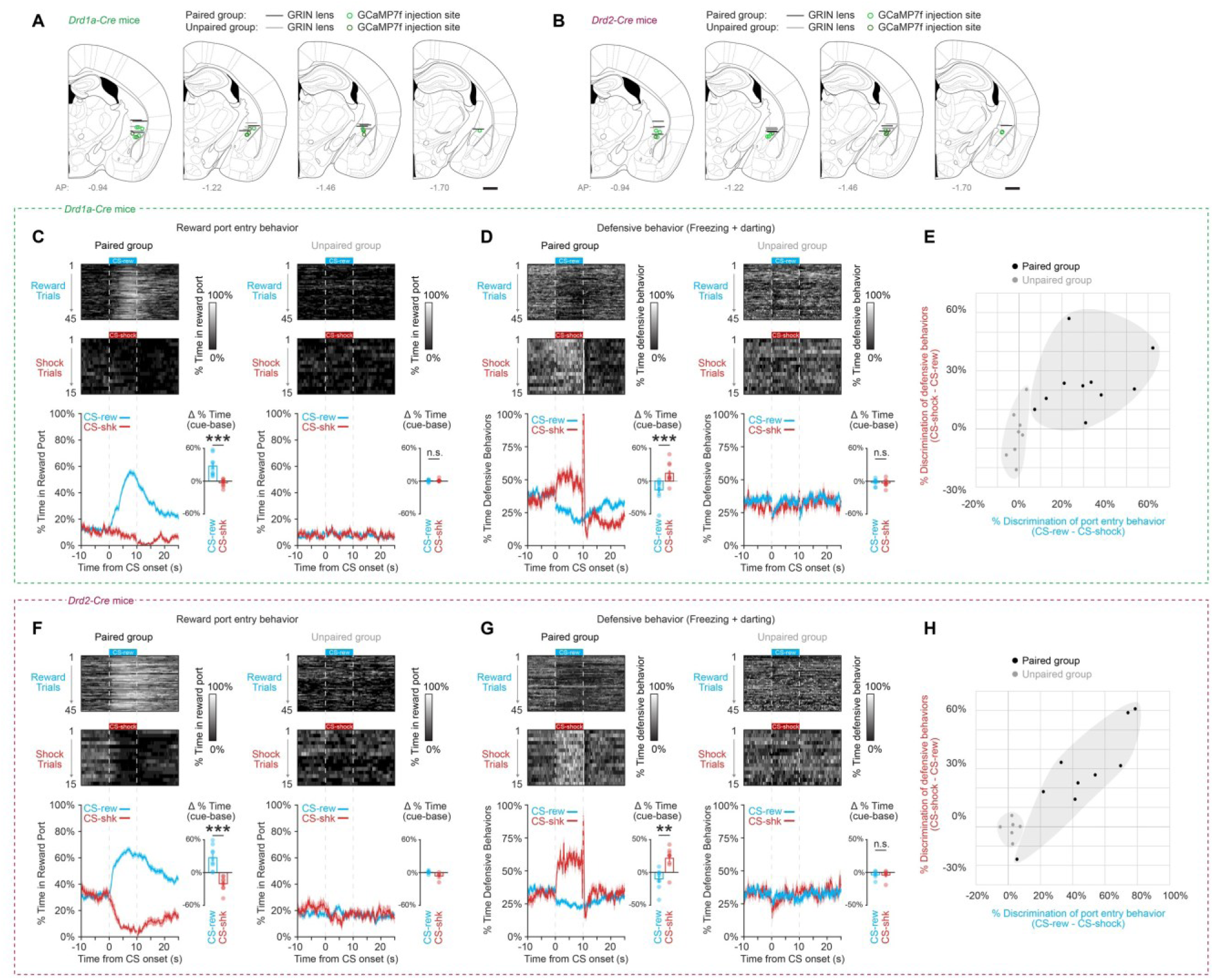
Targeting of ASt neurons and behavioral validation for *in vivo* calcium imaging. (**A-B**) Histologically verified GRIN lens implant locations and AAV1-Syn-FLEX-GCaMP7f viral injection sites in recording experiments targeting *Drd1a+* neurons (**A**) and *Drd2+* neurons (**B**) in the ASt. (**C-E**) Validation of conditioned behavioral responses during two-tone discrimination task in *Drd1a-Cre* mice (*Drd1a-Cre* Paired: *n =* 10 mice, *Drd1a-Cre* Unpaired: *n =* 8 mice). (**C**) Paired group mice showed distinct responses to the CS-shock and CS-reward tones in reward port entry behavior (Two-tailed paired t-test, *t9* = 6.400, ***p = 0.000125). (**D**) Paired group mice also showed distinct defensive responses to CS-shock and CS-reward tones (Two-tailed paired t-test, *t9* = 5.418, \*\**\*p* = 0.0004), consistent with successful discrimination between the two tones. Neither measure differed between cue types in unpaired mice (port entry, t7 = 0.373, p = 0.720; defensive responses, t7 = 0.300, p = 0.773). (**E**) ‘Discrimination scores’ of reward port entry behavior (CS-rew – CS-shock) plotted against defensive behavior (CS-shock − CS-rew) for each mouse. (**F-H**) Validation of conditioned behavioral responses during two-tone discrimination task in *Drd2-Cre* mice (*Drd2-Cre* Paired: *n =* 9 mice, *Drd2-Cre* Unpaired: *n =* 6 mice). Paired group mice showed distinct responses to the CS-shock and CS-reward tones in (**F**) reward port entry behavior (Two-tailed paired t-test, *t8* = 6.43, \*\**\*p* = 0.0002) and (**G**) defensive responses (Two-tailed paired t-test, *t8* = 3.381, \**\*p* = 0.0096), consistent with successful discrimination between the two tones. In unpaired mice, defensive responses did not differ between cue types (t5 = 0.110, p = 0.917); reward port entry differed slightly (t5 = 2.83, p = 0.0365), though the mean discrimination score was approximately 7-fold smaller in magnitude than in paired mice. (**H**) ‘Discrimination scores’ of reward port entry behavior (CS-rew − CS-shock) plotted against defensive behavior (CS-shock – CS-rew) for each mouse.

**Figure S14.**
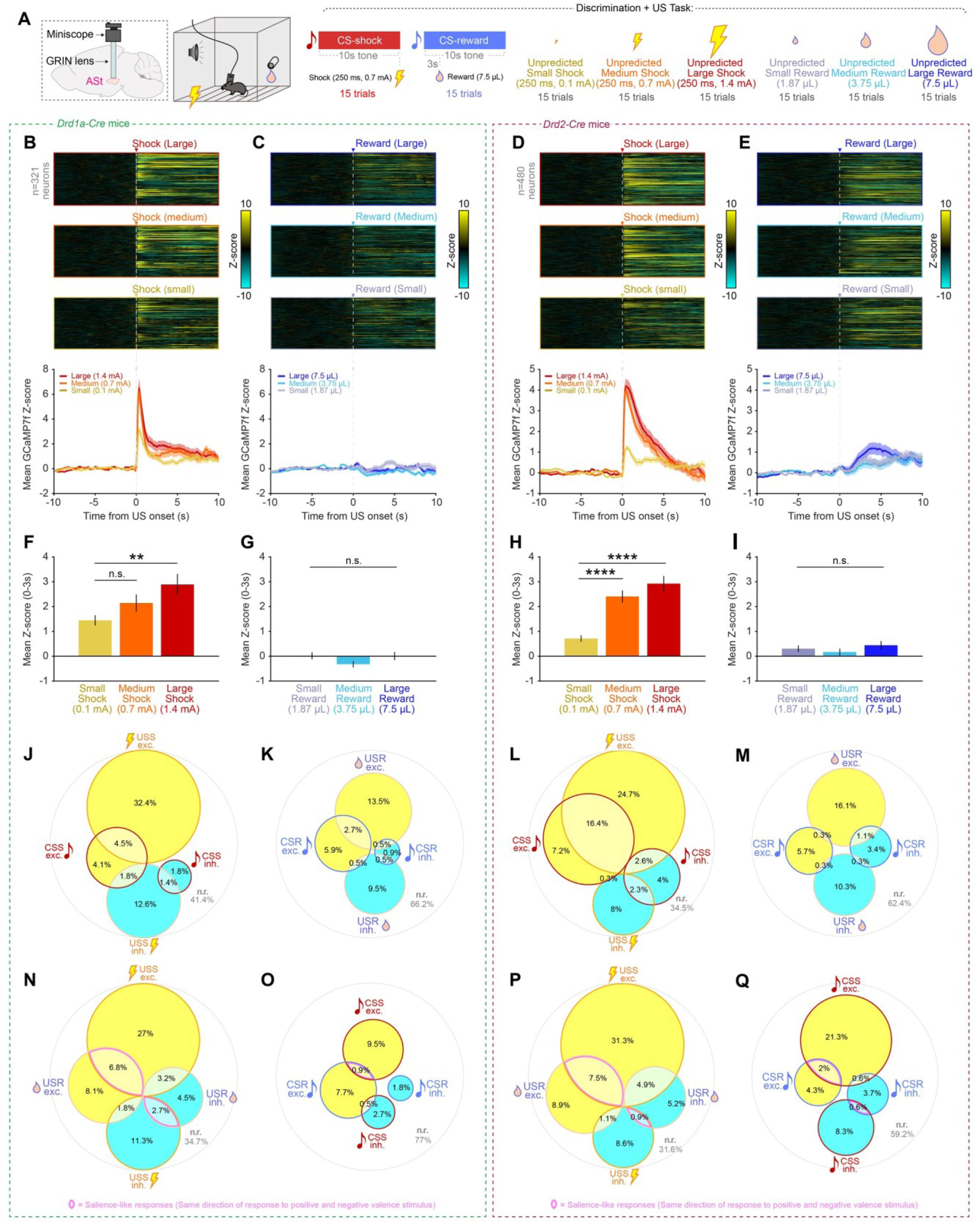
ASt neuron responses to unconditioned stimuli are heterogeneous and scale in magnitude for unpredicted shocks but not for unpredicted rewards. (**A**) Experimental task design for examining US responses in *Drd1a+* and *Drd2+* ASt neurons. In addition to normal CS-shock and CS-reward trials, mice were also presented with unpredicted deliveries of US-shock of different magnitudes (0.1, 0.7 or 1.4 mA), and US-reward of different magnitudes (1.87, 3.75 or 7.5 μL chocolate Ensure™). (**B-E**) Individual neuron traces (top) and group averages (bottom) of z-score changes in GCaMP7f fluorescence for (**B**) *Drd1a+* ASt neuron responses to US-shock, (**C**) *Drd1a+* ASt neuron responses to US-reward deliveries, (**D**) *Drd2+* ASt neuron responses to US-shock and (**E**) *Drd2+* ASt neuron responses to US-reward deliveries. (**F-I**) Quantification of mean z-score responses to US deliveries of different magnitudes (0-3s after US onset) for (**F**) *Drd1a+* ASt neuron responses to US-shock, (**G**) *Drd1a+* ASt neuron responses to US-reward, (**H**) *Drd2+* ASt neuron responses to US-shock, and (**I**) *Drd2+* ASt neuron responses to US-reward (One-way ANOVA: (**F**) F(2,960) = 4.53, p = 0.011; (**G**) F(2,960) = 1.77, p = 0.17; (**H**) F(2,1437) = 23.12, p = 1.3 × 10⁻ ¹⁰; (**I**) F(2,1437) = 0.86, p = 0.42; followed by post hoc t-tests with Bonferroni correction for 3 comparisons, **p < 0.01, ****p < 0.0001). (**J-Q**) Euler diagrams of *Drd1a+* ASt neuron co-responsiveness to (**J**) US-shock and CS-shock, and (**K**) US-reward and CS-reward. In all Euler diagrams, neurons were classified as ‘excited’ by the stimulus if their mean z-score was greater than 2.58 (which corresponds to a *p*-value of 0.01) in 0-3s post-stimulus onset window, and classified as ‘inhibited’ by the stimulus if their mean z-score was less than -2.58. (**L-M**) Euler diagrams of *Drd2+* ASt neuron co-responsiveness to (**L**) US-shock and CS-shock, and (**M**) US-reward and CS-reward. (**N-O**) Euler diagrams of *Drd1a+* ASt neuron co-responsiveness to (**N**) US-shock and US-reward, and (**O**) CS-shock and CS-reward. (**P-Q**) Euler diagrams of *Drd2+* ASt neuron co-responsiveness to (**P**) US-shock and US-reward, and (**Q**) CS-shock and CS-reward. Pink inner borders indicate ‘salience-like’ responses, where neurons respond in the same direction (excited or inhibited) to positive or negative valence stimuli. For data examining US responses (panels B-I), *Drd1a+*: *n =* 7 mice, 321 neurons, *Drd2+*: *n =* 8 mice, 480 neurons. For data examining CS and US co-responsiveness (panels J-Q), only data from paired group mice (where each CS had meaning) was used: *Drd1a+*: *n =* 4 mice, 222 neurons, *Drd2+*: *n =* 5 mice, 348 neurons.

**Figure S15.**
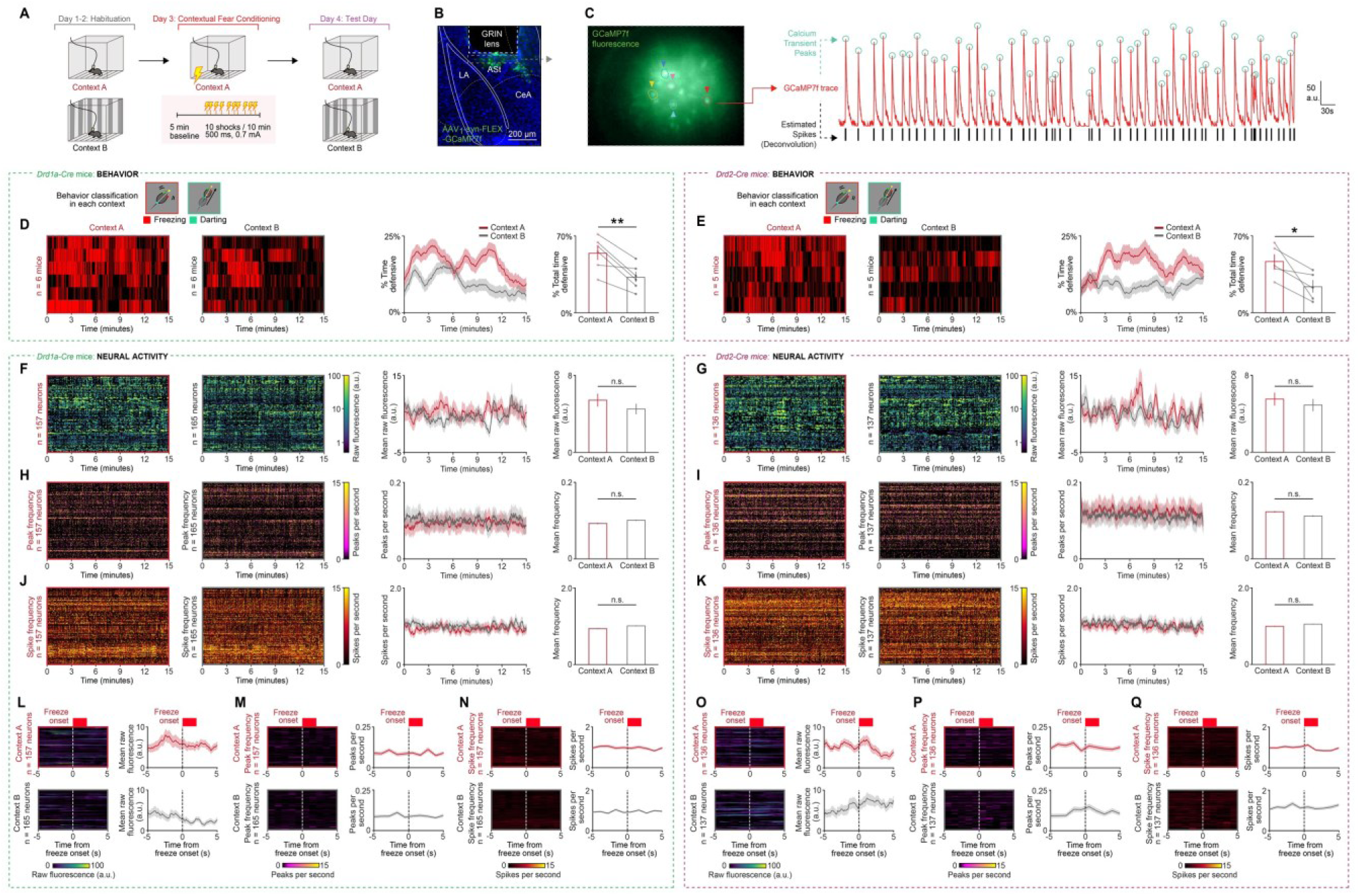
ASt neurons do not show changes in activity following contextual fear conditioning. (**A**) Contextual fear conditioning task experimental design. (**B**) GCaMP7f expression and lens targeting to ASt. (**C**) Calcium imaging field of view and representative traces of fluorescence, transient peaks detected and estimated spikes during contextual fear task. (**D-E**) Heatmaps and group averages of defensive behaviors during test day in Context A (conditioned) and Context B (unconditioned), in *Drd1a+* ASt neurons (**D**) and *Drd2+* ASt neurons (**E**). Mice in both groups spent significantly more total time freezing in Context A (conditioned) compared to Context B (unconditioned) (paired t-test, * p < 0.05, ** p < 0.01). (**F-G**) Heatmaps and group averages of raw neuron fluorescence in Context A and Context B on test day in *Drd1a+* ASt neurons (**F**) and *Drd2+* ASt neurons (**G**). (**H-I**) Heatmaps and group averages of frequency of calcium transient peaks in Context A and Context B on test day in *Drd1a+* ASt neurons (**H**) and *Drd2+* ASt neurons (**I**). (**J-K**) Heatmaps and group averages of frequency of estimated spike firing rates in Context A and Context B on test day in *Drd1a+* ASt neurons (**J**) and *Drd2+* ASt neurons (**K**). For F-K, paired t-test across mice all p > 0.05. (**L**) Heatmaps and group averages of *Drd1a+* ASt neuron raw fluorescence at the onset of freezing bouts in Context A and Context B. (**M**) Heatmaps and group averages of *Drd1a+* ASt calcium transient peaks at the onset of freezing bouts in Context A and Context B. (**N**) Heatmaps and group averages of *Drd1a+* ASt estimated spike firing rates at the onset of freezing bouts in Context A and Context B. (**O**) Heatmaps and group averages of *Drd2+* ASt neuron raw fluorescence at the onset of freezing bouts in Context A and Context B. (**P**) Heatmaps and group averages of *Drd2+* ASt calcium transient peaks at the onset of freezing bouts in Context A and Context B. (**Q**) Heatmaps and group averages of *Drd2+* ASt estimated spike firing rates at the onset of freezing bouts in Context A and Context B. No overall changes in neural activity to either context, or bouts of freezing while in either context, were observed in either *Drd1a+* or *Drd2+* ASt neurons. *Drd1a+* Context A: *n =* 6 mice, 157 neurons, *Drd1a+* Context B: *n =* 6 mice, 165 neurons, *Drd2+* Context A: *n =* 5 mice, 136 neurons, *Drd2+* Context B: *n =* 5 mice, 137 neurons.

**Figure S16.**
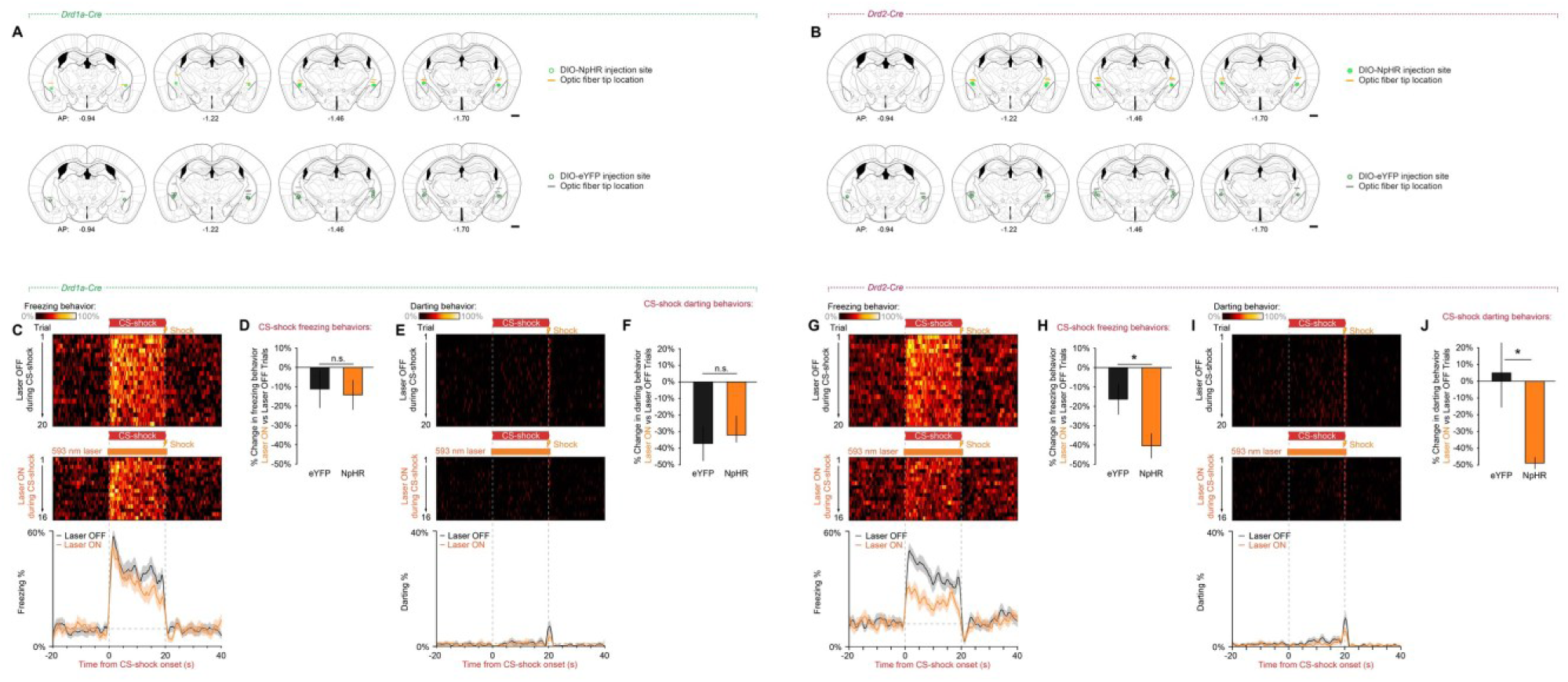
Targeting of *Drd1a+* and *Drd2+* ASt neurons and separate quantification of defensive behaviors during optogenetic inhibition. (**A-B**) Histologically verified optic fiber implant locations and injection sites of AAV-DIO-NpHR-eYFP and AAV-DIO-eYFP in optogenetic inhibition experiments in *Drd1a-Cre* and *Drd2-Cre* mice. Scale bars = 500 µm. (**C**) *Drd1a-Cre* freezing behavior heatmaps for CS-shock ‘laser OFF’ (top) and ‘laser ON’ trials (middle), and average freezing behavior during CS-shock trials with and without photoinhibition (bottom). (**D**) Inhibition of Drd*1a*+ ASt neurons had no effect on change in conditioned freezing behaviors to CS-shock in NpHR mice compared with eYFP controls (NpHR: −14.2 ± 7.7%; eYFP: −11.1 ± 9.8%; t13 = 0.223, p = 0.827, Student’s t-test; Drd1a+ NpHR: n = 6 mice, Drd1a+ eYFP: n = 9 mice). (**E**) *Drd1a-Cre* darting behavior heatmaps for CS-shock ‘laser OFF’ (top) and ‘laser ON’ trials (middle), and average darting behavior during CS-shock trials with and without photoinhibition (bottom). (**F**) Inhibition of Drd*1a*+ ASt neurons had no effect on change in conditioned darting behaviors to CS-shock in NpHR mice compared with eYFP controls (NpHR: −33.0 ± 8.1%; eYFP: −37.1 ± 10.5%; t13 = 0.279, p = 0.784, Student’s t-test; Drd1a+ NpHR: n = 6 mice, Drd1a+ eYFP: n = 9 mice). (**G**) *Drd2-Cre* freezing behavior heatmaps for CS-shock ‘laser OFF’ (top) and ‘laser ON’ trials (middle), and average freezing behavior during CS-shock trials with and without photoinhibition (bottom). (**H**) Inhibition of *Drd2+* ASt neurons resulted in a significantly greater reduction in conditioned freezing behaviors to CS-shock in NpHR mice compared with eYFP controls (NpHR: −40.4 ± 6.4%; eYFP: −16.3 ± 7.9%; t19 = 2.120, *p = 0.047, Student’s t-test; Drd2+ NpHR: n = 8 mice, Drd2+ eYFP: n = 13 mice). (**I**) *Drd2-Cre* darting behavior heatmaps for CS-shock ‘laser OFF’ (top) and ‘laser ON’ trials (middle), and average darting behavior during CS-shock trials with and without photoinhibition (bottom). (**J**) Inhibition of *Drd2+* ASt neurons resulted in a significantly greater reduction in conditioned darting behaviors to CS-shock in NpHR mice compared with eYFP controls (NpHR: −48.9 ± 3.5%; eYFP: 3.7 ± 19.4%; t19 = 2.097, *p = 0.0496, Student’s t-test; Drd2+ NpHR: n = 8 mice, Drd2+ eYFP: n = 13 mice). *p < 0.05. All data shown as mean ± SEM.

**Figure S17.**
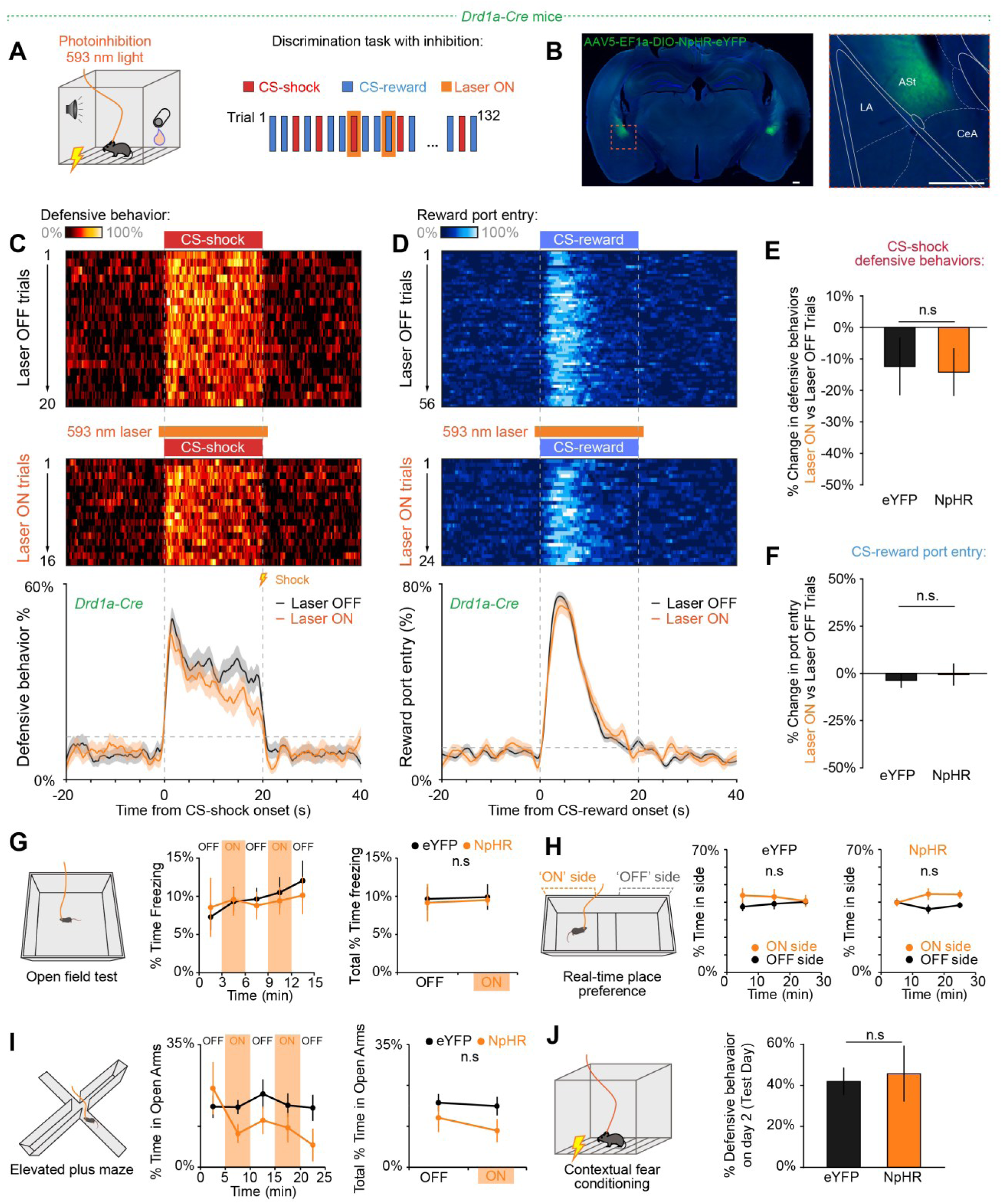
Optogenetic inhibition of D1+ ASt neurons does not affect behavioral responses to cues or context. (**A**) Behavioral paradigm for selective inhibition of *Drd1a+* ASt neurons during discrimination task. (**B**) Representative images of AAV-DIO-NpHR-eYFP expression in *Drd1a+* ASt neurons (Scale bars = 250 μm). (**C**) Defensive behavior heatmaps during ‘laser OFF’ (top) and ‘laser ON’ trials (middle) and group averages (bottom) during CS-shock trials with and without photoinhibition. (**D**) Port entry behavior heatmaps during laser OFF and ON (top, middle) and group averages (bottom) during CS-reward trials with and without photoinhibition. (**E**) Inhibition of Drd1a+ ASt neurons had no effect on conditioned defensive behaviors to CS-shock in NpHR mice compared with eYFP controls (t13 = 0.137, p = 0.893, Student’s t-test), and (**F**) had no effect on port entry during CS-reward (t13 = 0.493, p = 0.630, Student’s t-test). (**G**) Optogenetic inhibition of D1+ ASt neurons had no effect on overall locomotion in an open field task. (**H**) Inhibition of Drd1a+ ASt neurons did not cause preference or aversion for an area paired with inhibition in a real-time place preference task (Two-way RM ANOVA, NpHR: effect of side F(1,6) = 4.358, p = 0.0819; eYFP: effect of side F(1,16) = 0.6248, p = 0.4408). NpHR: n = 4 mice, eYFP: n = 9 mice. (**I**) Inhibition of *Drd1a+* ASt neurons caused no overall changes in exploration of open arms in an elevated plus maze. (**J**) Inhibition of *Drd1a+* ASt neurons had no effect on defensive behaviors on day 2 of a contextual fear conditioning task. *Drd1a+* NpHR: *n =* 6 mice, *Drd1a+* eYFP: *n =* 9 mice. All data shown as mean ±SEM.

